# Multicilia dynamically transduce Shh signaling to regulate choroid plexus functions

**DOI:** 10.1101/2025.01.21.633415

**Authors:** Suifang Mao, Rui Song, Shibo Jin, Song Pang, Aleksandra Jovanovic, Adam Zimmerman, Peng Li, Xinying Wu, Michael F. Wendland, Kerry Lin, Wei-Chi Chen, Semil P. Choksi, Gang Chen, Michael J. Holtzman, Jeremy F. Reiter, Ying Wan, Zhenyu Xuan, Yang K. Xiang, C Shan Xu, Srigokul Upadhyayula, Harald F. Hess, Lin He

## Abstract

Choroid plexus is a major site for cerebrospinal fluid (CSF) production, characterized by a multiciliated epithelial monolayer that regulates CSF production. We demonstrate that defective choroid plexus ciliogenesis or Intraflagellar transport yields neonatal hydrocephalus, at least in part, due to increased water channel Aqp1 and ion transporter Atp1a2 expression. We demonstrate choroid plexus multicilia as sensory cilia, transducing both canonical and non-canonical Shh signaling. Interestingly, it is the non-canonical Shh signaling that represses Aqp1 and Atp1a2 expression by Smo/Gαi/cAMP pathway. Choroid plexus multicilia exhibit unique ciliary ultrastructure, carrying features of both primary and motile cilia. Unlike most cilia that elongate during maturation, choroid plexus ciliary length decreases during development, causing a decline of Shh signaling intensity in developing choroid plexus, a derepression of Aqp1 and Atp1a2, and ultimately, an increased CSF production. Hence, developmental dynamics of choroid plexus multicilia dampens the Shh signaling intensity to promote CSF production.

## Introduction

The choroid plexus is a highly vascularized secretory neuroepithelium in brain ventricles, regulating the dynamic CSF production to support neuronal development and homeostasis ^1,2,3^. The choroid plexus consists of a multiciliated epithelial monolayer that envelops the underlying capillaries and connective tissue ^1,2^. Through the choroid plexus epithelium, directional transport of water, ions, glucose, proteins, and lipids is carried out from the choroidal blood supply into the brain ventricles ^4^. The choroid plexus expression of specific water channels, ion channels, and ion transporters is particularly important for CSF secretion ^5,6^. For example, deletion of *Aqp1*, the key choroid plexus water channel, significantly impairs CSF production in mice ^5^. CSF production is dynamically regulated during embryonic and perinatal development when an increase in CSF production coincides with the expansion of brain ventricles ^7,8,9^. An aberrant CSF production leads to a variety of developmental and physiological defects, including hydrocephalus, neurodevelopmental disorders, and neurodegeneration ^4,10,11^.

A hallmark of the choroid plexus is the monolayer of multiciliated cells that regulate CSF production. Cilia are microtubule-based cellular protrusions regulating diverse biological functions, including signaling transduction, fluid movement, cellular locomotion, and environmental sensing ^12–14^. In mice, mature choroid plexus multicilia are “9+0” non-motile multicilia ^15^, ultrastructurally distinct from primary cilia (a single “9+0” cilium that acts as solitary sensory hubs in a cell to transduce extracellular signaling ^16–19^) or motile multicilia (“9+2” multicilia, in dozens to hundreds per cell, which generate rhythmic beating for directional fluid movement ^20,21^). A previous study characterized choroid plexus cilia as a cilium type that contains features of both primary and motile cilia, which undergo axoneme regression during postnatal development ^22^. Yet the molecular functions and the developmental dynamics of choroid plexus cilia in the context of CSF production remain largely unknown.

Choroid plexus and ependymal epithelial cells in the mouse brain ventricles originate from the same neuroepithelial progenitors ^13^. Mature mouse choroid plexus cilia are predominantly “9+0” non-motile cilia, with transient motility observed at the neonatal stage ^15,23^; mature ependymal cilia are typical “9+2” motile cilia that beat in a synchronized manner to promote directional CSF flow ^24–27^. Both choroid plexus and ependymal epithelia express FoxJ1 ^24,28,29^, a key transcription factor for multiciliogenesis in airway epithelia ^30–31^, in ependymal epithelium ^24,28,29^, and zebrafish and xenopus choroid plexus ^32,33^. Using mouse genetics, we demonstrated FoxJ1 as an essential regulator for the ciliogenesis of the “9+0” choroid plexus multicilia in mice. Impaired choroid plexus cilia, either due to aberrant ciliogenesis or due to defective intraflagellar transport, contribute to the pathology of hydrocephalus, before the establishment of the CSF circulation directed by the beating ependymal cilia ^34–36^.

Here, our studies have demonstrated that choroid plexus “9+0” multicilia exhibit a distinct ultrastructural assembly, and function as sensory cilia that transduce both canonical and non-canonical Shh signaling. Our findings suggest that it is the non-canonical Shh signaling in the choroid plexus that represses the expression of Aqp1 (a water channel) and Atp1a2 (a subunit of Na^+^/K^+^-ATPase) in a cAMP-dependent mechanism, subsequently repressing CSF production in embryonic developmental stages. More importantly, the choroid plexus ciliary length is dynamically decreased during embryonic and perinatal development, causing the attenuation of the Shh signaling intensity, the derepression of Aqp1 and Atp1a2, and ultimately, the developmentally regulated increase in CSF production.

## Result

### FoxJ1 regulates choroid plexus ciliogenesis

*FoxJ1* is a key ciliogenesis transcriptional regulator for multi-motile cilia, mono-motile cilia, and “9+2” multi, non-motile olfactory cilia ^37–39^. *FoxJ1* starts to be expressed in mouse ependyma at E15.5-E18.5 with a peak induction at P4 to P6, and it governs ependymal multiciliogenesis in this critical developmental window ^24^. *FoxJ1* expression in choroid plexus appears earlier in development, initiating at E11.5 and peaking at E13.5 to E15.5 (Sup Fig. S1A) ^24,28,29^, implicating a possible role of FoxJ1 in choroid plexus ciliogenesis. *FoxJ1* deficiency caused a significant reduction of cilia number and ciliary length in choroid plexus epithelial cells (CPECs) (Fig. 1A, 1B, Sup Fig. S1B), with some *FoxJ1^-/-^* CPCEs exhibiting primary cilia features, such as a perpendicularly localized daughter centriole (Sup Fig. S1B). Similar defects were observed in *FoxJ1^-/-^* “9+2” motile cilia in ependymal and airway epithelia ^24,30,31^. Hence, our findings demonstrate FoxJ1 as an essential regulator for choroid plexus multiciliogenesis.

**Figure 1.**
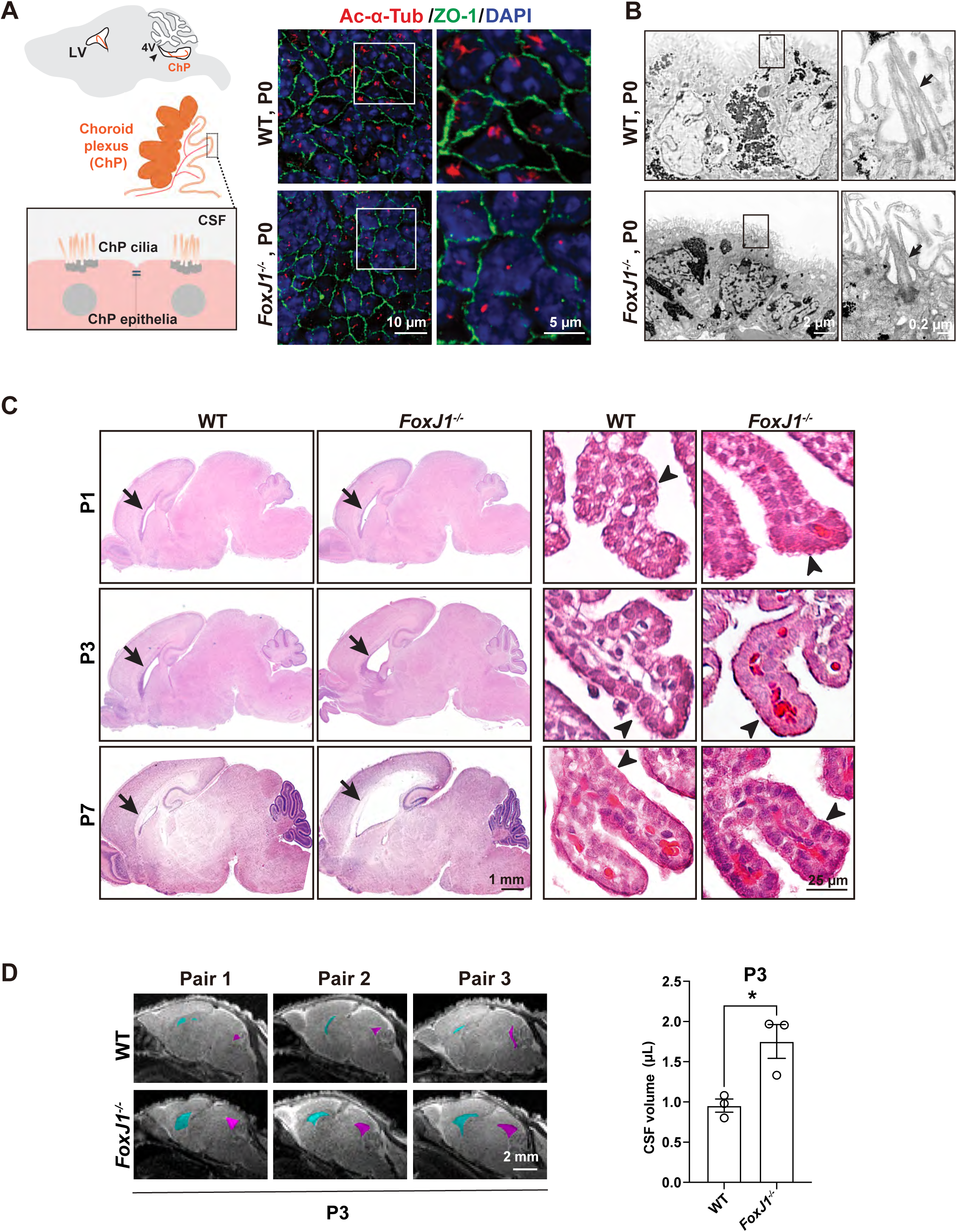
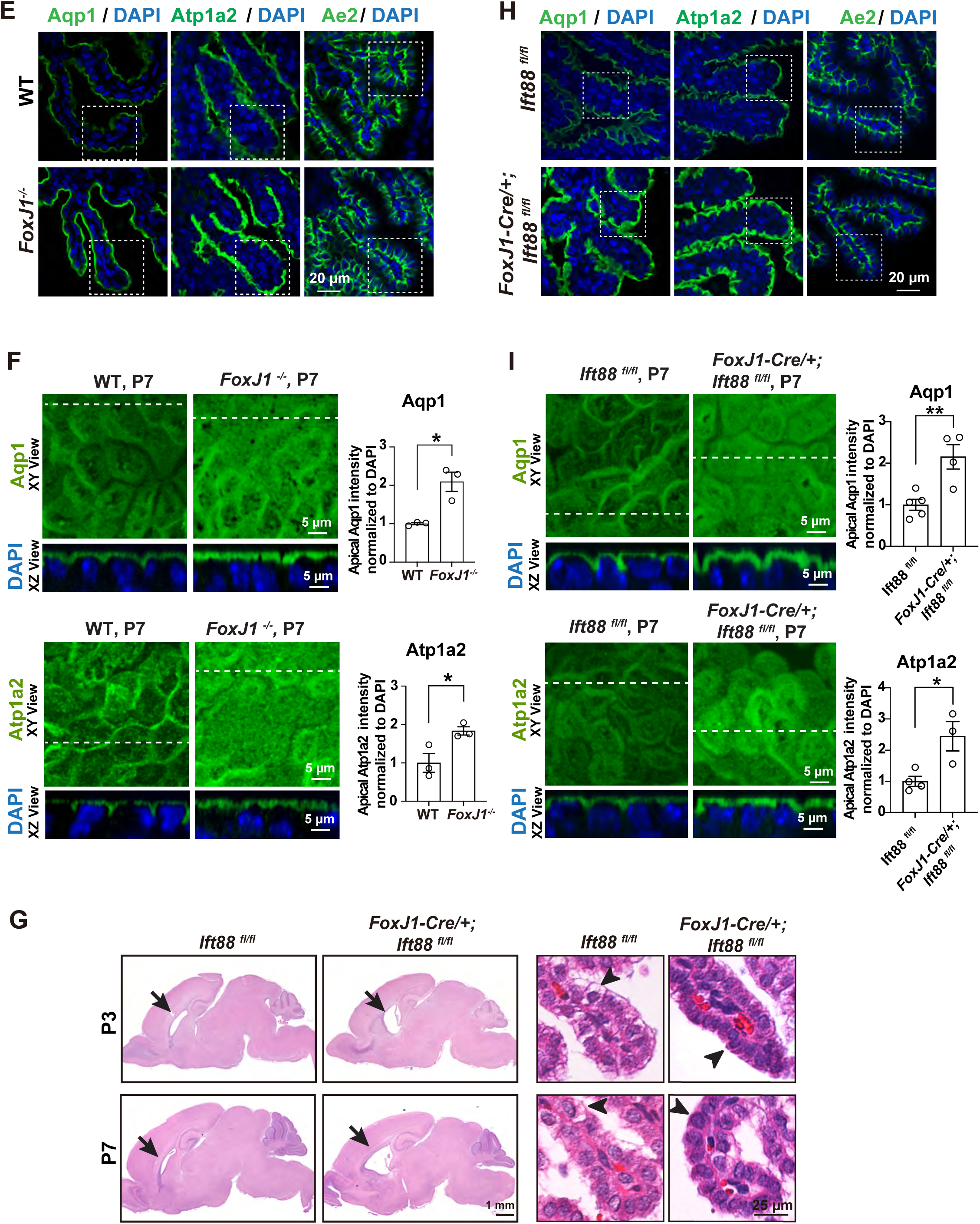
*Foxj1* deficiency impairs choroid plexus ciliogenesis and dysregulates CSF production. **A.** *FoxJ1* deficiency greatly decreases cilia number and ciliary length in the choroid plexus. **Left**, a schematic image showing the choroid plexus in the lateral ventricle and the 4^th^ ventricle, with a zoom-in image illustrating ciliated CPECs. **Right**, Representative immunostaining images of wildtype (WT) and *FoxJ1^-/-^* CPECs at P0. Ac-α-Tub (ciliary axonemes), ZO1 (tight junctions), and DAPI, n=3 littermate-controlled pairs. **B.** A TEM comparison between littermate-controlled P0 WT and *FoxJ1^-/-^*choroid plexus revealed a decreased cell volume, more compact cell morphology, shortened cilia, and a decreased cilia number in *FoxJ1^-/-^*CPECs. Arrows, cilia**. C.** *FoxJ1* deficiency causes enlarged lateral ventricles (arrows) and a decrease in intracellular vesicles in CPECs (arrowheads) in H&E analysis. Littermate controlled WT *and FoxJ1^-/-^* mice were compared at P1 (n=3), P3 (n=4) and P7 (n=6). **D.** Sagittal MRI scan images (left) and quantitation (right) demonstrate an increased CSF volume in lateral (cyan) and the 4th (magenta) ventricles of *FoxJ1^-/-^* mice. (n=3 littermate controlled paris at P3, \**P* = 0.0241, unpaired two-tailed Student’s t-test. **E-F.** Immunostaining on paraffin sections (E) and whole-mount tissue (F) showed an increased apical Aqp1 and Atp1a2 expression in P7 *FoxJ1^-/-^* CPECs. n= 3, littermate controlled WT and *FoxJ1^-/-^* mice; apical Aqp1, \**P* = 0.0199; apical Atp1a2, \**P* = 0.0286. **G.** In *FoxJ1-Cre/+; Ift88^fl/fl^* mice, *Ift88* deficiency in ciliated cells causes enlarged lateral ventricles (arrows) and a decrease in intracellular vesicles in CPECs (arrowheads) in H&E analyses. Littermate controlled *Ift88^fl/fl^ and FoxJ1-Cre/+; Ift88^fl/fl^* mice were compared at P3 (n=3) and P7 (n=3). **H-I.** Immunostaining on paraffin sections (H) and whole-mount tissue (I) showed an increased apical Aqp1 and Atp1a2 expression in P7 *FoxJ1-Cre/+; Ift88^fl/fl^* CPECs. Littermate controlled *Ift88^fl/fl^ and FoxJ1-Cre/+; Ift88^fl/fl^* mice were compared. Apical Aqp1, *Ift88^fl/fl^* (n=5) vs. *FoxJ1-Cre/+; Ift88^fl/fl^* (n=4), \*\**P* = 0.0061; apical Atp1a2, *Ift88^fl/fl^*(n=4) vs. *FoxJ1-Cre/+; Ift88^fl/fl^* (n=3), \**P* = 0.0215. All statistics are calculated with nested unpaired two-tailed Student t-test unless specified. All error bars are s.e.m. unless specified.

### Choroid plexus cilia defects impair CSF production and contribute to hydrocephalus

*FoxJ1* deficiency in mice leads to hydrocephalus (Fig. 1C), which was originally reported at P7 to P14 ^24,30,31^ (Fig. 1C, Sup Fig. S1C) and attributed to an obstructed CSF circulation due to motility defects of ependymal cilia ^24–27^. However, hydrocephalus in *FoxJ1*^-/-^ mice occurred much earlier in development, as enlarged brain ventricles and an increased CSF volume were evident from P1 to P3 (Fig. 1C-1D, Sup Fig. S1D), a developmental stage before ependymal cilia establish a coordinated beating for CSF circulation ^34,35,40^. In ependyma, motile cilia first emerged in immature ependymal cells from P2 to P6 as short, dispersed cilia with random basal body orientation ^36,41–43^. Uncoordinated cilia beating in ependymal, accompanied by a weak CSF flow, starts to emerge around P4-P5 ^34,35,40^; followed by a coordinated cilia beating and a strong, directional CSF flow that is fully established by P14-P15 ^34–36^. Hence, the pathogenesis of FoxJ1-dependent, neonatal hydrocephalus is unlikely to be caused solely by impaired ependymal cilia.

Since FoxJ1 regulates the ciliogenesis of the choroid plexus in addition to that of ependyma, defective choroid plexus cilia could also contribute to the pathogenesis of neonatal hydrocephalus in *FoxJ1*^-/-^ mice (Sup Fig. S1C, S1D). When a confluent *FoxJ1*^-/-^ CPEC monolayer was sandwiched by two culture chambers *in vitro*, a greater volume increase was observed in the apical chamber (Fig. S1E), while the tight cell-cell junction was intact (Fig. 1A). These findings implicated an increased secretory function in *FoxJ1*^-/-^ CPECs, consistent with their compacted cell morphology and reduced intracellular vesicles (Fig. 1C). A critical step of CSF secretion is the directional water and ion transport from the choroidal blood supply into the brain ventricles through specific water/ion channels/transporters across the choroid plexus epithelium ^10,44^. In P7 *FoxJ1*^-/-^ CPECs, we observed an increase in the apical and total expression of Aqp1 and Atp1a2 (Fig. 1E-1F, Sup Fig. S1F-S1G), both of which are key regulators for CSF production ^5,6,45^. In comparison, FoxJ1 deficiency did not impact the expression of other choroid plexus ion channels, including Ae2 (a Na^+^-independent Cl^-^/HCO_3_ ^-^ exchanger) and NKCC1 (a Na+/K+/Cl- cotransporter) (Sup Fig. S1F, S1H). Altogether, defective CEPC cilia caused by *FoxJ1* deficiency increased Aqp1 and Atp1a2 expression, consequently, giving rise to an increase in secretary function.

We next characterized a different cilia mutant, *FoxJ1-Cre/+; Ift88^fl/fl^*, to validate the effect of choroid plexus cilia on Aqp1 and Atp1a2 expression and neonatal hydrocephalus. *Ift88* deficiency in FoxJ1-expressing CPECs caused impaired intraflagellar transport, giving rise to reduced ciliary length and cilia number (Sup Fig. S1I). Similar to *FoxJ1*^-/-^ mice, *FoxJ1-Cre/+; Ift88^fl/fl^* mice exhibited neonatal hydrocephalus with an increased Aqp1 and Atp1a2 expression in CPECs; both defects precede the establishment of ependymal cilia motility or directional CSF circulation (Fig. 1G, Sup Fig. S1J, S1L). The same phenotype was observed in yet another mouse model with defective intraflagellar transport, *Tg737^orpk^* mice ^46^, highlighting the importance of choroid plexus cilia in neonatal hydrocephalus. Hence, neonatal and postnatal hydrocephalus caused by ciliopathies result from an aberrant CSF production caused by choroid plexus cilia defects, and an obstructed CSF flow caused by ependymal cilia defects.

### Choroid plexus cilia mediate a non-canonical Shh signaling to repress Aqp1 and Atp1a2 expression

Mature choroid plexus cilia resemble primary cilia in their “9+0” axoneme microtubule configuration 15^,23^. Primary cilia is an essential hub to transduce Shh signaling ^47–49^. In the canonical pathway, Shh triggers the ciliary exit of its receptor Patched ^50^, promotes Smoothened (Smo) translocation into the cilia, and activates a Gli-dependent transcriptional program ^49,51^.

Similarly, choroid plexus multicilia also transduce Shh signaling (Fig. 2A-2C, Sup Fig. S2A). A ciliary Smo translocation was not only observed in choroid plexus culture upon Shh treatment (Fig. 2A, Sup Fig. S2A) but also detected in E14.5 choroid plexus tissue with abundant Shh expression ^52^ (Fig. 2B, Sup Fig. S2C). More importantly, Smo activation triggers *Gli1* induction in the canonical Shh pathway (Fig. 2C, Sup Fig. S2B-S2C).

**Figure 2.**
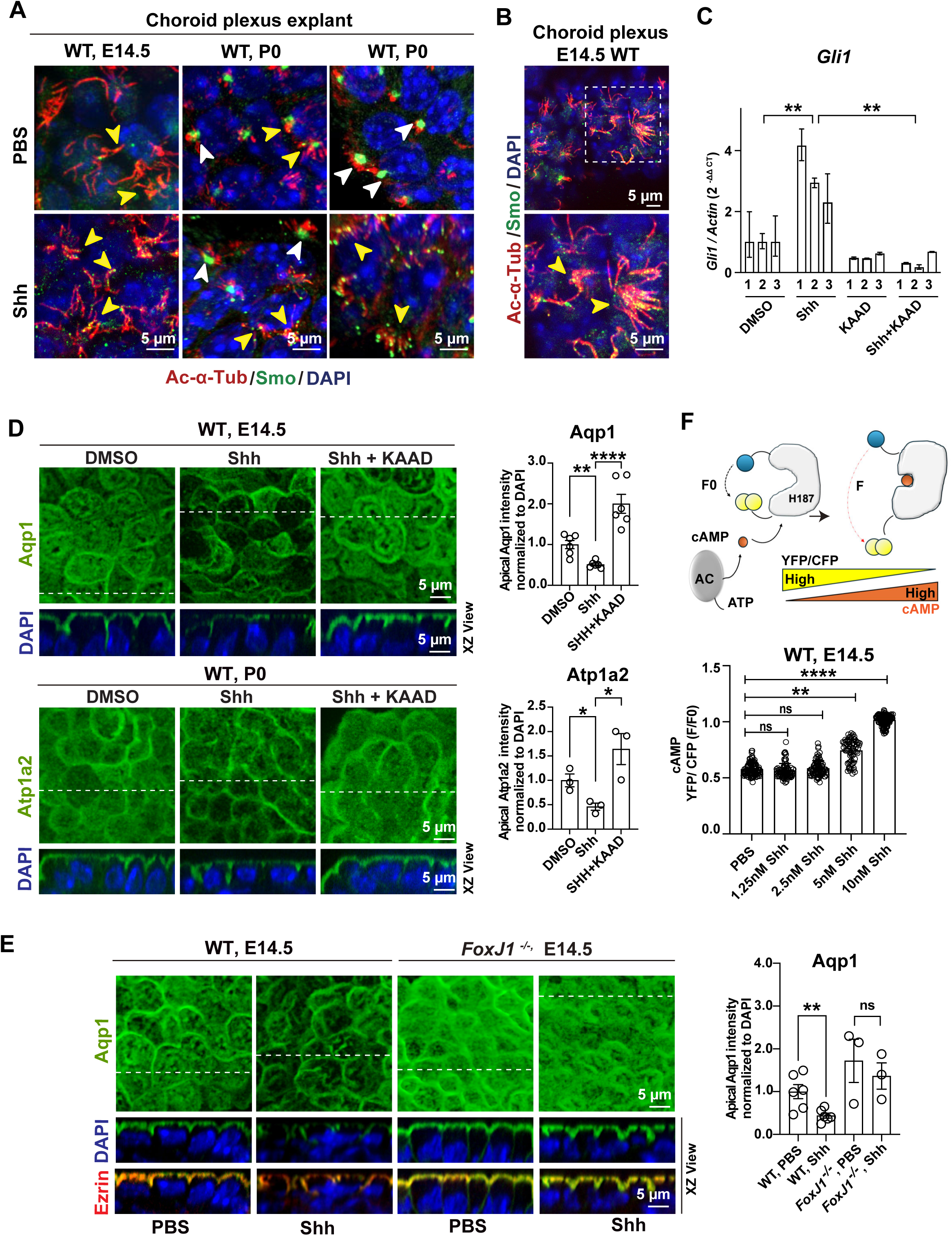
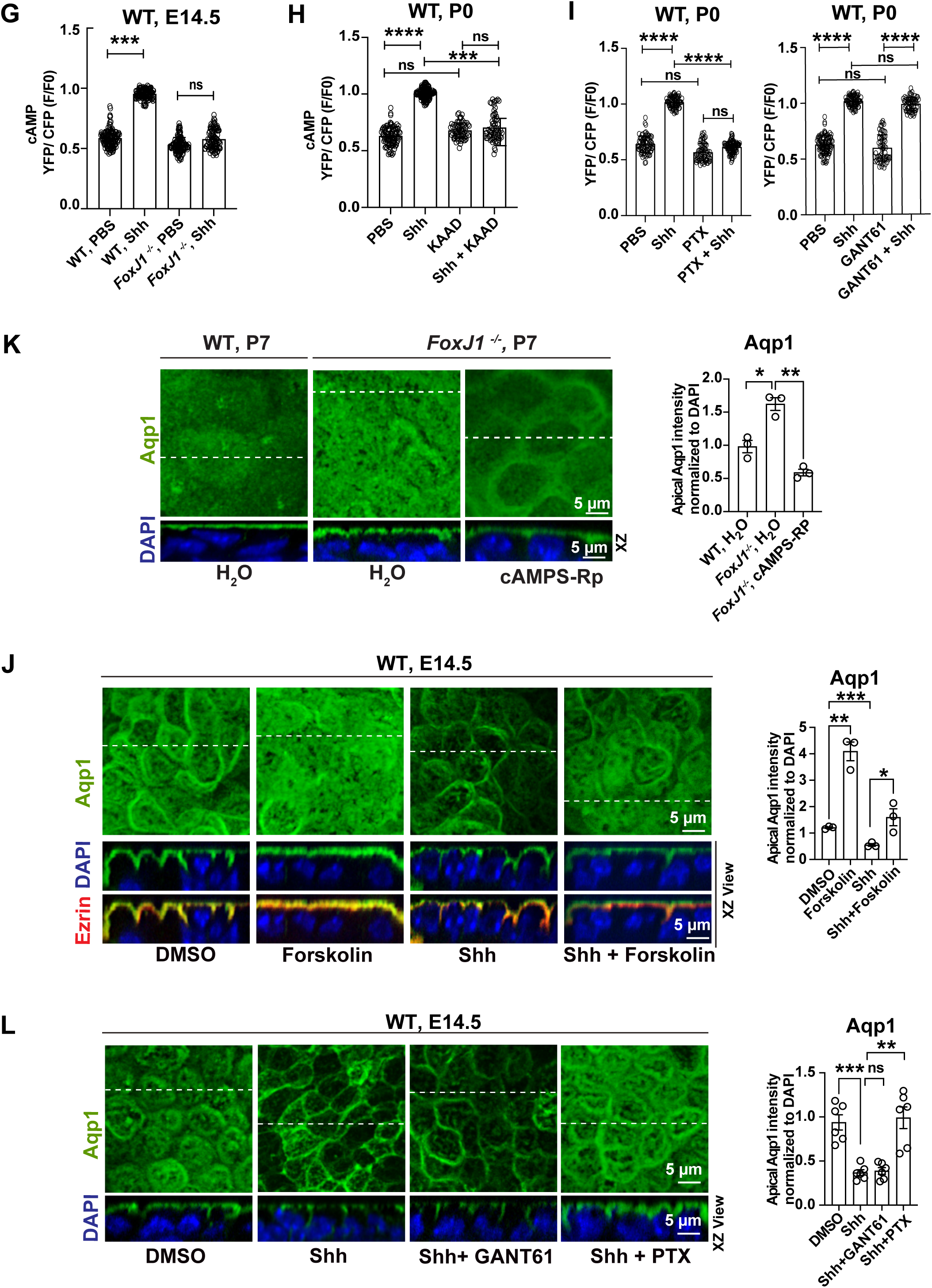
Choroid plexus cilia mediate Shh signaling to regulate Aqp1 and Atp1a2 expression. **A, B.** Immunostaining of Smo demonstrated its translocation into cilia in explant culture upon Shh treatment (E14.5 and P0) (**A**, n=3), or in whole mount tissue (E14.5) (**B**,n=3). **C**. In real-time PCR analysis, E14.5 WT Choroid plexus explant induces *Gli1* upon Shh treatment (10nM), and this induction was abolished by Smo inhibitor Cyclopmaine-KAAD (KAAD). DMSO vs. Shh, n=3, \*\**P* = 0.00178; Shh vs. Shh + KAAD, n=3, \*\**P* = 0.0085, error bars, s.d; unpaired two-tailed Student’s t-test. **D.** Immunostaining demonstrates an inhibitory effect of Shh signaling (10nM) on the apical expression of Aqp1 (top) and Atp1a2 (bottom) in WT choroid plexus explants, which was blunted by Smo inhibition using KAAD. Apical Aqp1 expression (n=6), DMSO vs. Shh, \*\**P* = 0.001; Shh vs. Shh + KAAD, \*\*\*\**P* < 0.0001. Apical Atp1a2 expression (n=3), DMSO vs. Shh, \**P* = 0.0497; Shh vs. Shh + KAAD, \**P* = 0.0257. **E.** Shh (10nM) downregulates apical Aqp1 expression in E14.5 wild-type choroid plexus explants, but not in littermate-controlled *FoxJ1^-/-^* samples. WT PBS vs. WT Shh (n=6), \*\**P* = 0.01; *FoxJ1^-/-^* PBS vs. *FoxJ1^-/-^* Shh (n=3), n.s.. **F-I.** Choroid plexus cilia mediate Shh signaling to reduce cAMP level. Error bars, s.d.. **F.** Top, a diagram illustrates the Epac-S^H187^ (exchange protein directly activated by cAMP) cAMP sensor, which decreases FRET upon cAMP binding. Bottom, E14.5 WT choroid plexus explants were treated with increasing concentrations of Shh, followed by ISO, and finally, with Forskolin (FSK, an activator of adenylyl cyclase) to achieve the maximal cAMP production. The dosage-dependent effect of Shh to reduce cAMP was measured by its inhibition on ISO-induced cAMP production. PBS (n=5) vs. 1.25nM Shh (n=3), n.s.; PBS (n=5) vs. 2.5nM Shh (n=3), n.s.; PBS (n=5) vs. 5nM Shh (n=4), \*\**P* = 0.0025; PBS (n=5) vs. 10nM Shh (n=5), \*\*\*\**P* < 0.00001; error bars, s.d.; nested one-way ANOVA with Sidak multiple comparison test. **G.** *FoxJ1* deficiency in choroid plexus abolished Shh inhibition on cAMP. WT PBS (n=5) vs. WT Shh (n=5), *** *P* = 0.0002; *FoxJ1^-/-^* PBS (n=3) vs. *FoxJ1^-/-^* Shh (n=4), n.s.; error bars, s.d.. **H, I.** Shh repression on cAMP depends on intact Smo and Gαi. In P0 WT choroid plexus explants, Shh (10nM) inhibition on cAMP was abolished by Smo inhibition (KAAD treatment, **H**) or by Gαi inhibition (PTX treatment), but not by Gli inhibition (GANT61 treatment). **H,** PBS vs. Shh (n=3), \*\*\*\**P* < 0.0001; Shh vs. Shh + KAAD (n=3), \*\*\**P* = 0.0009; error bars, s.d.. **I**. Left panel, PBS (n=4) vs. Shh (n=3), \*\*\*\**P* < 0.0001; Shh (n=3) vs. Shh + GANT61 (n=3), n.s.; GANT61 (n=3) vs. Shh + GANT61 (n=3), \*\*\*\**P* < 0.0001. Right panel, PBS (n=4) vs. Shh (n=3), \*\*\*\**P* < 0.0001; Shh (n=3) vs. Shh + PTX (n=3), \*\*\*\**P* < 0.0001; PTX (n=3) vs. Shh + PTX (n=3), n.s.; error bars, s.d. **J.** Immunostaining analyses showed that constitutive cAMP production due to FSK treatment impairs the inhibitory effect of Shh on the apical expression of Aqp1 in E14.5 WT choroid plexus,. DMSO vs. Forskolin (n=3), \*\**P* = 0.0012; DMSO vs. Shh (n=3), \*\*\**P* = 0.0004; Shh vs. Shh + Forskolin (n=3), \**P* = 0.032. **K.** Inhibition of cAMP-induced PKA activation (cAMPS-Rp treatment) downregulates the apical Aqp1 expression in P7 *FoxJ1*^-/-^ choroid plexus, as shown by immunostaining. WT H_2_O vs. *FoxJ1^-/-^* H_2_O (n=3), \**P* = 0.0254; *FoxJ1^-/-^* H_2_O vs. *FoxJ1^-/-^* cAMPS-Rp (n=3), \*\**P* = 0.0043. **L.** In immunostaining analyses, the inhibitory effect of Shh on apical Aqp1 expression in E14.5 WT choroid plexus was suppressed by Gαi inhibition (PTX treatment), but not by Gli inhibition (GANT61 treatment). DMSO vs. Shh (n=6), \*\*\**P* < 0.0001; Shh vs. Shh + GANT61 (n=6), n.s.; Shh vs. Shh + PTX (n=6), \*\**P* = 0.0004. All statistics are calculated with nested unpaired two-tailed Student t-test unless specified otherwise. All error bars are s.e.m. unless specified otherwise.

In addition to the induction of a Gli transcriptional program, Shh signaling in CPECs also yields a Smo-dependent decrease in apical and total expression of Aqp1 and Atp1a2 (Fig. 2D, Sup Fig. S2D-S2F), both at the mRNA level (Fig. S2G) and at the protein apical expression (Fig. 2D, Sup Fig. S2D). The inhibitory effect of Shh on Aqp1 and Atp1a2 depends on choroid plexus cilia, as *FoxJ1* deficiency abolished this effect (Fig. 2E, Sup Fig S2H). Unexpectedly, *FoxJ1^-/-^*choroid plexus cilia partially retained canonical Shh response, as measured by *Gli1* induction and Smo translocation (Sup Fig. S2I, S2J). Hence, choroid plexus cilia likely mediate Shh signaling to decrease Aqp1 and Atp1a2 expression in a Smo-dependent, but Gli-independent mechanism.

Shh signaling can be transduced via a non-canonical pathway that is independent of Gli ^53,54^. As the central transducer of the Shh pathway, Smo is a seven transmembrane domain protein, whose active structural conformation resembles that of an active Gαi-coupled G-protein-coupled receptor (GPCR) ^55–57^. In response to Shh, activated Smo can couple to Gαi to reduce cAMP production and inhibit PKA-dependent phosphorylation ^58^. Since impaired choroid plexus cilia elevated cAMP ^46,59^, we hypothesized that choroid plexus cilia transduce a non-canonical Shh signaling to downregulate cAMP. Using a choroid plexus explant culture, we boosted intracellular cAMP by isoproterenol (ISO), an agonist of the β-adrenergic receptor, and measured the inhibitory effect of Shh on ISO-induced cAMP using FRET (Fig. 2F) ^60^. Consistent with our hypothesis, Shh repressed intracellular cAMP in the choroid plexus in a dose-dependent manner (Fig. 2F), and this effect was abolished by defective cilia (Fig. 2G), Smo inhibition and Gαi inhibition (Fig. 2I), but not by Gli inhibition (Fig. 2I, Sup Fig. S2K).

We next tested if the non-canonical Shh signaling that reduced cAMP acts to decrease Aqp1 and Atp1a2 expression. Forskolin (FSK), an adenylyl cyclase activator, not only induced a strong apical Aqp1 expression in the choroid plexus but also blunted the inhibitory effect of Shh on Aqp1 (Fig. 2J). Conversely, cAMPS-Rp, a competitive antagonist of cAMP-induced PKA activation, reduced the aberrantly Aqp1 elevation in *FoxJ1^-/-^* choroid plexus (Fig. 2K). Finally, the inhibitory effect of Shh on Aqp1 expression was dampened by Gαi inhibition, but not Gli inhibition (Fig. 2L, Sup Fig. S2K). These findings support a model in which a non-canonical Shh signaling acts through choroid plexus cilia to downregulate apical Aqp1 expression via a cAMP-dependent mechanism.

### Choroid plexus cilia exhibit distinct ultrastructure

We employed TEM to characterize the ultrastructures of choroid plexus cilia (Fig. 3A, 3B), and compare them to those of primary cilia and motile cilia. While choroid plexus cilia resemble primary cilia with a “9+0” microtubule configuration and a lack of coordinated motility (Fig. 3B), they also resemble motile multicilia in multiplicity per cell and FoxJ1-dependent ciliogenesis (Fig. 1A, 3B, 3C, Sup Fig. S1B, S3A, and Table S1). All three cilia exhibit 9 radially arranged distal appendages (DAs) (Fig 3B, 3C). Yet choroid plexus cilia differ in their heterogeneous assembly of basal bodies that mostly contain 1, 2, 3, or 4 SDAs (Fig. 3C; Sup Fig. S3A). In contrast, primary cilia basal bodies contain 9 symmetrically distributed SDAs, and motile cilia basal bodies carry a single, uniformly oriented basal foot ^18,19^ (Fig. 3A, Sup Table S1). The “9+0” microtubule configuration, nine radially localized DAs, and basal bodies with heterogeneously organized SDAs are all conserved in mature human CPECs (Sup Fig. S3B). Interestingly, a subset of choroid plexus cilia has an electronic dense core, surrounded by 9 pairs of microtubule doublets (“9+c” configuration), which only appears in the proximal axoneme segment (Fig. 3C; Sup Fig. S3A). In a small subset of immature, neonatal choroid plexus cilia, a “9+2” and a “9+1” microtubule configuration are also reported, coinciding with a transient acquisition of uncoordinated motility ^23^.

**Figure 3.**
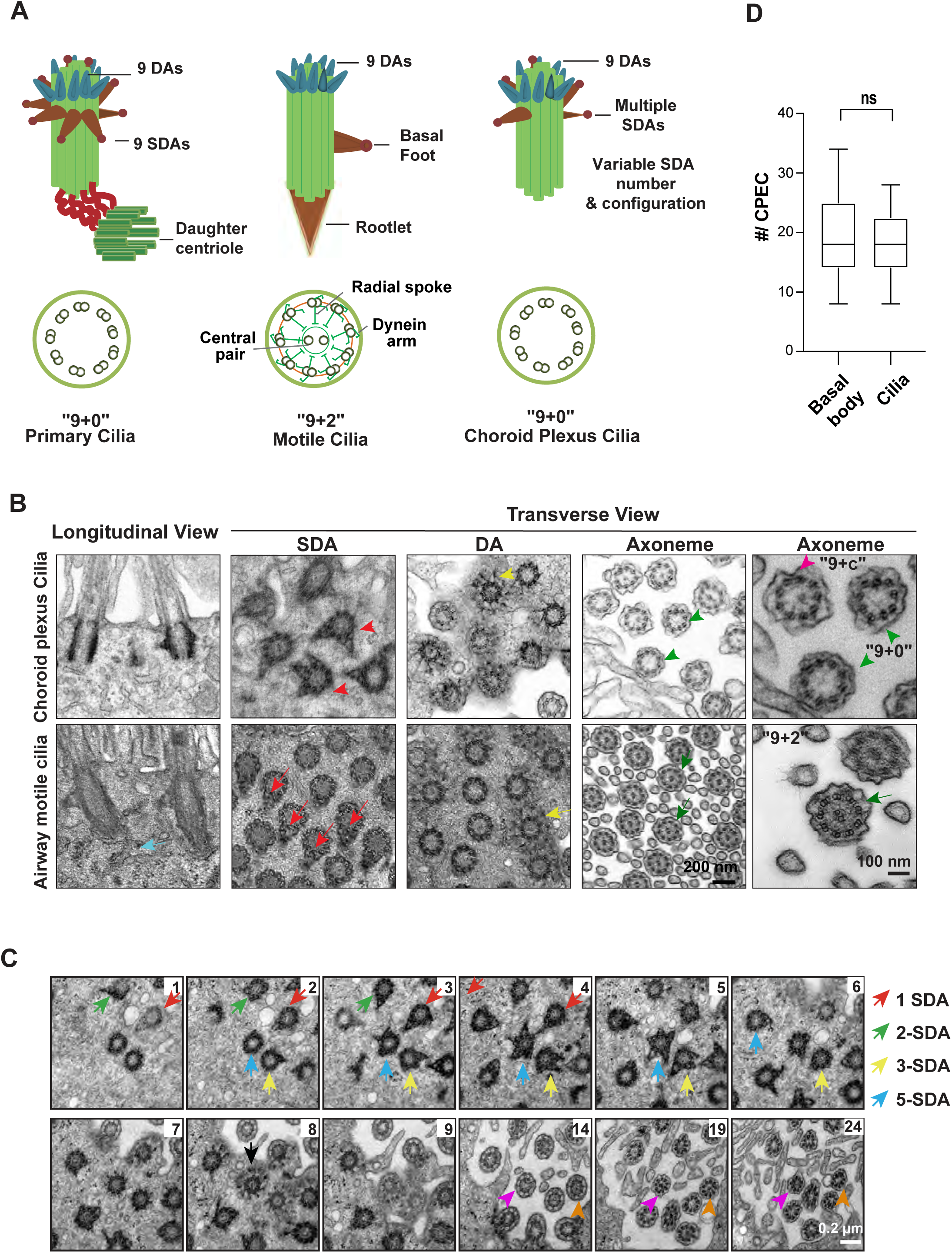
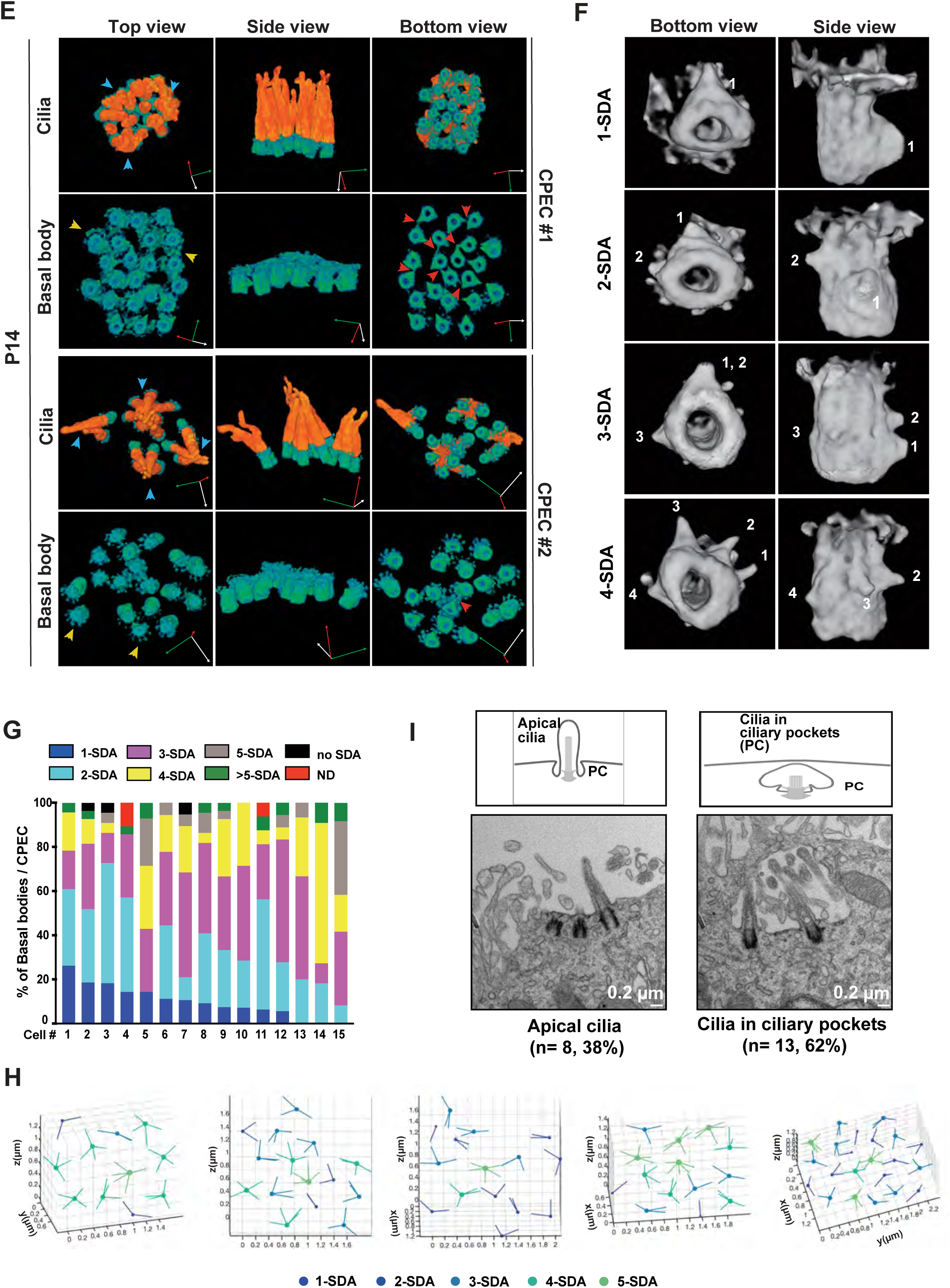
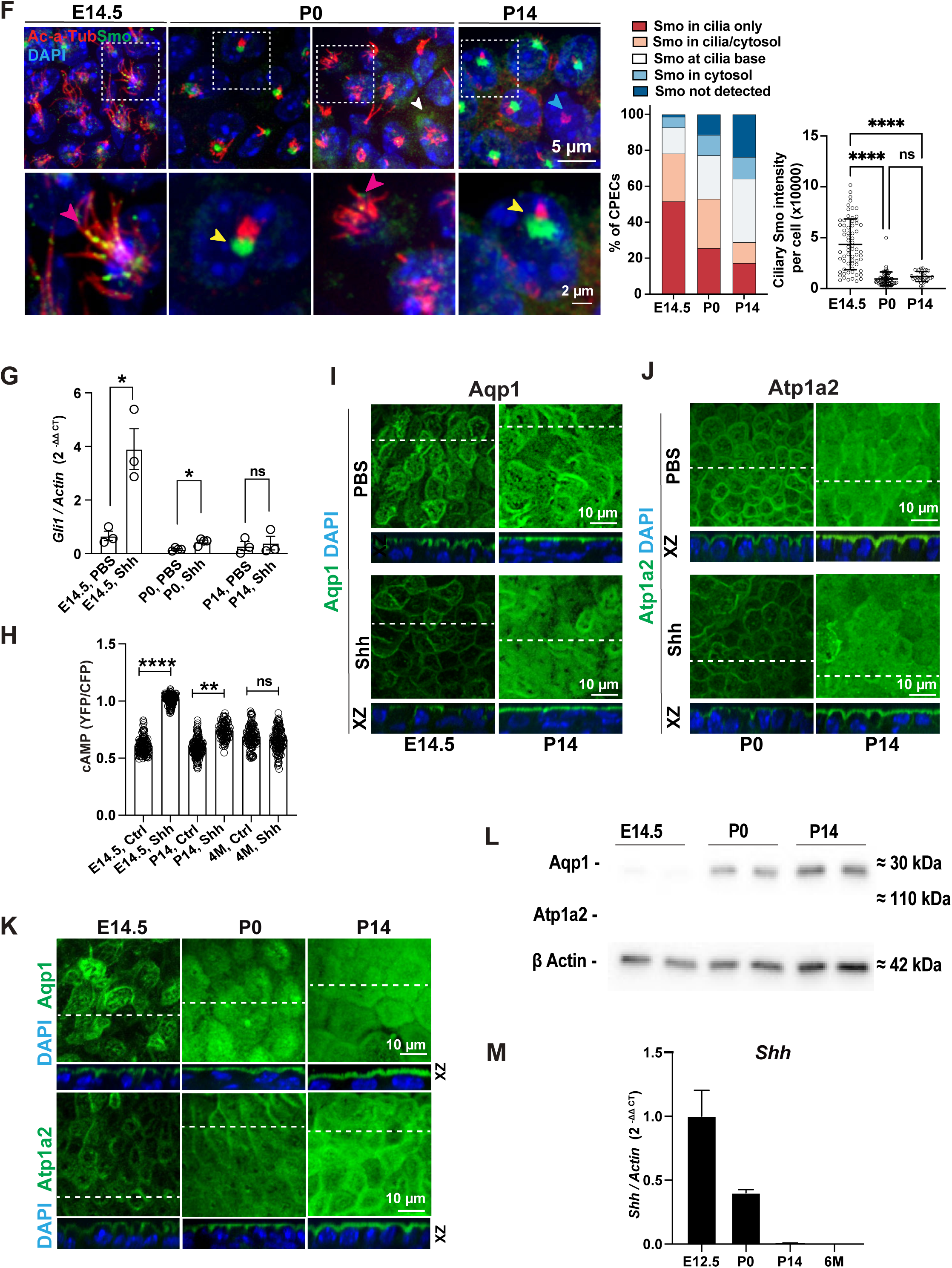
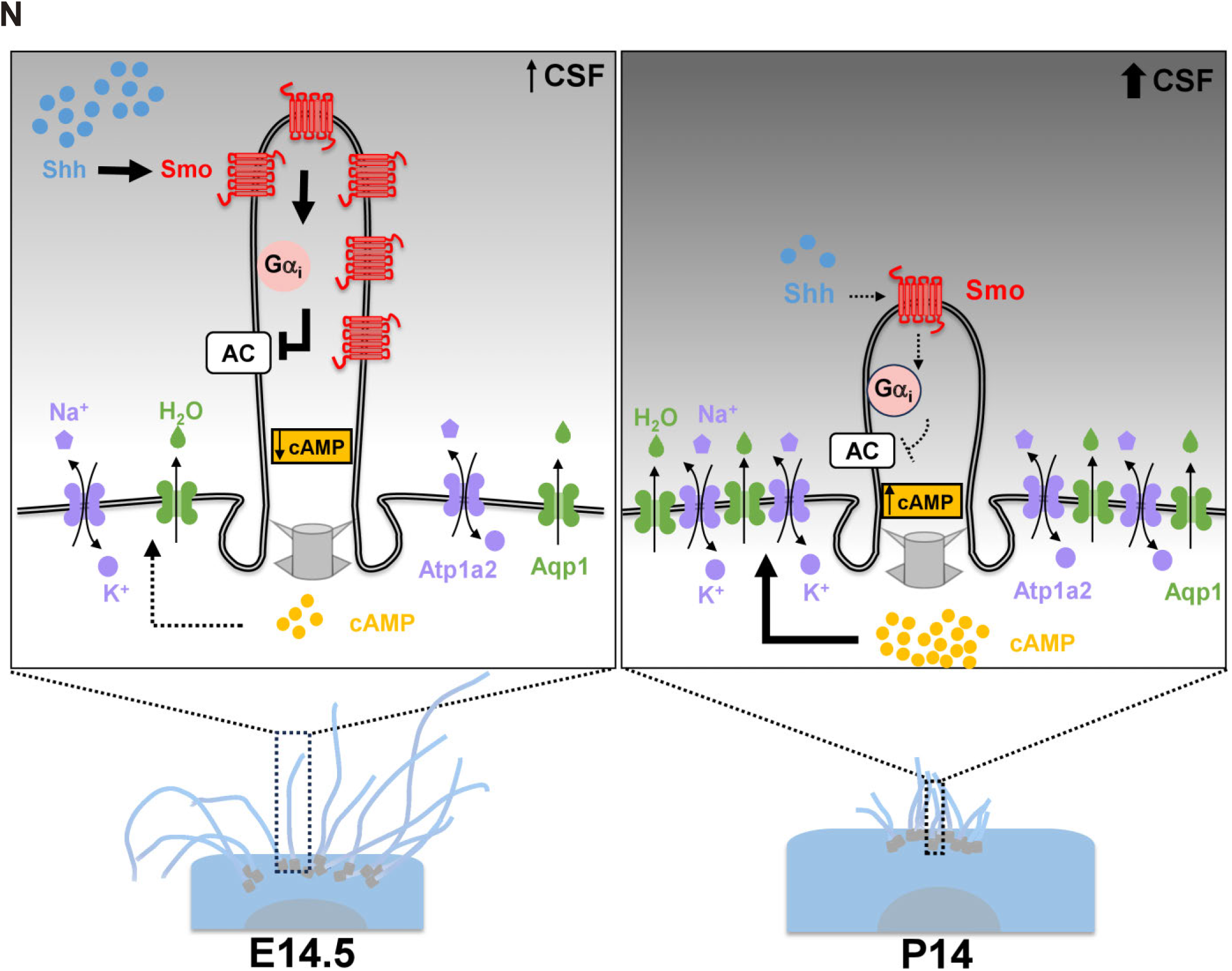
Mature choroid plexus multicilia exhibit distinct ciliary ultrastructure. **A.** A diagram summary illustrates an ultrastructural comparison among “9+0” primary cilia, “9+2” multi-motile cilia, and “9+0” choroid plexus multicilia. **B.** TEM analyses reveal an ultrastructural comparison between mouse airway motile cilia and choroid plexus multicilia at P14. Blue arrow, rootlet; green arrowhead, green arrow, and pink arrowhead, “9+0”, “9+2” and “9+c” microtubule configuration, respectively; red arrow, basal feet; red arrowhead, SDA; yellow arrowhead and arrow, DA. **C.** Selected serial TEM sections (50 nm/section, serial number indicated) from a 24-section series (Sup Fig S1A) revealed a distinct basal body assembly along the proximal to distal (PD) axis. Colored arrows, the same basal body/cilia along the PD axis with 1 (red), 2 (green), 3 (yellow), or 5 (blue) SDAs. Black arrow, DA. Some ciliary axonemes harbor an electronic dense core in the ciliary segment proximal to the basal bodies (pink and orange arrowheads). **D.** FIB-SEM quantification indicates an average of ∼18 cilia and basal bodies per CPEC (n=17). Cilia number per cell vs. basal body number per cell, n.s., unpaired two-tailed Student’s t-test; error bars, s.d. **E.** FIB-SEM 3D reconstructions of two representative mouse P14 CPECs, with one forming a single multicilia cluster with closely aggregated basal bodies (CEPC #1), and the other forming multiple subclusters of multicilia with more dispersed basal bodies (CEPC #2). Orange, axoneme; blue-green, basal body; blue arrowheads, aggregated tip of clustered cilia; yellow arrowheads, DAs; red arrowheads, basal body connections via SDA-SDA or SDA-basal body contact. 3D axes, X (red), Y (green), and Z (white). Bounding box dimensions: CPEC#1, top, 3.224 µm x 2.824 µm x 2.904 µm; bottom, 1.856 µm x 2.56 µm x 1.664 µm. CPEC#2, top, 3.144 µm x 4.456 µm x 2.776 µm; bottom, 1.288 µm x 1.328 µm x 1.024 µm. **F, G.** FIB-SEM 3D reconstruction images (**F**) and quantitations (286 basal bodies from 15 P14 CPECs, **G**) reveal the morphology of basal bodies with heterogeneous SDA configurations. **H.** A FIB-SEM analysis reveals a unique basal body organization, in which 4 or 5-SDA basal bodies are localized at the center of a CPEC, while 2 and 3-SDA basal bodies reside in the periphery. **I.** TEM images (left) and diagrams (right) show that choroid plexus multicilia are either localized at the apical plasma membrane (38%) or within a ciliary pocket (62%) (n=21).

Using Focused Ion Beam Scanning Electron Microscopy (FIB-SEM) analyses ^61,62^, we further analyzed 17 P14 CPECs. We obtained 286 imaged basal bodies (Sup Fig. S3C) and modeled the CPEC ciliary and basal body ultrastructure in 3D (Sup Movie S1). P14 CPECs exhibited heterogeneity in the numbers of cilia, with an average of ∼18 cilia and ∼18 basal bodies per cell (Fig.3D). Choroid plexus basal bodies have no ciliary attachment (Fig. 3E, 3F), and contain varying numbers of asymmetrically arranged SDAs (Sup Fig. S3D, Fig. 3G), often with one dominant SDA larger in size (Sup Fig. S3E). Multiple basal bodies within each CPEC form a clustered bundle, characterized by SDA-SDA or SDA-basal body contact (Fig. 3E). Basal bodies with more SDAs (4SDA or 5SDA) were often localized in the center of the basal body network, while those with fewer SDAs (2SDAs or 3SDAs) were frequently in the periphery (Fig. 3H). Such a heterogeneous basal body assembly likely stabilizes a cytoskeleton network that facilitates the clustering of multicilia, either within ciliary pockets or at the apical membrane (Fig. 3I). Choroid plexus cilia are previously characterized as nodal-like “9+0” cilia in a TEM study ^22^, yet our FIB-SEM data comprehensively and accurately depict their distinct ultrastructure, highlighting their unique basal body network as a structural basis for their biological functions.

### A decrease in choroid plexus ciliary length in development attenuates Shh signaling

The differentiated, multiciliated CPECs emerge around E11 and become mature by P14 ^23,63^. Unexpectedly, CPECs exhibit a decreased ciliary length during maturation (Fig. 4A-4C), with prominent, long, and scattered cilia (E12.5-E14.5) maturing into short and clustered cilia buried beneath the microvilli (P14-2 month) (Fig. 4A, 4C). This differs from most motile multicilia that elongate during maturation (Fig. 4D). During choroid plexus maturation, the ciliary length is reduced, the cilia and basal body numbers remain unaltered (Fig. S4A), and the cell size increases (Fig. S4B). Since an increased cell size of CPEC was previously shown to correlate with an increased secretory capacity ^64^, the mature CPECs are characterized by short, clustered cilia with a strong secretary function. Our studies differ from a previous one that focused on postnatal choroid plexus cilia regression ^22^. In the 4th ventricle choroid plexus, the most significant ciliary length reduction occurs between E12.5 and P14 (Fig 4A-4C), coinciding with functional maturation of choroid plexus cilia and CPECs, CSF volumetric growth, and water/ion channel and transporter induction ^65,66^.

**Figure 4.**
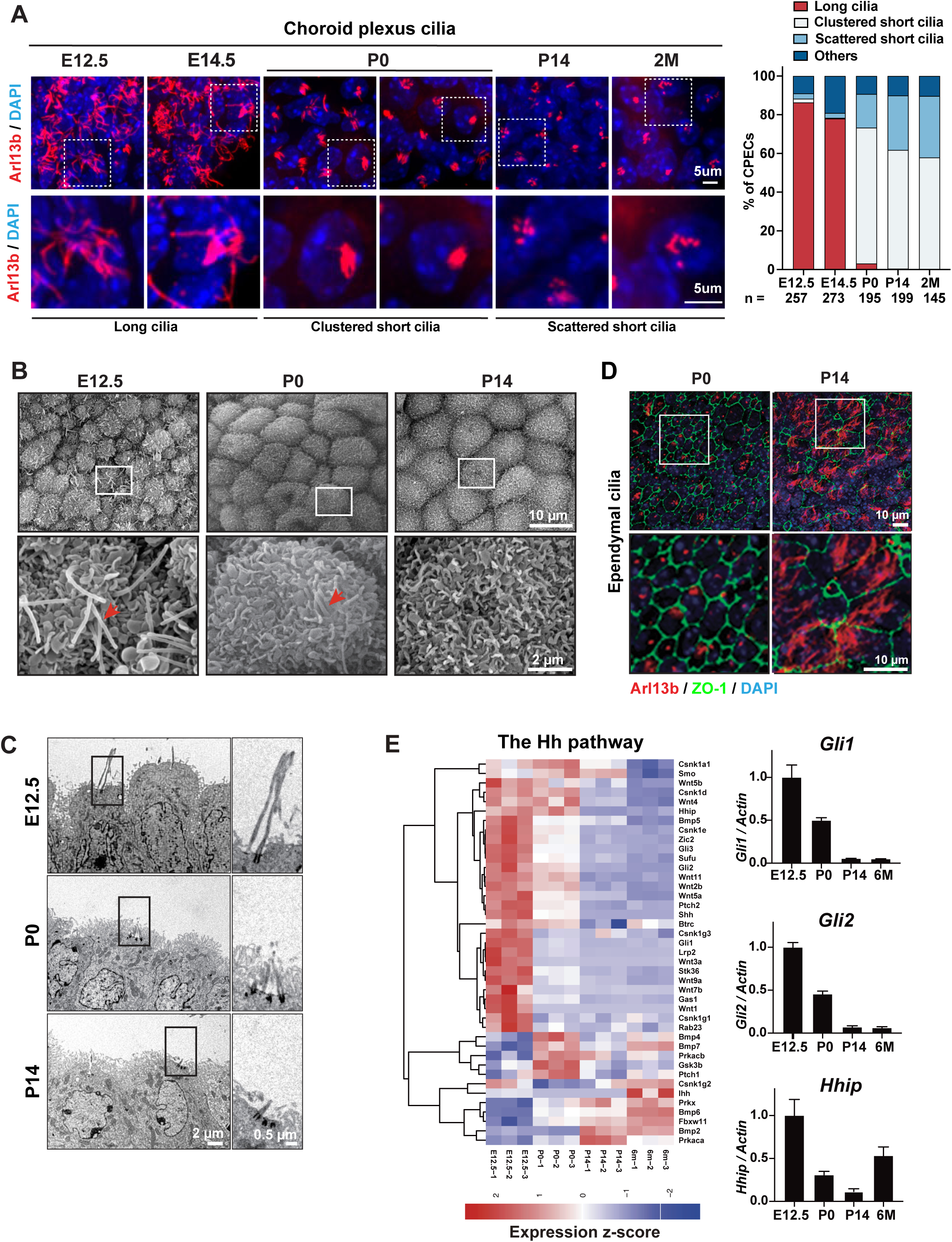
A decrease in ciliary length attenuates Shh signaling in choroid plexus development. **A-C.** Choroid plexus cilia transition from long, spreading cilia to short, clustered cilia from E12.5 to P14. Images of Immunostaining of cilia (Arl13b, **A** left), SEM (**B**), and TEM (**C**) analyses, as well as quantitation of axoneme morphology (**A** right), were shown at different developmental stages. The red arrow in **B**, cilia. **D.** Ependymal cilia elongate axonemes during maturation, as shown by immunostaining of Ac-α-Tub (ciliary axonemes), ZO1 (apical tight junction), and DAPI. **E.** The canonical Shh signaling decreases during choroid plexus development, as shown by a heatmap (left) of dynamically expressed Shh pathway components in RNA-seq analyses (E12.5, P0, P14, and 6-month choroid plexus), followed by the real-time PCR validation (right) of the reduced expression of the canonical Shh pathway targets (*Gli1*, *Gli2*, and *Hhip*) in development. n=3 for each developmental stage, error bars, sem. **F**. Immunostaining analyses show a reduced Smo translocation into choroid plexus cilia during embryonic and postnatal development (E14.5, P0, and P14). Pink arrows, Smo in ciliary axonemes; yellow arrows, Smo at ciliary base; white arrow, dispersed Smo in the cytosol; blue arrow, a CPEC without Smo staining.; n=3 for each developmental stage; E14.5 vs. P0, **** *P* < 0.0001; E14.5 vs. P14, **** *P* < 0.0001; P0 vs. P14, ns, *P* = 0.8938; one-way ANOVA and Sidak multiple comparisons tests. **G, H.** Choroid plexus cilia mediate an attenuated canonical and non-canonical Shh response during development (E14.5, P0, and P14), as measured by *Gli1* expression (**G**) and cAMP repression (**H**), respectively. **G**. E14.5 PBS vs. E14.5 Shh (n=3), \**P* = 0.0315; P0 PBS vs. P0 Shh (n=4), \**P* = 0.0171; P14 PBS vs. P14 Shh (n=3), n.s., unpaired two-tailed Student t-tests. **H.** E14.5 PBS vs. E14.5 Shh (n=5), \*\*\*\**P* < 0.0001; P14 PBS vs. P14 Shh (n = 4), \*\**P* = 0.0018; 4 month PBS vs. 4 month Shh (n = 4), n.s.; nested unpaired two-tailed Student’s t-test. **I, J.** Immunostaining demonstrated a declined Shh inhibitory effect on Aqp1 (**I**) and Atp1a2 (**J**) during choroid plexus development. Quantitation for apical Aqp1 and Atp1a2 is in Sup S4E, S4F. **K, L.** Aqp1 and Atp1a2 expression increase in choroid plexus during development, as measured by immunostaining (**K**, n=3) and Western blotting (**L**, n=2). **M.** Real-time PCR analyses demonstrated a decreased Shh mRNA expression in the choroid plexus during development (E12.5, P0, P14, and 6 months). n=3 for each developmental stage; error bars, s.e.m.. **N**. A model diagram illustrates a working model, in which the dynamic decrease of choroid plexus ciliary length in development attenuates the intensity of a non-canonical Shh signaling, causing an increased cAMP level, upregulating the apical expression of Aqp1 and Atp1a2, and ultimately, contribute to an increase in CSF production.

We then compared the transcriptomes of E12.5, P0, P14, and 6-month-old choroid plexus using RNA-seq (Sup Table S2), and identified a significant enrichment of Hh pathway components in genes downregulated during development (Sup Fig. S4C). The decline of Shh signaling in choroid plexus development was further validated by the gradual decrease of *Gli1*, *Gli2,* and Hedgehog-interacting protein 1 (*Hhip1*) expression (Fig. 4E), and by a diminishing ciliary Smo accumulation in shortened cilia (Fig. 4F, Fig. S4D). Hence, the decreased choroid plexus ciliary length in development is correlated with and likely gives rise to, a decline of Shh signaling activity. We compared the canonical Shh signaling (measured by *Gli1* expression) and non-canonical Shh signaling (measured by cAMP inhibition) in E14.5, P0, and P14 choroid plexus, and observed a developmentally regulated decline of signaling intensity (Fig. 4G, 4H). This decline was also evident in the inhibitory effect of Shh on apical Aqp1 and Atp1a2 expression (Fig. 4I, 4J, Sup Fig. S4E, S4F), consistent with an increase in Aqp1 and Atp1a2 expression in developing choroid plexus (Fig. 4K, 4L, Sup Fig. S4G), and ultimately, a rise in CSF production (Sup Fig. S4H).

Not surprisingly, the downregulated Shh signaling in choroid plexus development was not only solely mediated by ciliary length decrease. A dwindling Shh expression was also observed in the developing choroid plexus (Fig. 4M). Hence, both Shh production and ciliary sensing for Shh are developmentally dampened to collectively achieve a sharp decline of Shh signaling intensity, generating a rapid increase in cAMP and a surge in Aqp1 and Atp1a2 expression to ultimately promote CSF production (Fig. 4N).

## Discussion

Emerging evidence suggests that ciliary structure and function could be more heterogeneous and dynamic than previously recognized ^12,22,67–69^. Our studies revealed that choroid plexus cilia are sensory multicilia that transduce Shh signaling to activate Smo, downregulate cAMP and Aqp1/Atp1a2 expression, and ultimately regulate CSF production. A dynamic decrease of choroid plexus ciliary length during development leads to a decrease in Shh signaling intensity, and derepression of Aqp1 and Atp1a2, and possibly contributes to a surge in CSF production (Fig. 4N).

### Choroid plexus cilia have distinct ultrastructural features and function

Choroid plexus multiciila are a type of sensory cilia that also exhibit transient motility at the neonatal stage. This is consistent with their hybrid ciliary ultrastructure that resembles aspects of primary cilia and motile cilia. Consistently with a previous study ^22^, we observed an undefined, electron-dense central structure in the proximal axoneme segment of some cilia, possibly the remnant of what confers the transient motility at the neonatal stage ^15,23^. Mature CPECs exhibit a clustered basal body network, mediated through a heterogeneous assembly of SDAs. We speculate that SDAs play an important role in cilia clustering during their maturation, which could regulate the extent of sensory ciliary function and the intensity of signaling transduction.

The microtubule configuration in choroid plexus ciliary axonemes varies among species. The “9+2” choroid plexus cilia are observed in newly weaned pigs ^70^, Xenopus ^32^, and zebrafish ^3^, which, at least in some cases, direct CSF flow through their motility. The “9+0” configuration is reported in pig, mouse, rat and human ^71^ (Sup Fig. S2B), consistent with a sensory function. Interestingly, microtubule configuration in choroid plexus cilia also depends on developmental stages. In mice, where most choroid plexus cilia are “9+0”, a small subset are transiently “9+2” or “9+1” at the neonatal stage ^15^. The different choroid plexus ciliary ultrastructure across species and developmental stages likely implicate a dynamic and fast-evolving functionality.

### Choroid plexus cilia mediate a non-canonical Shh signaling to regulate CSF production

Two independent mouse cilia mutants, due to either *FoxJ1* or *Ift88* deficiency, exhibit neonatal hydrocephalus *in vivo*. In both, the volumetric CSF increase appears before the coordinated ependymal cilia beating initiates a CSF flow (Fig. 1C, 1D, S1D). Mature choroid plexus cilia transduce Shh signaling to activate a Smo/Gαi-dependent pathway, reduce cAMP, repress Aqp1 and Atp1a2 expression, and possibly decrease water, Na^+^ and K^+^ transport in CSF production. Hence, ciliopathy in the choroid plexus likely elevates CSF production, and ciliopathy in ependyma likely obstructs CSF circulation. Both could contribute to the neonatal hydrocephalus phenotype.

Choroid plexus cilia transduce both canonical and non-canonical Shh signaling. The *FoxJ1* deficient choroid plexus still produces a single or a small number of cilia that partially transduce the canonical Shh signaling via Gli-mediated transcription (Fig. 1A, S2I, S2J), but they no longer mediate any non-canonical Shh signaling. Hence, choroid plexus cilia are heterogeneous in their ciliogenesis and in their functions to transduce Shh signaling.

### Decreased ciliary length in choroid plexus development regulates Shh signaling intensity

While most cilia elongate axonemes during maturation ^20,21^, choroid plexus cilia decrease in length (Fig. 4A-4C). *Ho et al* observed a modest ciliary length decrease in postnatal, lateral choroid plexus (birth to 2 years) ^22^, yet the most rapid and significant ciliary length reduction occurs from E12.5 to P14, at least in the 4th ventricle choroid plexus (Fig. 4A-4C). This time window coincides with a decline of Shh signaling intensity, an elevated Aqp1 and Atp1a2 expression, and a rapid increase in CSF production. Two mechanisms contribute to a rapid decline of Shh signaling intensity in choroid plexus development. Shh expression declines significantly (Fig. 4M), and the shorter, mature choroid plexus cilia decreased Shh response (Fig. 4F, 4G, 4H). Hence, the developmentally regulated cilia length can fine-tune the intensity of a signaling pathway to achieve a dynamic regulation in choroid plexus function.

### Limitation of the Study

A significant limitation of this study is that the field does not have a feasible, quantitative assay to measure CSF secretion in neonatal mice. Due to this technical limitation, it is not possible yet to establish a direct *in vivo* functional link between choroid plexus cilia defects and an increased CSF production by CPECs. Throughout the paper, the authors have used the Aqp1 and Atp1a2 expressions in CPECs as a proxy for the secretary function of the choroid plexus. Furthermore, the mechanism through which cAMP regulates Aqp1 and Atp1a2 expression is unclear. Finally, while active Smo structurally resembles an active Gαi-coupled G-protein-coupled receptor (GPCR), it is unclear how Smo signals through Gαi protein in CPECs.

## Resource Availability section

### Lead contact

Further information and requests for resources and reagents should be directed to and will be fulfilled by the lead contact, Lin He.

### Materials availability

All data generated during this study are included in this published article. This study did not generate new unique reagents.

## Data and code availability

The developed analysis tools are available from the corresponding author upon request. Any additional information required to reanalyze the data reported in this paper is available from the lead contact upon request.

## Supporting information

Movie S1. A 3D reconstruction of P14 choroid plexus cilia

## Acknowledgments

We thank J Nealon, P Lu, M. Kinisu, T Machen, P Lishko, D Bautista, and C Ott, L. Liu, E Monuki, R. Fame, E Vladar and S Liddelow for technical input and stimulating discussion, K McDonald and P Kysar for SEM and TEM analyses, S Brody for *Foxj1^-/-^* mice, B. K Yoder for *Ift88^fl/fl^* mice, D Holtzman and Y Zhang for *FoxJ1-Cre* mice, the UC-Berkeley Electron Microscope Laboratory for assistance in electron microscopy sample preparation and data collection, and M Lehtinen for advice and reagents for the choroid plexus and CSF-related experiments. L.H. is a Thomas and Stacey Siebel Distinguished Chair Professor, supported by a Howard Hughes Medical Institute (HHMI) Faculty Scholar award, a Bakar Fellow award at UC Berkeley, and grants from the National Institutes of Health (1R01GM114414, R01NS120287) and from Nan Fung life sciences. R.S. was supported by a K99 Pathway to independent award by NIH K99HL128912. S.M. was supported by the Postdoctoral Fellowship Award from TRDRP (T30FT1019). S.P. C.S.X. and H.F.H. are supported by the Howard Hughes Medical Institute. Y. K Xiang is supported by the VA Merit grants IK6BX005753 and I01BX005100. A.Z. and S.U. are partially funded by the Philomathia Foundation. S.U. is supported by Lawrence Berkeley National Lab’s LDRD, Sloan Foundation, and the Chan Zuckerberg Initiative Imaging Scientist program. L.H. J.F.R. and S.U. are supported by the Chan Zuckerberg Biohub – San Francisco Investigator program.

## Author Contributions

Equal contributions were made by multiple co-first authors, whose findings were described in the chronicle order below. R.S. discovered the distinct choroid plexus ciliary ultrastructure, and then collaborated with S.P., C.S.X., and H.H. to perform FIB-SEM. S.P. prepared FIB-SEM samples, enabled precise targeting of regions of interest, and performed image acquisition and post-image processing in 42 imaging days. C.S.X. designed and built the enhanced FIB-SEM system, established post-image raw data registration and alignment pipeline, and optimized imaging conditions. S.P. and C.S.X. performed the initial data analyses and produced the first 3D movie. Subsequently, S.U. led S.J, A.Z., X.W., R.S., and P.L., to perform detailed FIB-SEM analyses and quantitation. S.U. developed algorithms for FIB-SEM analyses and supervised 3D image processing, segmentation, rendering, and movie production. S.J. rendered 3D models and produced 3D movies; A.Z., P.L, and X.W. handled 3D image segmentation, image processing, and 3D quantitation. G.C. and Y.W. performed TEM studies on the human choroid plexus.

R.S. identified *FoxJ1* as an essential regulator for choroid plexus ciliogenesis and discovered an early onset hydrocephalus in *FoxJ1^-/-^*mice; S.J. and R.S. discovered elevated Aqp1 expression in *FoxJ1^-/-^* mutants. S.M., M.F.W., W.C. and K.L. defined the exact onset of hydrocephalus in *FoxJ1^-/-^*mice and demonstrated the elevated Aqp1 expression with high-quality data.. S.M further characterized additional ion channels and transporters in mouse mutants with impaired choroid plexus cilia.

R.S. discovered the decreased ciliary length in developing choroid plexus. S.M. and W.C. provided quantitative data to validate this finding. S.J. Z.X. and L.H. performed an RNA-seq experiment, discovered and validated a dwindling Shh signaling in developing choroid plexus.

S.M. made a parallel finding using *in vitro* essays. R.S. and S.J. made the initial discovery of a dwindling Shh signaling in developing choroid plexus by Smo localization; S.M. perfected and expanded the data and produced final images and quantification for the manuscript.

R.S., with input from S.M., conceived the hypothesis that cAMP regulates Aqp1 downstream of Shh. R.S. and S.J. functionally validated the role of cAMP on Aqp1 expression. A.J., S.J. S.M., and K.Y. then characterized the non-canonical Shh-Smo-cAMP response mediated by choroid plexus cilia. S.J. initially discovered that choroid plexus cilia mediate Shh signaling to downregulate Aqp1. S.M. demonstrated the regulation of Aqp1 and Atp1a2 by a non-canonical Shh-Smo-Gαi signaling through choroid plexus cilia. S.M., with assistance from A.J., demonstrated that the Shh/cAMP/Aqp1-Atp1a2 signaling in choroid plexus attenuates in development, provided the key support that Shh represses Aqp1 and Atp1a2 via a non-canonical signaling, and generated all data for manuscript revision. S.M. R.S., S.J., S.P, and A.J. produced most data in the figures, R.S., S.M., and L.H. drafted the manuscript, S.P., and S.U. revised the manuscript, and all other co-authors proofread the manuscript.

## STAR★Methods

### Key resources table

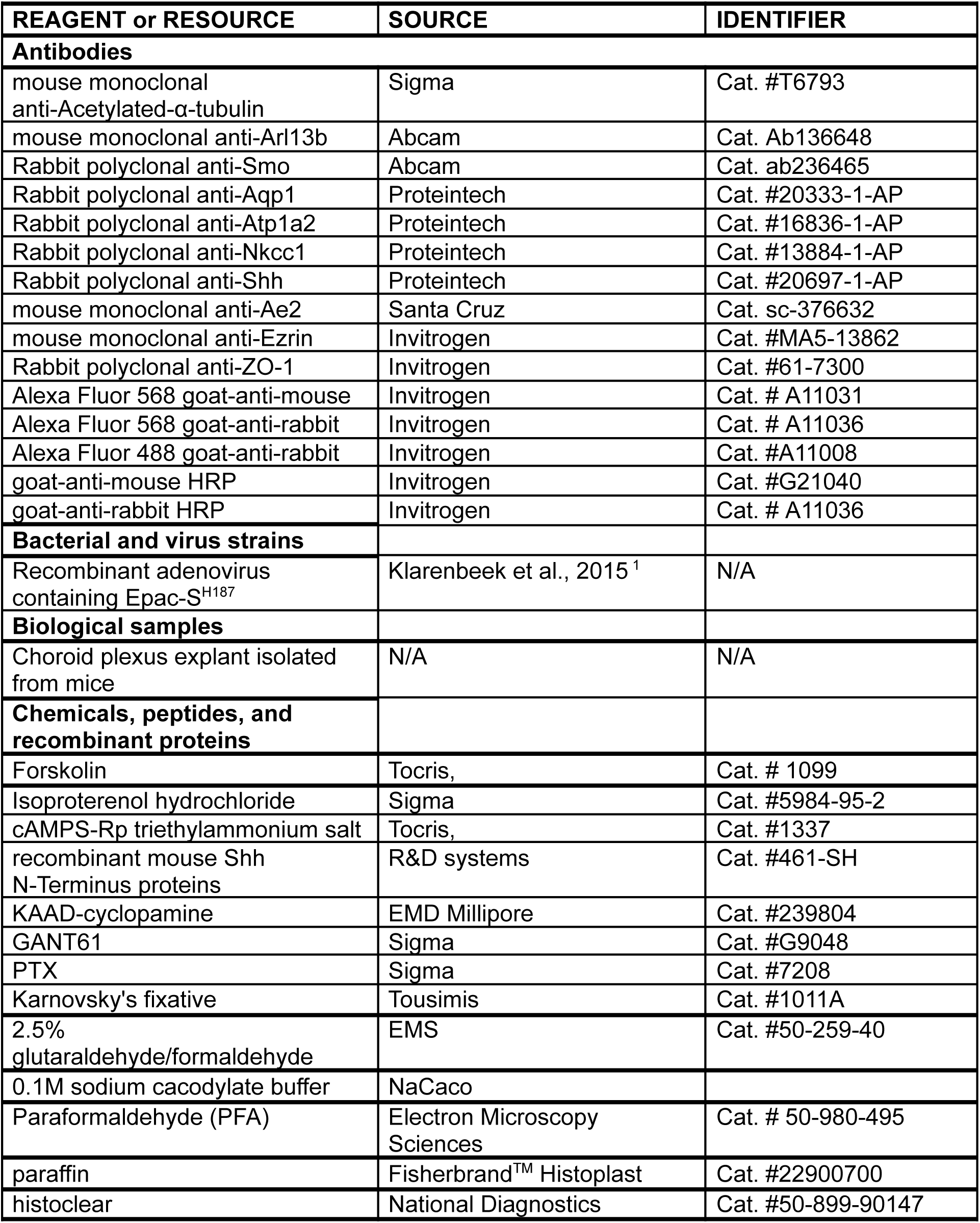
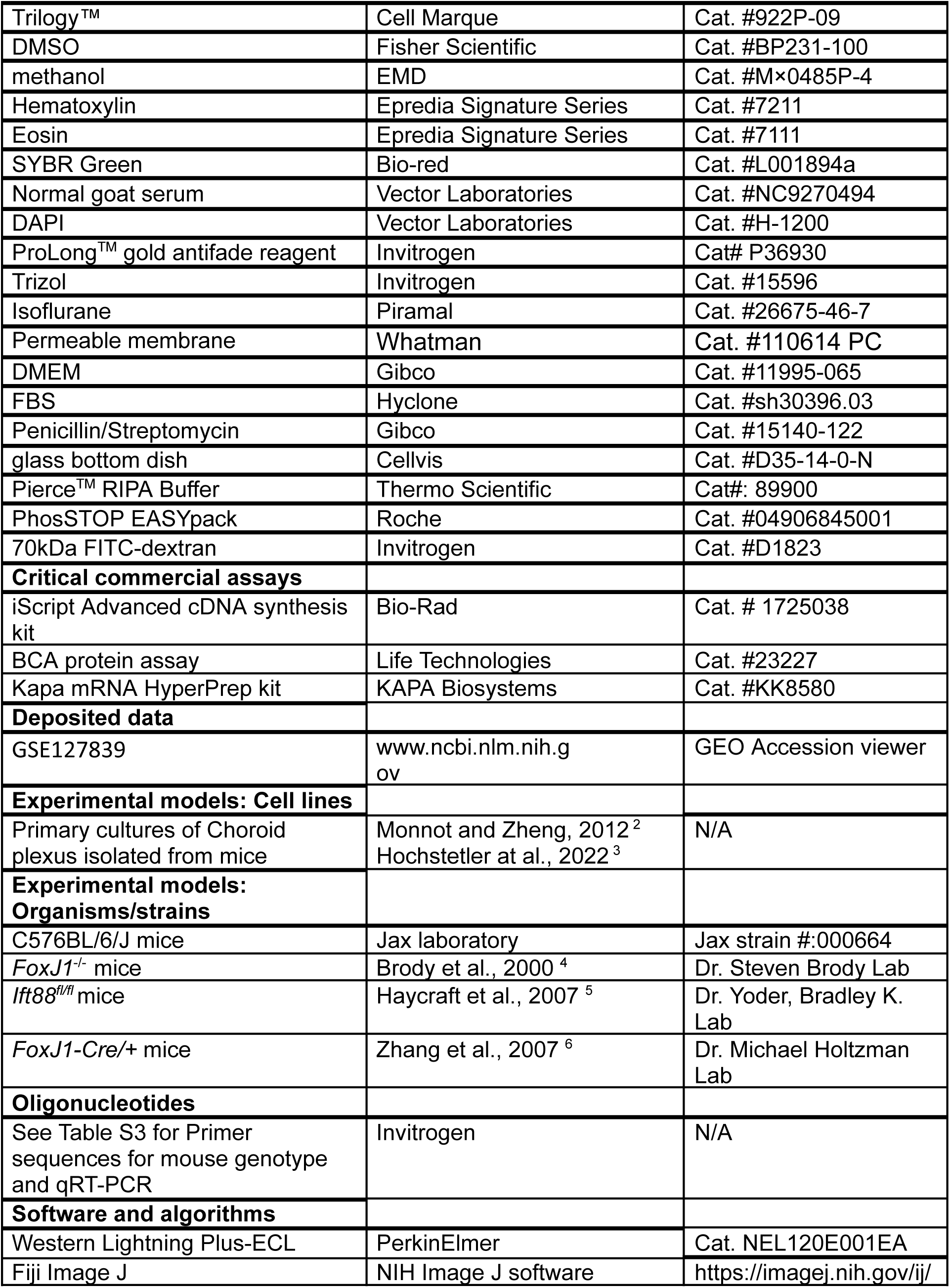
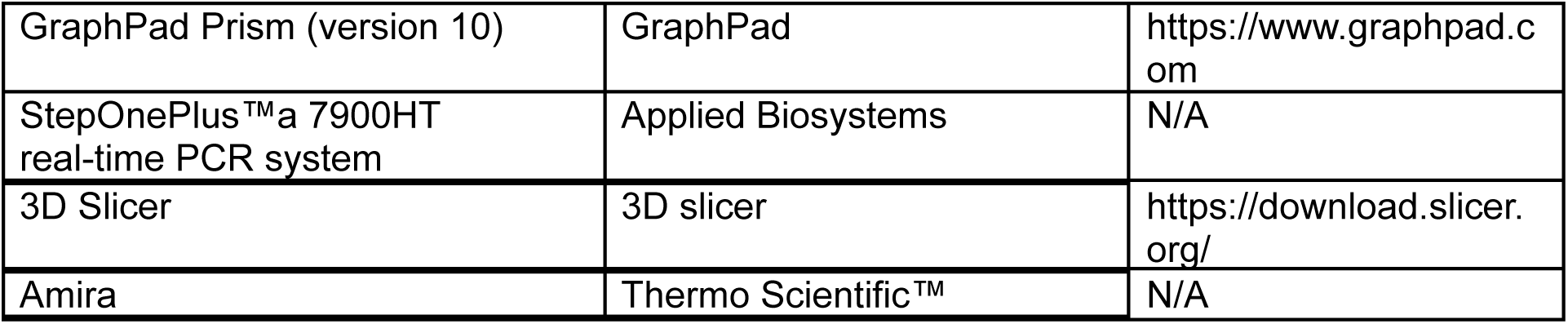

## Experimental model and study participant details

### Mouse breeding and genotyping

*FoxJ1*^-/-^ mice were a generous gift from Dr. Steven Brody ^4^; *Ift88^fl/fl^* mice were provided by Dr. Yoder, Bradley K. ^5^; *FoxJ1-Cre/+* mice were provided by Dr. Michael Holtzman ^6^. These mice were generated and maintained on a 129/B6 mixed genetic background and were housed in a non-barrier animal facility at UC Berkeley. All primer sequences for genotype are documented in Supplementary Table S3. All wild-type mice used in this study are C576BL/6/J mice (Jax strain #:000664), either purchased commercially or propagated in-house. The University of California, Berkeley’s Animal Care and Use Committee (ACUC) approved the animal protocol for this study. All animal experiments conform to the regulatory standards

## Method details

### Histology analyses

For all histology analyses, we perfused anesthetized mice with 4% paraformaldehyde (PFA) before surgical dissection. Whole brains were collected from littermate-controlled, wildtype and *FoxJ1*^-/-^ mice (P1, P3 and P7), or from littermate-controlled *Ift88^fl/fl^* and *FoxJ1-Cre/+; Ift88^fl/fl^* mice (P3 and P7). Samples were fixed overnight in 4% PFA (Electron Microscopy Sciences, Cat. # 50-980-495), cut into sagittal slices 2 mm in thickness, and processed and embedded with paraffin (Fisherbrand^TM^ Histoplast, Cat. #22900700)) by standard procedures ^7^. The paraffin sections 5-µm in thickness were stained with hematoxylin (Epredia Signature Series Hematoxylin, Cat. 7211) and eosin (Epredia Richard-Allan Scientific Signature Series Eosin-Y, 7111) to characterize the morphology of brain ventricles and choroid plexus epithelium.

### Immunofluorescence staining

For immunostaining on paraffin slides, paraffin-embedded sections, 5um in thickness, were deparaffinized by histoclear (National Diagnostics, Cat. 50-899-90147), rehydrated in a series of ethanol solutions (100% -> 95% -> 90% -> 70%) and water, treated with heat-induced antigen retrieval (Trilogy™, Cell Marque, Cat. 922P-09) and subjected to immunofluorescence staining. For whole-mount immunostaining, choroid plexus or ependymal tissues were fixed for 2 hours at −20 °C using 80% methanol (EMD, #M×0485P-4) in 20% DMSO (Fisher Scientific, #BP231-100). Fixed tissues were rehydrated in a series of methanol solutions (95%, 90%, 75%, 50%, 25% Methanol) and PBS (PH=7.4), for 5 min at room temperature for each condition, before immunostaining. In both whole mount and paraffin section immunofluorescence staining, rehydrated tissues/slides were blocked in 10% normal goat serum (Vector Laboratories, #NC9270494) in PBS (PH=7.4) for 1 hour at room temperature, incubated at 4°C overnight with primary antibodies (5% normal goat serum in PBS, PH =7.4), washed with PBS for 3 times at room temperature, and incubated with appropriate secondary antibodies (5% normal goat serum in PBS5, PH =7.4) at 4°C overnight. Finally, the samples were counterstained with 4’,6-diamidino-2-phenylindole (DAPI, Vector Laboratories, Cat. #H-1200) at 1µg/mL for 10 minutes at room temperature, before mounting with ProLong^TM^ gold antifade reagent (Invitrogen, Cat# P36930).

Primary antibodies include those against Acetylated-α-tubulin (Sigma, Cat# T6793, 1:300), Arl13b (Abcam, Cat. Ab136648, 1:400), ZO-1 (Invitrogen, Cat. #61-7300, 1:500), Aqp1 (Proteintech, Cat. #20333-1-AP, 1:200), Atp1a2 (Proteintech, Cat. #16836-1-AP, 1:200), Ae2 (Santa cruz, sc-376632, 1:200), Smo (Abcam, Cat. ab236465, 1:100), Ezrin (Invitrogen, Cat. #MA5-13862, 1:200), Shh (Proteintech, Cat. #20697-1-AP, 1:200). The secondary antibodies include goat-anti-mouse 568 (Invitrogen, Cat# A11031, 1:400), goat-anti-rabbit 568 (Invitrogen, Cat. # A11036, 1:400), goat-anti-rabbit 488 (Invitrogen, Cat. #A11008, 1:400).

### Confocal imaging and quantification

For immunofluorescence on paraffin sections, confocal images were captured using spinning disc confocal microscopy (Nikon TE2000-E or Andor BC43) as a single slice width at 0.3μm. The integrated density of cellular fluorescence signals of a specific protein, such as Aqp1 and Atp1a2, was measured by ImageJ for one whole cell and quantified based on the method previously described ^8^. Briefly, for each biological sample, 4-6 images were imaged for analysis. The sum of Aqp1/ATp1a2 immunofluorescence intensities for each sample was divided by the number of nuclei (counted by DAPI staining) to quantify the signals for each cell.

For whole mouse immunofluorescence staining, confocal images were Z-stacked for quantification using ImageJ. The apical fluorescence intensity of a protein, such as Aqp1 or Atp1a2, is measured by the stacked images positive for Ezrin co-staining whenever possible. However, Ezrin expression alters in response to Shh or FSK treatment (Sup Fig. 2F), making Ezrin an undesirable internal reference for some experiments. When Ezrin cannot be used as a reference, apical fluorescence intensity is measured by 7.5 μm-depth (∼25 images along the z-axis, 0.3um per image) stacked images above DAPI-stained nuclei, and DAPI is used as an internal reference across samples. The total cellular fluorescence intensity of a protein is measured by a z-stack of all slices stacked in the full depth of a cell and then normalized to that of nuclei (DAPI) as an internal control. Each dot in our quantitation plots represents the ratio of Aqp1/DAPI averaged from 3-6 representative images in a given biological sample. For fluorescence intensity measurement by Image J in each channel, cellular fluorescence intensity = Integrated Density – (Area of selected cell X Mean fluorescence of background readings) (https://theolb.readthedocs.io/en/latest/imaging/measuring-cell-fluorescence-using-imagej.html) For quantitative comparisons, an average of 300∼600 cells were measured for fluorescence intensity in each biological sample/condition.

### Real-time PCR analyses

Total RNA was isolated by Trizol (Invitrogen, Cat. #15596) from choroid plexus tissues following the manufacturer’s protocol. For mRNA quantitation, RNA was reversely transcribed using an iScript Advanced cDNA synthesis kit (Bio-Rad, Cat. # 1725038) with random primers. Resulted cDNA was then subjected to SYBR Green-based, real-time PCR analyses (Bio-rad, Cat. # L001894a) using StepOnePlus™a 7900HT real-time PCR system (Applied Biosystems). *β-Actin* was used as an endogenous control. All primer sequences for real-time PCR analyses are documented in Supplementary Table S3. The relative gene expression of genes was plotted by using the (ΔΔct) or 2^-ΔΔct^ method to show the fold change of mRNA expression.

### MRI-based CSF measurement

Magnetic Resonance Imaging (MRI) was performed using a 7.0 Tesla Bruker Pharmascan system located at the small animal imaging center at UC Berkeley. Neonatal or postnatal mice were anesthetized with isoflurane (Piramal, Cat. 26675-46-7) and positioned in a mouse brain surface array coil (Bruker 2×2) designed for mouse brain imaging. The preparation was inserted into a transmit-only volume excitation coil installed in the magnet. After localizer scans, each mouse was imaged in the coronal and the sagittal views using Rapid Acquisition with Relaxation Enhancement (RARE) pulse sequence, with TR = 5000 ms, TE: T1= = 12 ms, T2 = 60 ms, FOV = 16mm, matrix = 128×128 (resolution = 0.125 mm/pixel), 29 slices = 29 with a thickness of 0.25mm for P0, P1 and P3 mice, and 0.35mm for P14 mice, RARE factor = 8, bandwidth = 81521.79 Hz. Scan time for each image was 10 min 40 sec. Each image voxel represents 0.0039 mm^3^ in volume. Total CSF volume is measured based on sagittal MRI (µL or mm^3^) = sum of total area (mm2) x thickness (mm).

### Scanning Electron Microscopy (SEM) analysis

Choroid plexus from E12.5, P0, and P14 wildtype mice were fixed overnight in Karnovsky’s fixative (Tousimis, Cat. #1011A) in 0.1 M sodium phosphate buffer (Sorenson’s) under room temperature. The fixed tissue was then washed with Sorenson’s sodium phosphate buffer and post-fixed in 1% OsO4 in Sorenson’s for 1 hour. Tissue was dehydrated by passing through a graded series of ethanol solutions, then critical point dried using a Tousimis 931 Super Critical Point Dryer. The tissue was mounted on aluminum stubs and sputter coated with gold using a PELCO SC-7 coater. SEM images were viewed and captured on a FEI XL30 TMP SEM in the electron microscopy facility of the University of California at Davis.

### Transmission Electron Microscopy (TEM) analysis

Mouse and human choroid plexus tissues were fixed in 2.5% glutaraldehyde/formaldehyde in 0.1M sodium cacodylate buffer, pH 7.2 (EMS, Hatfield, PA, USA) for at least 1 hr under room temperature. Samples were rinsed (3× 10 min) in 0.1M sodium cacodylate (NaCaco) buffer, pH=7.2, and immersed in 1% osmium tetroxide with 1.6% potassium ferricyanide in 0.1M NaCaco buffer for 1 hour. Samples were subsequently rinsed (3×10 min at room temperature) in 0.1M NaCaco buffer and then in distilled water (3×10 min at room temperature) before being subjected to an ascending ethanol gradient (10min; 35%, 50%, 70%, 80%, 90%, 100%) followed by pure acetone (2× 10 min). Samples were progressively infiltrated while rocking with Epon resin (EMS, Hatfield, PA, USA) and polymerized at 60 °C for 24-48 hours. 70 nm ultrathin sections were cut using a Reichert-Jung Ultracut E ultramicrotome and collected onto formvar-coated 200 mesh copper grids, or 50 nm serial sections were picked up on 1 x 2 mm slot grids covered with 0.6% Formvar film. Grids were post-stained with 2% uranyl acetate followed by Reynold’s lead citrate, for 5 min each. Sections were imaged using a Tecnai 12 120kV TEM (FEI, Hillsboro, OR, USA) and data was recorded using an UltraScan 1000 with Digital Micrograph 3 software (Gatan Inc., Pleasanton, CA, USA) ^9^.

### Focus ion beam scanning electron microscopy (FIB-SEM)

#### a.#EM Sample preparation

Choroid plexus samples were first fixed in 2.5% formaldehyde and 2.5% glutaraldehyde in 0.1M Sodium Cacodylate Buffer, PH=7.4 (Emsdiasum, Cat# 15949), and then stained with a modified osmium-thiocarbohydrazide-osmium (OTO) method in combination with microwave assisted processing, followed by HPF-FS as previously described ^10^. Briefly, samples were subjected to HPF-FS, and freeze-substituted with 4% osmium tetroxide, 0.1% uranyl acetate, and 5% ddH2O in acetone; then thin embedded and polymerized in Durcupan resin.

#### b.#FIB-SEM Sample preparation

Durcupan-embedded mouse choroid plexus samples were each initially mounted to the top of a 1 mm copper post, which was in contact with the metal-stained sample for belter charge dissipation. Subsequently, the vertical sample post was trimmed to a small block containing a Region of Interest (ROI) with a width perpendicular to the ion beam, and a depth in the direction of the ion beam, as previously described ^11^. The trimming was guided by X-ray tomography data obtained by a Zeiss Versa XRM-510 and optical inspection under a microtome. Thin layers of conductive material of 10-nm gold followed by 100-nm carbon were coated on the trimmed samples using a Gatan 682 High-Resolution Ion Beam Coater. The coating parameters were 6 keV, 200 nA on both argon gas plasma sources, and 10 rpm sample rotation with a 45-degree tilt.

#### c.#3D large volume high-resolution FIB-SEM imaging

FIB-SEM prepared P14 mouse choroid plexus sample was managed by a customized Zeiss NVision40 FIB-SEM system previously described ^12,13^. Each sample was biased at 400 V to improve image contrast by filtering out secondary electrons ^12^. The block face was imaged by a 3 nA electron beam with 1.5 keV landing energy at 1.25 MHz. The x-y pixel resolution was set at 6 or 8 nm. A subsequently applied focused Ga+ beam of 27 nA at 30 keV strafed across the top surface and ablated away 6 or 8 nm of the surface. The newly exposed surface was then imaged again. The ablation-imaging cycle continued approximately once every two minutes for two weeks to complete FIB-SEM imaging of one sample, maintaining an 8 x 8 x 8 nm³ voxel resolution throughout the entire volume. Each acquired image stack formed a raw volume, followed by post-processing involving image registration and alignment using a Scale Invariant Feature Transform (SIFT) algorithm. The aligned stacks have a final isotropic volume of 60 x 60 x 60 µm³, enabling viewing in any arbitrary orientation.

#### d.#FIB-SEM Analysis & Visualization

##### d.1#Region of Interest (ROI) selection

All FIB-SEM volumes were first loaded into Fiji, and the region of interest (ROI) containing the cilia/basal body region was selected from the tip of the cilia to the proximal end of the basal bodies in each cell to generate the 3D volume by using software 3D Slicer. Thermo Scientific™ Amira 2019.2-2019.4 (Amira) software was used to navigate the large FIB-SEM data files across the different time domains to identify regions with relevant cellular morphology. The raw data was first visualized in 3D using Amira, and the use of median filters along with partial transparency enabled efficient identification and cropping of ROIs containing relatively high signal-to-noise-ratio basal bodies and cilia. 17 ROIs were selected and cropped in the time domain P14. Raw FIB-SEM data varied in quality and contrast across ROIs and time domains. 15 ROIs from each time domain were selected for quality and high contrast between the basal bodies and their surroundings. High-quality ROIs had the fewest FIB-SEM drift artifacts shearing the cilia or basal bodies and were subjected to further analysis for SDA, basal body number, and cilia length.

##### d.2#Segmentation of individual basal bodies and cilia

Each basal body was segmented as a secondary ROI. The SDA number of each basal body was confirmed by both volume rendering and surface view. Ilastik (version 1.3.3b2) was used to segment basal bodies and cilia from their surroundings ^14^. Accuracy and computation time were balanced to produce probability mask files. Using custom MathWorks® MATLAB (R2019a) software scripts the basal body and cilia were separated using their probability mask Ilastik outputs. Results were manually curated for accuracy in Amira, Fiji ^15^, and ITK-SNAP ^16^. Depending on the severity of missing data in the basal body and cilia probability masks, ITK-SNAP (version 3.6.0) or Ilastik was used for manual image curation.

##### d.3#Movie generation

The movies were first generated in Amira using cropped volumes, 3D median filter processed volumes, and segmented cilia/basal bodies. Adobe Premiere Pro CC 2019 was used to compress the movies.

### Choroid plexus explant culture and treatment

Intact choroid plexus was dissected from the fourth ventricles in the brain of E14.5, P0, P7, or P14 mice, washed in 1× PBS, and then transferred onto a permeable membrane (Whatman, Cat. # 110614 PC) that floated in the choroid plexus culture media containing DMEM (Gibco, Cat. #11995-065) with 10% FBS (Hyclone, Cat. #sh30396.03) and 1% Penicillin/Streptomycin (Gibco, Cat. #15140-122). Freshly isolated choroid plexus explants were treated for 24 hours with Forskolin (25μM in DMSO, Tocris, Cat. # 1099), recombinant mouse Shh N-Terminus Protein (20nM in PBS, R&D systems, Cat #, 461-SH), KAAD-cyclopamine (200nM in DMSO, EMD Millipore, Cat. #239804), GANT61 (25μM in DMSO, Sigma Cat. #G9048), PTX (500 ng/mL in H_2_O, Sigma Cat. #7208), cAMPS-Rp triethylammonium salt (50μM in H_2_O, Tocris, Cat. #1337), SANT-1 (250nM in DMSO, Tocris, Cat. #1974) or any combinations for 24h before subjecting to downstream analyses. Choroid plexus explants treated with PBS, DMSO, or H_2_O were used as controls depending on the experiments.

### cAMP measurement in choroid plexus explants using a FRET Assay

Choroid plexus tissue explants were prepared as described above. 4th ventricle choroid plexus explants from E14.5, P0, P14, or 4 months mice were infected with a recombinant adenovirus containing Epac-S^H187^, an EPAC-based cAMP FRET biosensor ^1,17^. The FRET biosensor, Epac-S^H187^, consists of the cAMP-binding, Rap-1 activating protein EPAC, sandwiched between donor- and acceptor fluorescent proteins, CFP (cyan fluorescent protein) and YFP (yellow fluorescent protein) ^1^. The binding of cAMP to the EPAC domain induces conformational changes that decrease FRET between YFP and CFP. After 24 hours of viral incubation, infected explants were transferred into a glass bottom dish (Cellvis, Cat#: D35-14-0-N) coated with matrigel (Corning, Cat. #354230), incubated in choroid plexus culture media for another 16-20 hours. ∼10 min before the FRET recording, the culture media was replaced with a PBS-based culture condition with an appropriate drug treatment scheme.

In control experiments, the baseline YFP/CFP ratio would be recorded for 3 minutes, after choroid plexus culture media was replaced by a PBS-based culture condition. The samples were then stimulated by Isoproterenol hydrochloride (ISO, 1 μM in PBS, Sigma Cat. #5984-95-2), a non-selective β-adrenergic agonist, to stimulate cAMP production. The alteration of the cAMP level was recorded by the YFP/CFP ratio until the recorded data reached the plateau. This process usually takes up to 10 minutes. After reaching the plateau, the samples were stimulated with forskolin (FSK, 10uM, Sigma Cat. #66575-29-9), an adenylyl cyclase activator, to maximally induce cAMP production. FRET images were acquired on a Zeiss AX10 microscope with a 40x/1.3 oil-immersion objective lens (Oberkochen, Germany) and a cooled charge-coupled device (CCD) camera. Dual emission ratio imaging was acquired with a 420DF20 excitation filter, a 450DRLP dichroic mirror, and two emission filters (475DF40 for cyan and 535DF25 for yellow). The acquisition was set with 200ms exposure in both channels and 20s elapses, and lasted 30-40 minutes for each sample. FRET image acquisition and intensity ratio measurement (YFP/CFP) were conducted through Metafluor software (Molecular Devices, Sunnyvale, CA). The background was subtracted in images from both channels, and yellow to cyan (YFP/CFP) ratios were calculated at different time points. YFP/CFP ratios were normalized to baseline (before isoproterenol stimulation) and maximal changes in YFP/CFP ratios were plotted on the bar graphs.

To test whether Shh can decrease ISO-induced cAMP production, choroid plexus explants were incubated with recombinant mouse Shh N-Terminus protein in PBS for 10 min before the recording (10nM in PBS, R&D systems, Cat #, 461-SH). The recording procedures are described in the control experiments.

To test whether the inhibition of Smo, Gli transcription factors or Gαi impairs the inhibitory effects of Shh on ISO-induced cAMP production, the samples were pretreated with KAAD-cyclopamine (200nM in DMSO, EMD Millipore, Cat. #239804) in PBS for 3h, with GANT61 (25μM in DMSO, Sigma Cat. #G9048) in PBS for 10 minutes, or with PTX (500 ng/mL in H_2_O, Sigma Cat. #7208) in PBS for 3h, respectively. Following the incubation, Shh N-Terminus protein was subsequently added to the PBS-based culture for 10 min before recording. The recording procedures are described in the control experiments.

### Immunoblotting

4th ventricle cChoroid plexus samples were collected from E14.5, P0 and P14 wildtype and *Foxj1^-/-^* mice. For E14.5 and P0, 3 and 2 choroid plexus collections were pulled as one biological sample, respectively. Each sample was homogenized in protein lysis buffer (Pierce^TM^ RIPA Buffer, Thermo Scientific, Cat#: 89900) that contained 1x phosphatase inhibitor cocktail (PhosSTOP EASYpack, Roche, Cat #: 04906845001) and 1x protease inhibitor cocktail (cOmplete Mini, EDTA-free, Roche, Cat#: 11836170001), quantified by a standard BCA protein assay (Life Technologies, Cat #: 23227), and resolved on 10% SDS-PAGE gel, then transferred to a 0.45 μm nitrocellulose membrane. After blocking with 5% non-fat milk in TBST (Tris-buffered saline, 0.5% Tween 20) at room temperature for 1h, the membrane was incubated overnight at 4°C with appropriate primary antibodies diluted in 5% non-fat milk in TBS-T. The membrane was subsequently washed with TBS-T three times for 10 minutes at room temperature, and then incubated with appropriate horseradish peroxidase (HRP)-conjugated secondary antibodies including goat-anti-mouse HRP (Invitrogen, Cat. #G21040) and goat-anti-rabbit HRP (Invitrogen, Cat. # A11036, 1:400) (Factinisher Scientific Cat. #. 31460 and Cat. G-21040) for 1 hour at room temperature. After three TBS-TPBS washes, chemiluminescence detection was performed using Western Lightning Plus-ECL (PerkinElmer, Cat. NEL120E001EA). β-Actin was used as a protein loading control. Primary antibodies used in this experiment include mouse monoclonal antibody against mouse β-actin (Proteintech Cat. 66009-1-Ig, 1:2000), rabbit polyclonal antibody against Aqp1 (Proteintech, Cat. #20333-1-AP, 1: 1000), rabbit polyclonal antibody Atp1a2 (Proteintech, Cat. #16836-1-AP, 1: 1000). The optical density of the band was analyzed using the NIH Image J software (https://imagej.nih.gov/ij/). The arbitrary units (A.U.) were defined as the ratio of the optical density of the protein of interest over the beta-actin control.

### CPEC Cerebrospinal-Fluid Secretion

To assess CSF secretion by cultured CPECs, P1 choroid plexus explants were isolated ^2^ and cultured on permeable, polyester 24-well transwell filter plates coated with collagen for 12 days ^3^. CPECs were washed three times in CSF secretion buffer (CSFB, in mM: 122 NaCl, 4 KCl, 1 CaCl2, 1 MgCl2, 15 Na-HCO3, 15 HEPES, 0.5 Na2HPO4, 0.5 NaH2PO4 and 17.5 glucose, pH7.3, 37°C, with 5 μg/ml insulin added ^18^. CPECs were preincubated with CSFB for one hour and replaced with CSFB containing 1.0 μM 70 kDa fluorescent-dextran (FITC-dextran) at both apical and basolateral chambers. Secreted CSF volume (in μl/cm^2^) was calculated as a function of FITC-dextran concentration change. Filters were incubated at 37°C, 5% CO2, and 95% relative humidity for 1 h ^18^. 150 μl samples were analyzed in a fluorescent plate reader. CSF volumes secreted were then calculated according to the equation ^20^.

Vc = {V_0_ x [C_Dex_ (initial) - C_Dex_ (final)] } / C_Dex_ (final)

Where, Vc = CSF volume [μl], C_Dex_ (initial) = initial FITC-dextran concentration [μM], C_Dex_ (final) = final FITC-dextran concentration [μM], V_0_ = initial volume applied [μl].

### RNA-seq

Using RNA-seq, we characterized dynamic gene expression changes in the choroid plexus during 4 different developmental stages, including E12.5, P0, P14, and 6 months. 100 ng of total RNA from the choroid plexus at each stage was used for RNA-seq library construction using the Kapa mRNA HyperPrep kit (KAPA Biosystems, Cat. #, KK8580). The fast data from RNA-seq were processed by Kallisto for quantification ^19^. Spearman correlation was calculated to identify genes with consistent increasing or decreasing expression patterns along the developmental time course for the choroid plexus. The heatmap plots produced in R. Fisher-exact tests were used to measure the significance of the overlapping between gene sets. DAVID bioinformatics tools were used for pathway and function enrichment analysis ^20^.

### Quantification and statistical analysis

Statistical analysis was performed using GraphPad Prism (version 10). Appropriate statistical tests were selected based on the distribution of data and sample size. Data are presented as mean ± standard error of the mean (s.e.m) or mean ± standard deviation of the mean (s.d). *P* values in the manuscript present the following star code: ns: p > 0.05 (non-significant),* *P* < 0.05, ** *P* <0.01, *** *P* <0.001, **** *P* < 0.0001. The statistical tests used in each figure are listed below.

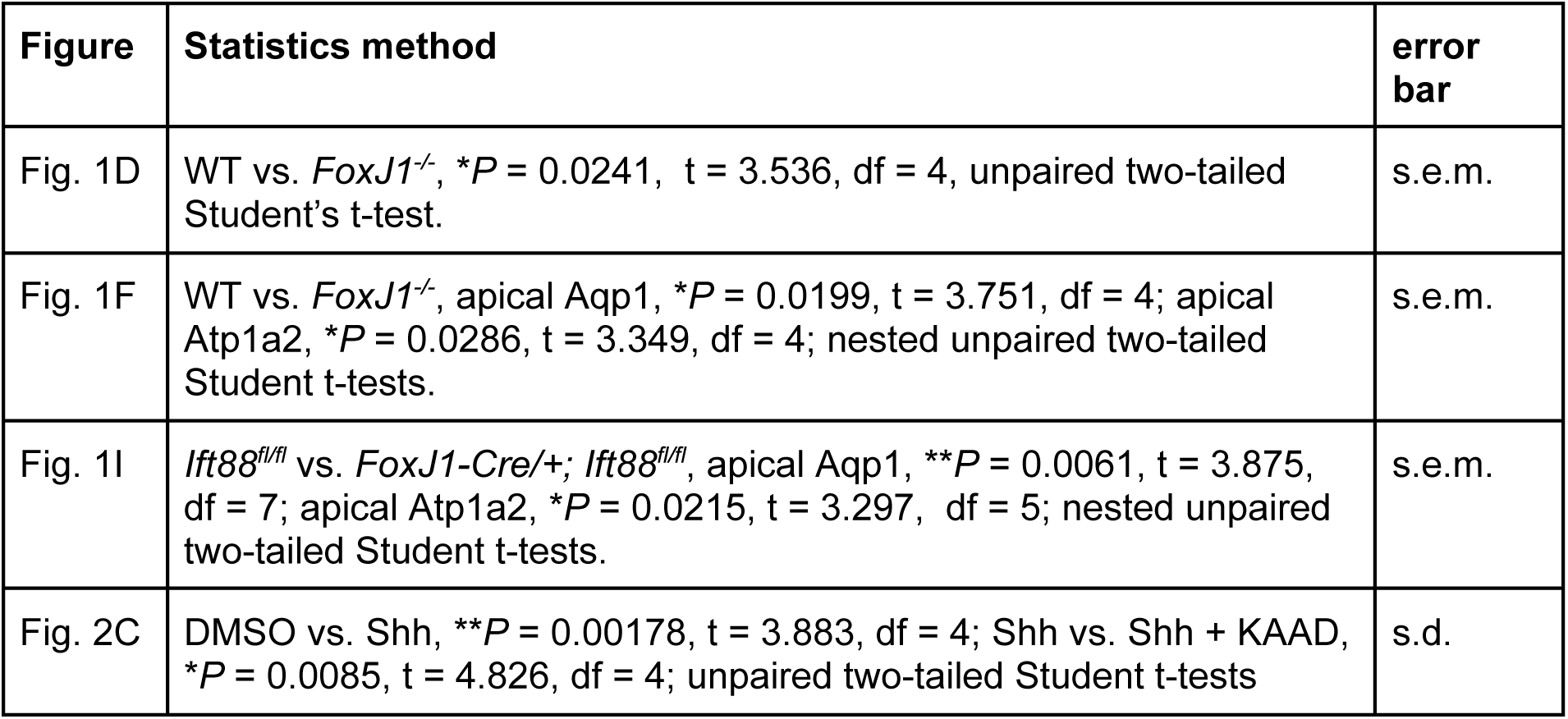
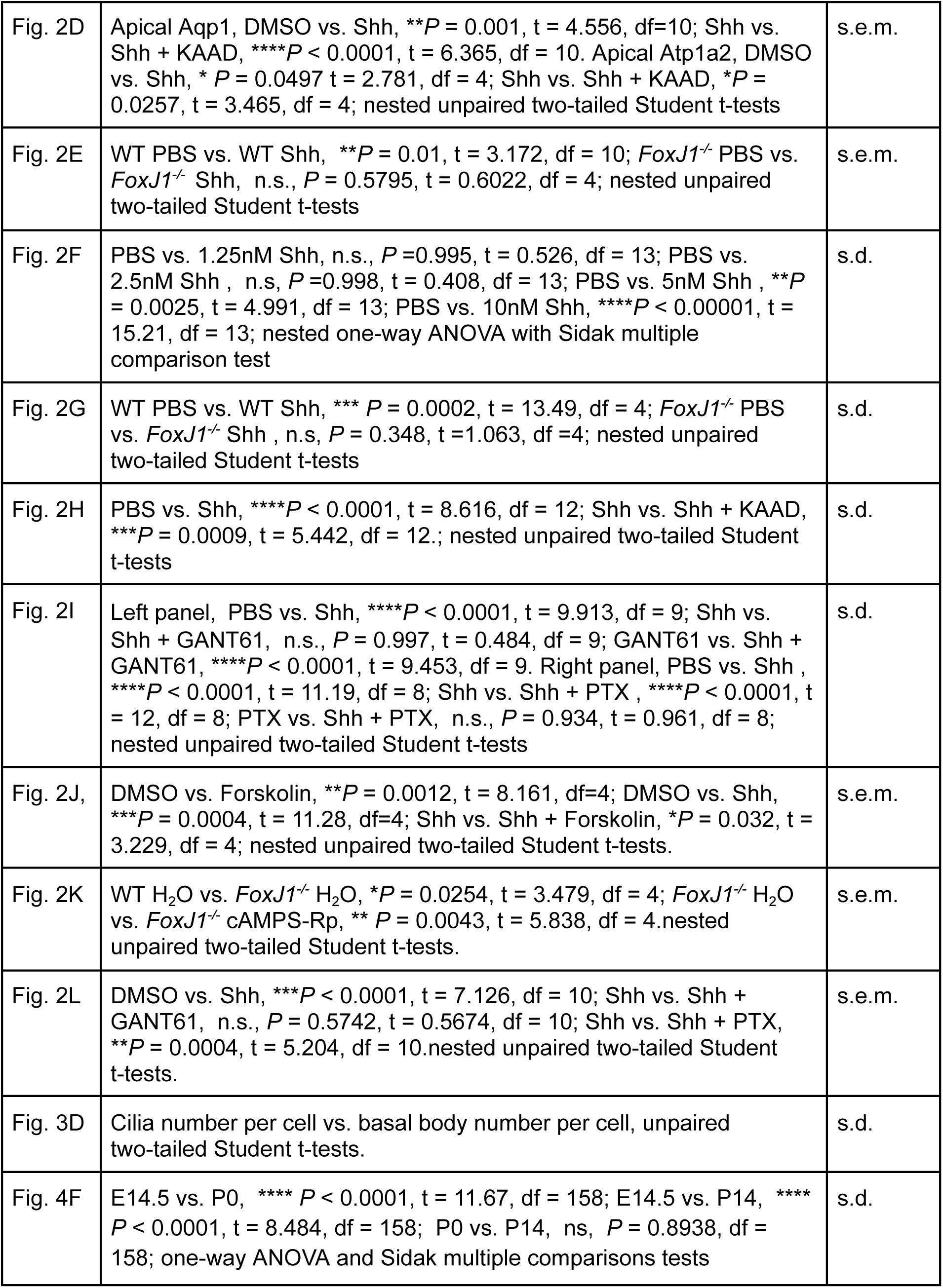
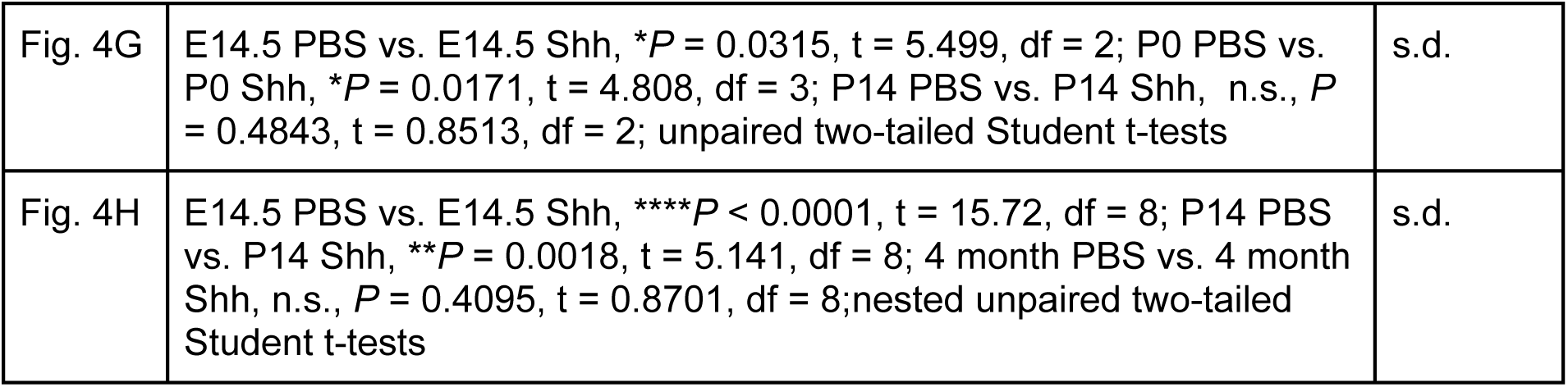

## Supplementary figure legend

**Supplementary Figure S1.**
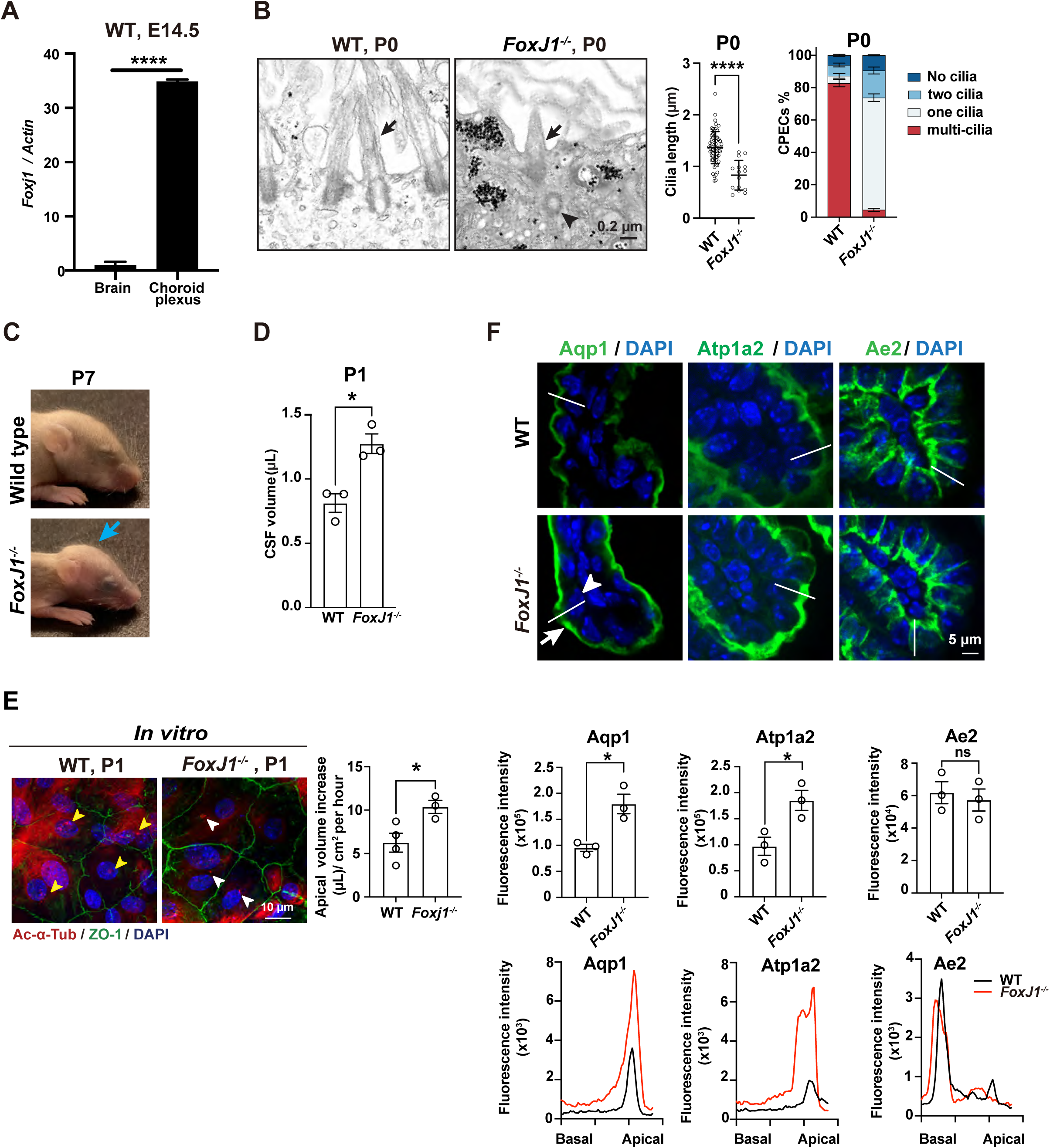
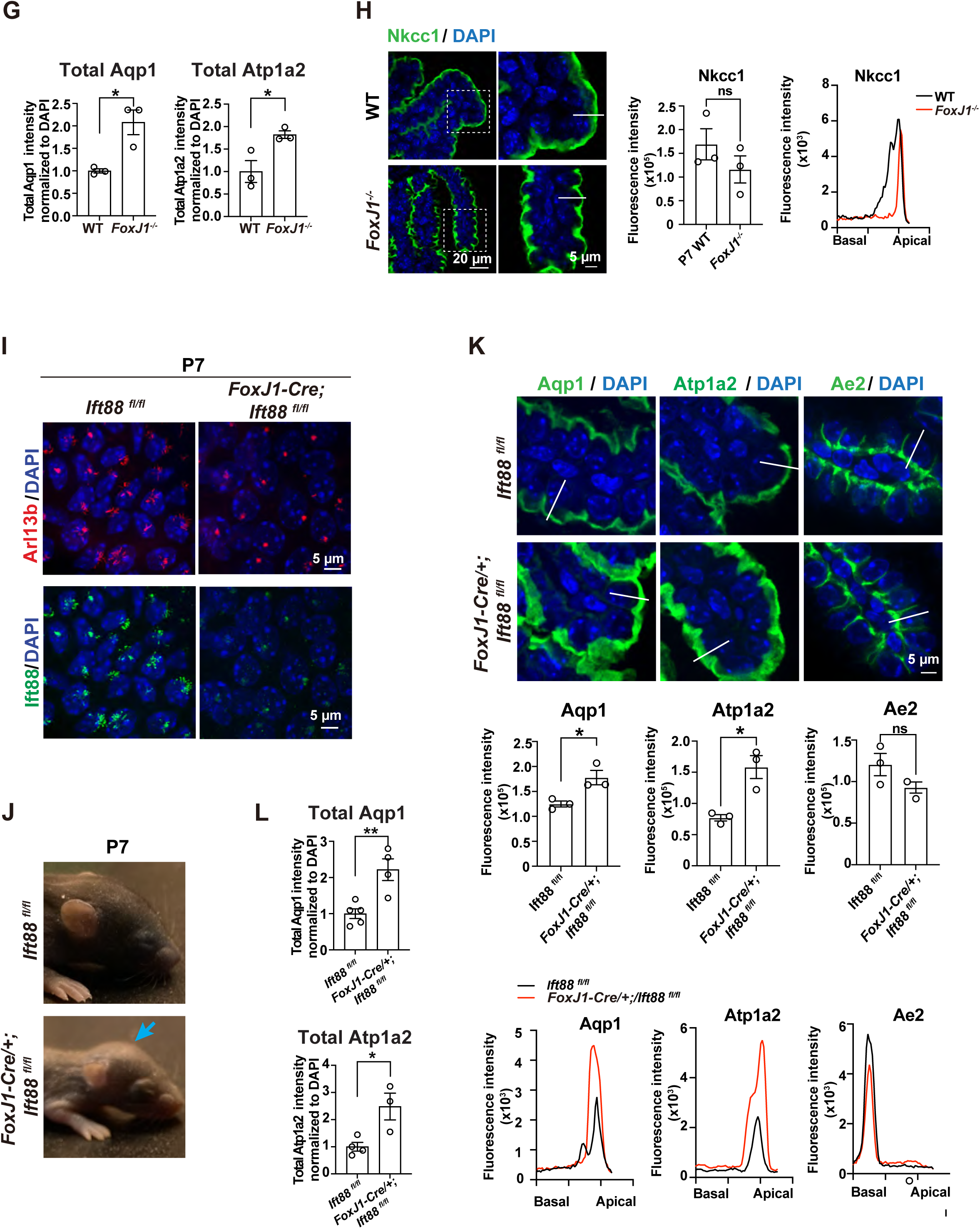
*Foxj1* deficiency impairs choroid plexus ciliogenesis and causes hydrocephalus. **A.** *FoxJ1* is enriched in the choroid plexus of the E14.5 fetal brain. Choroid plexus and whole brain tissue were each collected from 2 E14.5 embryos and subjected to real-time PCR analyses for *FoxJ1*. Error bars, sem; \*\*\*\**P* = 1.25 × 10^-5^, t = 282.9, df = 2; unpaired two-tailed Student’s t-test. **B.** In TEM analyses, *FoxJ1^-/-^* choroid plexus cilia are characterized by fewer and shorter ciliary axonemes and the presence of daughter centrioles. Black arrows, ciliary axonemes; black arrowhead, daughter centriole; scale bar, 0.2μm**. Middle**, cilia length measured by TEM is significantly reduced in P0 *FoxJ1^-/-^* choroid plexus, 1.365μm in P0 wild-type (n=64) vs. 0.831μm in P0 *FoxJ1^-/-^* (n=15), \*\*\*\**P* < 0.0001, t = 6.058, df = 77; unpaired two-tailed student t-test. **Right**, the quantification of cilia numbers patterns in CPECs from Fig 1A. n=3; error bars, s.e.m. CPECs with multi-cilia, wild-type vs. *FoxJ1*^-/-^, \*\*\*\**P* < 0.0001, t = 29.82, df = 8; CPECs with one cilia, wild-type vs. *FoxJ1*^-/-^, \*\*\*\**P* < 0.0001, t = 24.68, df = 8; CPECs with two cilia, wild-type vs. *FoxJ1*^-/-^, \**P* = 0.0192, t = 3.854, df = 8; CPECs with no cilia, wild-type vs. *FoxJ1*^-/-^, n.s. *P* = 0.6568, t = 1.285, df = 8; two-way ANOVA multiple comparisons. **C.** P7 *FoxJ1*^-/-^ mice exhibit a bulging forehead, consistent with the hydrocephalus phenotype. **D.** MRI analysis reveals an increased CSF volume in P1 *FoxJ1*^-/-^ mice. Quantitative measurements of CSF volume were shown, n=3; error bars, s.e.m.; wild-type vs. *FoxJ1*^-/-^, \**P* = 0.011, t=4.433, df=4; unpaired two-tailed Student’s t-test. **E.** An *In vitro* ChP culture model for CSF secretion. **Left,** *In vitro,* ChP culture recapitulates that Foxj1 deficiency decreases cilia number and ciliary length. Representative images are shown for Ac--Tub (red, ciliary axonemes), ZO-1(green), and DAPI (blue). **Right,** P1 WT and P1 *FoxJ1^-/-^* ChP epithelial cells were cultured for 12 days. ChP epithelial cell secretion volume was measured by FITC-dextran concentration change in one hour. CSF secretion, WT (n=4) vs. *FoxJ1*^-/-^ (n=3), \**P* = 0.0348, t = 2.874, df = 5; unpaired two-tailed Student t-test**. F.** *FoxJ1* deficiency increases the total expression of Aqp1 and Atp1a2. Three pairs of independent, littermate-controlled wild-type and *FoxJ1*^-/-^ mice were quantified for total Aqp1, Atp1a2, and Ae2 expression using immunostaining. **Top**, a zoom-out image of Aqp1, Atp1a2 and Ae2 in wild-type and *FoxJ1*^-/-^ choroid plexus. **Middle**, quantitation of relative fluorescence intensity of Aqp1, Atp1a2, and Ae2 in P7 WT vs. *FoxJ1*^-/-^ choroid plexus. Each dot is the average intensity from 3-6 images in one biological sample, n=3. Aqp1, WT vs. *FoxJ1*^-/-^, \**P* = 0.0129, t = 4.273, df = 4; Atp1a2, WT vs. *FoxJ1*^-/-^, \**P* = 0.0275, t = 3.391, df = 4; Ae2, WT vs. *FoxJ1*^-/-^, n.s. *P* = 0.6619, t = 0.4714, df = 4; unpaired nested two-tailed Student t-test**. Bottom**, the intensity of Aqp1, Atp1s2, and Ae2 across one cell line-scanned in Fig.1E. **G.** Quantitation for total Aqp1 and Atp1a2 in Fig. 1F. Total Aqp1, WT vs. *FoxJ1*^-/-^ ChP, \**P* = 0.0254, t = 3.477, df = 4; total Atp1a2, WT vs. *FoxJ1*^-/-^ ChP, \*\**P* = 0.0252, t = 3.487, df = 4; nested unpaired two-tailed Student t-test**. H.** FoxJ1 deficiency does not affect the apical expression of Nkcc1 in the choroid plexus. **Left,** representative images were shown for immunofluorescence staining on Nkcc1 in P7 WT and *FoxJ1*^-/-^ ChP paraffin sections. Line scan is for cellular fluorescence signals. **Right**, quantitation of relative fluorescence intensity and cellular location of Nkcc1 in P7 WT vs. *FoxJ1*^-/-^ ChP. Each dot represents the average intensity from 3-6 images in each biological sample. **I.** FoxJ1-dependent Ift88 deficiency impairs choroid plexus ciliogenesis. Ift88 deficiency decreases cilia number and ciliary length. P7 littermate control *Ift88 ^fl/fl^*and FoxJ1-Cre; *Ift88 ^fl/fl^* ChP explant were compared for cilia phenotype using immunostaining of Arl13b (red, ciliary axonemes), Ift88 (green) and DAPI (blue). **J**. *FoxJ1-Cre/+; Ift88^fl/fl^* mice exhibit a bulging forehead, consistent with the hydrocephalus phenotype. **K.** Ift88 deficiency in multiciliated cells increases the total expression of Aqp1 and Atp1a2. Three pairs of independent, littermate-controlled *Ift88^fl/fl^* and *FoxJ1-Cre/+; Ift88^fl/fl^*mice were compared. **Top**, zoom-out image of Aqp1, Atp1a2 and Ae2 in *Ift88^fl/fl^*and *FoxJ1-Cre/+; Ift88^fl/fl^* choroid plexus. **Middle**, quantitation of relative fluorescence intensity of Aqp1, Atp1a2, and Ae2 in P7 *Ift88^fl/fl^* vs. *FoxJ1-Cre/+; Ift88^fl/fl^*choroid plexus. Each dot is the average intensity from 3-6 images in one biological sample, n=3. Aqp1, *Ift88^fl/fl^* vs. *FoxJ1-Cre/+; Ift88^fl/fl^*, \**P* = 0.0275, t = 3.391, df = 4; Atp1a2, *Ift88^fl/fl^* vs. *FoxJ1-Cre/+; Ift88^fl/fl^*, \**P* = 0.0131, t = 4.254, df = 4; Ae2, WT vs. *FoxJ1*^-/-^, n.s. *P* = 0.1384, t = 1.847, df = 4; unpaired two-tailed Student t-test**. Bottom**, the intensity of Aqp1, Atp1s2, and Ae2 across one cell line-scanned in Fig.1H. **L.** Quantitation for total Aqp1 and Atp1a2 in Fig. 1I. Total Aqp1, *Ift88^fl/fl^* (n=5) vs. *FoxJ1-Cre/+; Ift88^fl/fl^* (n=4), \*\**P* = 0.0052, t = 4.001, df = 7; total Atp1a2, *Ift88^fl/fl^* (n=4) vs. *FoxJ1-Cre/+; Ift88^fl/fl^* (n=3), \**P* = 0.0225, t = 3.259, df = 5; nested unpaired two-tailed Student t-test.

**Supplementary Figure S2.**
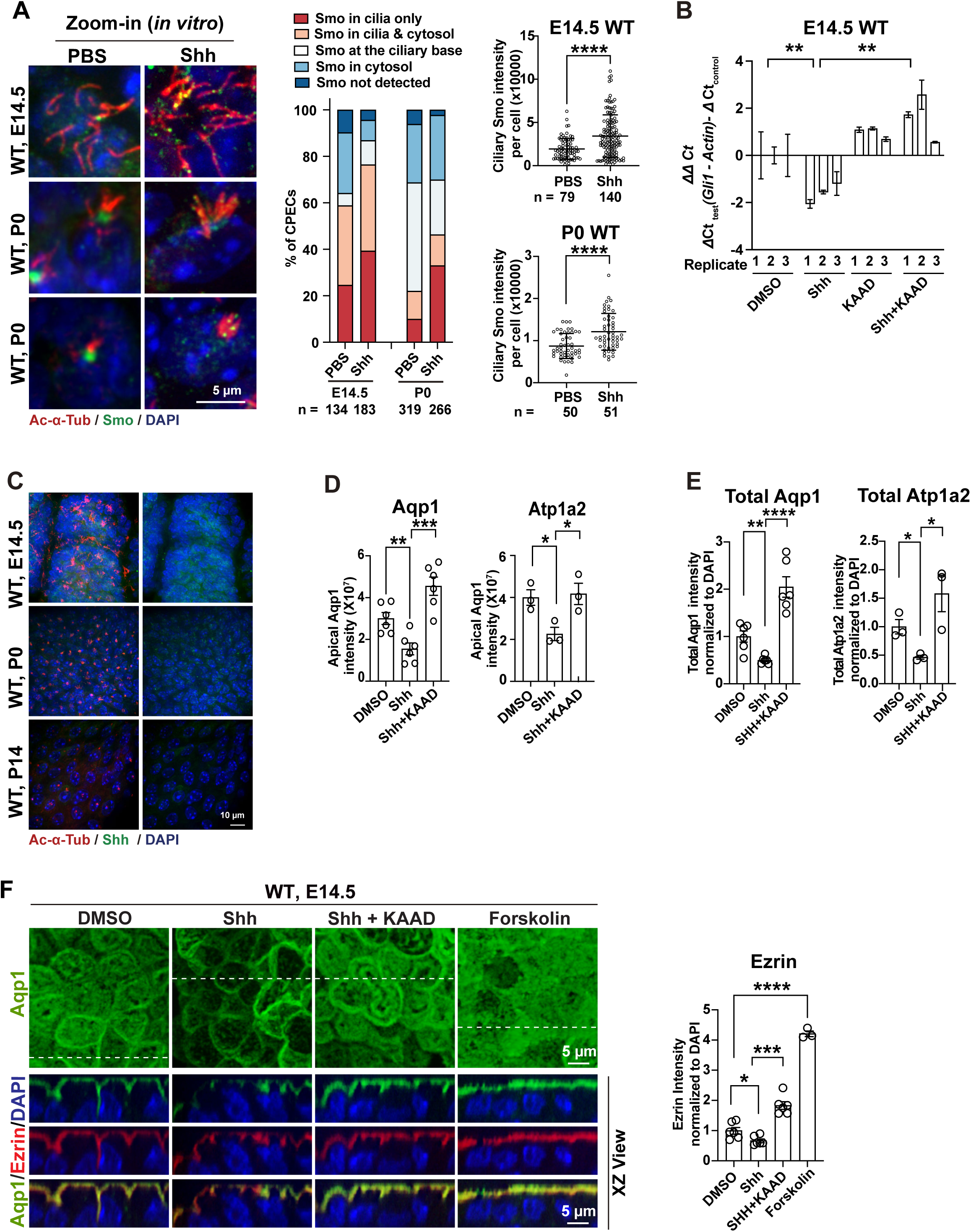
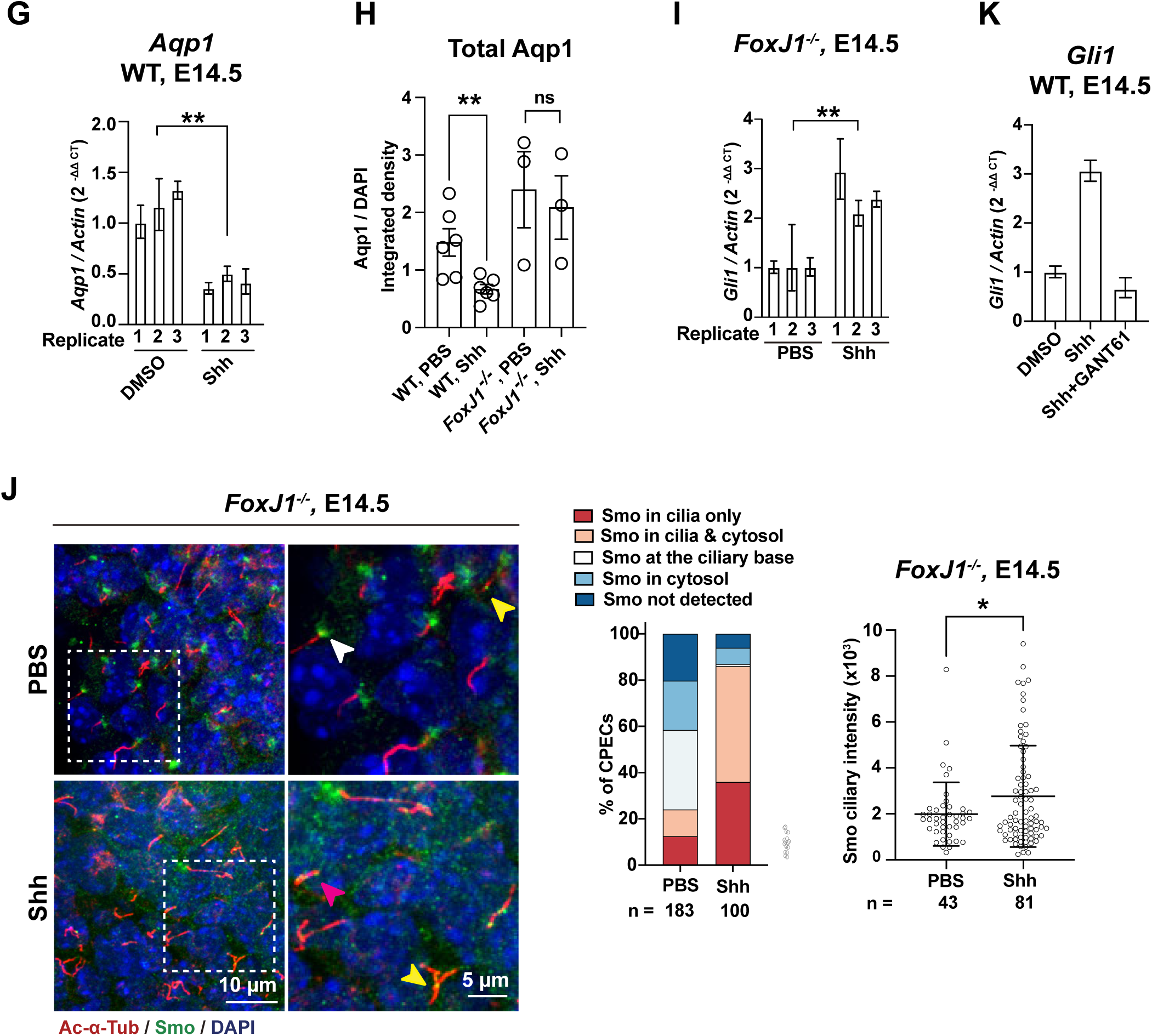
Choroid plexus cilia mediate Shh signaling to regulate the cAMP level and Aqp1 expression. **A.** Smo is translocated into choroid plexus cilia in response to Shh. E14.5 and P0 wildtype choroid plexus explants were treated with PBS or 10nM Shh (a Shh N-terminus protein) before being subjected to Smo immunostaining. Representative zoom-in images of Smo ciliary accumulation in response to Shh treatment in E14.5 and P0 WT (left), quantitation of Smo staining patterns (middle), and quantitation of ciliary Smo staining intensity per cell (right) were shown. E14.5 WT PBS vs. E14.5 WT Shh (n=3), \*\*\*\**P* < 0.0001, t = 5.064, df = 215; P0 WT PBS vs. P0 WT Shh (n=3), \*\*\*\**P* < 0.0001, t = 4.545, df = 99; unpaired two-tailed Student’s t-test. **B.** E14.5 WT Choroid plexus multicilia mediate the canonical Shh signaling, as demonstrated by Shh-dependent *Gli1* induction in real-time PCR analyses. *Gli1* mRNA was measured by real-time PCR analyses and plotted as ΔΔCT. ΔΔCT = ΔΔCT_test_ (*Gli1*-*Actin*) – ΔΔCT_control_(*Gli1*-*Actin*). **C.** Representative image of WT CPECs co-stained for Shh (green) and Ac-α-Tub (red) in E14.5, P0, and P14. **D.** Shh represses the apical expression of Aqp1 and Atp1a2 in the choroid plexus. Apical Aqp1 expression (n=6), DMSO vs. Shh, \*\**P* = 0.0048, t = 2.606, df = 10; Shh vs. Shh + KAAD, \*\*\**P* = 0.0002, t = 5.758, df = 10. Apical Atp1a2 expression (n=3), DMSO vs. Shh, \**P* = 0.0378, t = 3.055, df = 4; Shh vs. Shh + KAAD, \**P* = 0.0285, t = 3.353, df = 4; nested unpaired two-tailed Student t- test. **E.** Shh represses the total expression of Aqp1 and Atp1a2 in the choroid plexus. Total Aqp1 was measured in E14.5 WT CPECs; DMSO vs. Shh (n=6), \*\**P* = 0.0021, t = 4.106, df=10; Shh vs. Shh + KAAD (n=6), \*\*\*\**P* < 0.0001, t = 7.102, df = 10. Total Atp1a2 was measured in P0 CPECs; DMSO vs. Shh (n=3), \**P* = 0.0378, t = 3.055, df = 4; Shh vs. Shh + KAAD (n=3), \**P* = 0.0285, t = 3.353, df = 4; nested unpaired two-tailed Student t-test. **F.** cAMP stimulator forskolin increases Ezrin expression, whereas Shh inhibits Ezrin expression, which can be blocked by Smo inhibitor KAAD in E14.5 WT CPECs. **Left,** representative images were shown for immunofluorescence staining on Aqp1(green), Ezrin (red), and DAPI (blue). **Right**, quantitation on Ezrin was normalized to DAPI. Each dot is the average Ezrin/DAPI from 3-6 areas from one biological sample. DMSO (n=6) vs. Shh (n=6), \**P* = 0.0242, t = 2.652, df = 10; Shh (n=6) vs. Shh+ KAAD (n=6), \*\*\**P* < 0.001, t = 8.292, df = 10; DMSO (n=6) vs. forskolin (n=3), ****P < 0.0001, t = 18.85, df = 7; nested unpaired two-tailed Student t-test. **G.** Shh signaling activation increases *Aqp1 mRNA* in E14.5 WT CPECs. In response to treatment with Shh (a Shh N-terminus protein, 10nM). *Aqp1* mRNA was measured by real-time PCR analyses. For *Aqp1*, DMSO (n=3) vs. Shh (n=3), \*\**P* = 0.0019, t = 7.277, df = 4; unpaired two-tailed Student t-tests. **H.** *FoxJ1^-/-^* choroid plexus cilia fail to transduce Shh signaling to downregulate Aqp1. E14.5 wild type and *FoxJ1^-/-^* CPECs were treated by PBS or Shh (a Shh N-terminus protein, 10nM), and the total Aqp1 level was quantified using immunofluorescence staining. WT PBS vs. WT Shh (n=6), \*\**P* = 0.0092, t = 3.216, df = 10; *FoxJ1^-/-^* PBS vs. *FoxJ1^-/-^* Shh (n=3), n.s. *P* = 0.7370, t = 0.3601, df = 4; nested unpaired two-tailed Student t-tests. **I.** *FoxJ1* deficiency does not abolish the ability of choroid plexus cilia to transduce canonical Shh signaling. In response to treatment with Shh (a Shh N-terminus protein, 10nM), *Gli1* was induced in E14.5 *FoxJ1^-/-^* CPECs as measured by real-time PCR analyses. DMSO vs. Shh (n=3), \*\**P* = 0.0041, t = 5.919, df = 4; unpaired two-tailed Student t-tests. **J**. Smo is translated into *FoxJ1^-/-^* choroid plexus cilia in response to Shh signaling. E14.5 CPECs were treated by PBS or Shh (a Shh N-terminus protein, 10nM), before being subjected to Smo immunostaining. Representative images (left), quantitation of Smo staining patterns (middle), and quantitation of ciliary Smo staining intensity (right) were shown. *FoxJ1^-/-^* PBS vs. *FoxJ1^-/-^* Shh (n=3): \**P* = 0.0389, t = 0.2088, df = 122; unpaired two-tailed Student t- tests. **K.** GANT61 acts as a potent Gli inhibitor in CPECs. E14.5 wildtype CPECs were treated by DMSO, Shh (a Shh N-terminus protein, 10nM) or GANT61 (25μM) + Shh (10nM), and Gli1 expression was measured by real-time PCR analyses. GANT61 efficiently abolished Shh-induced *Gli1* induction in the choroid plexus.

**Supplementary Figure S3.**
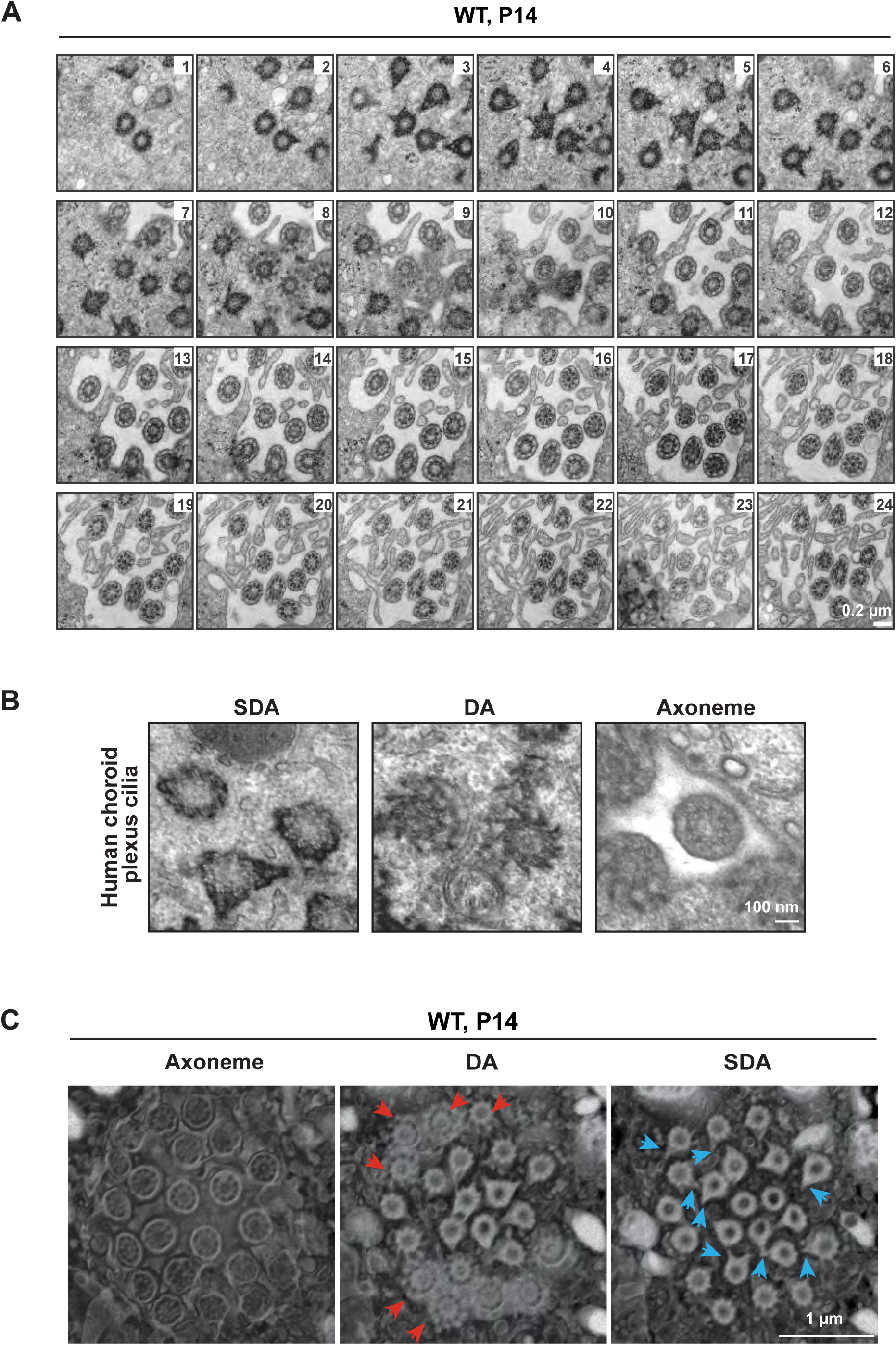
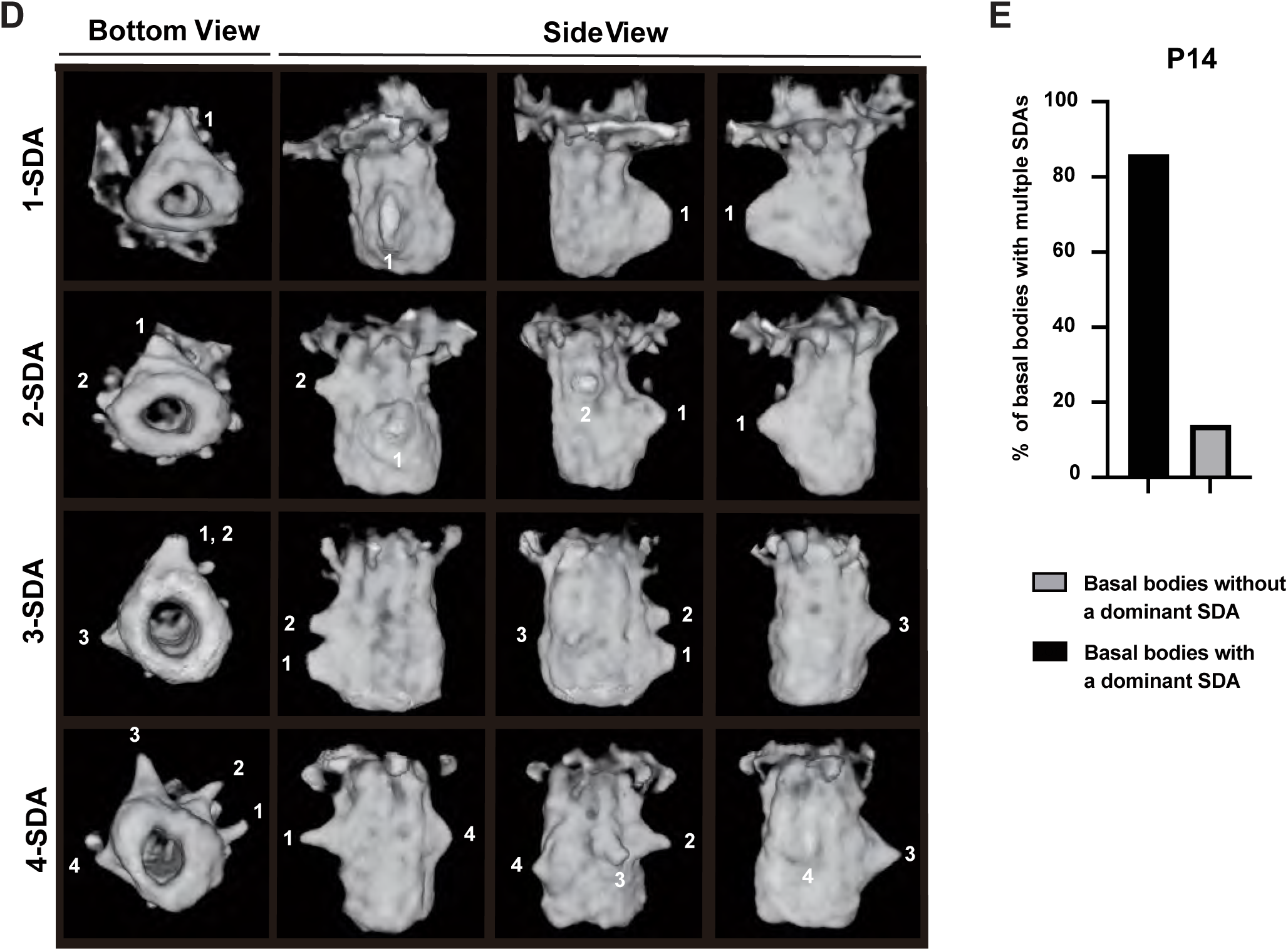
Mature choroid plexus multicilia exhibit distinct ciliary ultrastructure. **A**. 24 serial TEM images (50 nm/section) on mouse P14 choroid plexus reveal distinct basal body ultrastructure along the proximal to distal axis. These cilia are characterized by nine symmetrically arranged DAs, one or multiple SDAs, and a “9+0” axoneme microtubule arrangement. The number on each image indicates the sequence of the image in the consecutive TEM series. Scale bar, 0.2μm. **B.** Human choroid plexus cilia resemble mouse choroid plexus cilia in ultrastructure, characterized by a “9+0” microtubule configuration, 9 radially arranged DAs, and basal bodies with multiple asymmetrically arranged SDAs. Scale bar, 100 nm. **C.** On-Face views of 3D rendering of choroid plexus ciliary axoneme, DAs (red arrows), and SDAs (blue arrows) using FIB-SEM data. We used a customized enhanced platform throughout a volume of 60 x 60 x 60 µm with 8 x 8 x 8 nm^3^ isotropic voxel sampling to analyze 17 P14 choroid plexus epithelial cells (CPECs) and obtained 286 imaged basal bodies (∼87% of all captured basal bodies). Raw file physical size, 4.536 µm x 3.928 µm x 3.2 µm. The DA and SDA views are clipped (0.216 µm and 0.288 µm) relative to the axoneme volume rendering. Scale bar, 1μm. **D.** Representative 3D rendering of mouse P14 choroid plexus basal bodies with 1, 2, 3, or 4 SDAs, viewed from four different angles (bottom view and 3 side view angels). Numbers indicate different SDAs of the same basal body. **E.** Choroid plexus basal bodies with multiple SDAs often contain a dominant SDA. Using FIB-SEM data, we analyzed 229 imaged basal bodies from 13 mouse P14 CPECs. ∼85% of basal bodies with multiple SDAs carry a dominant SDA in size.

**Supplementary Figure S4.**
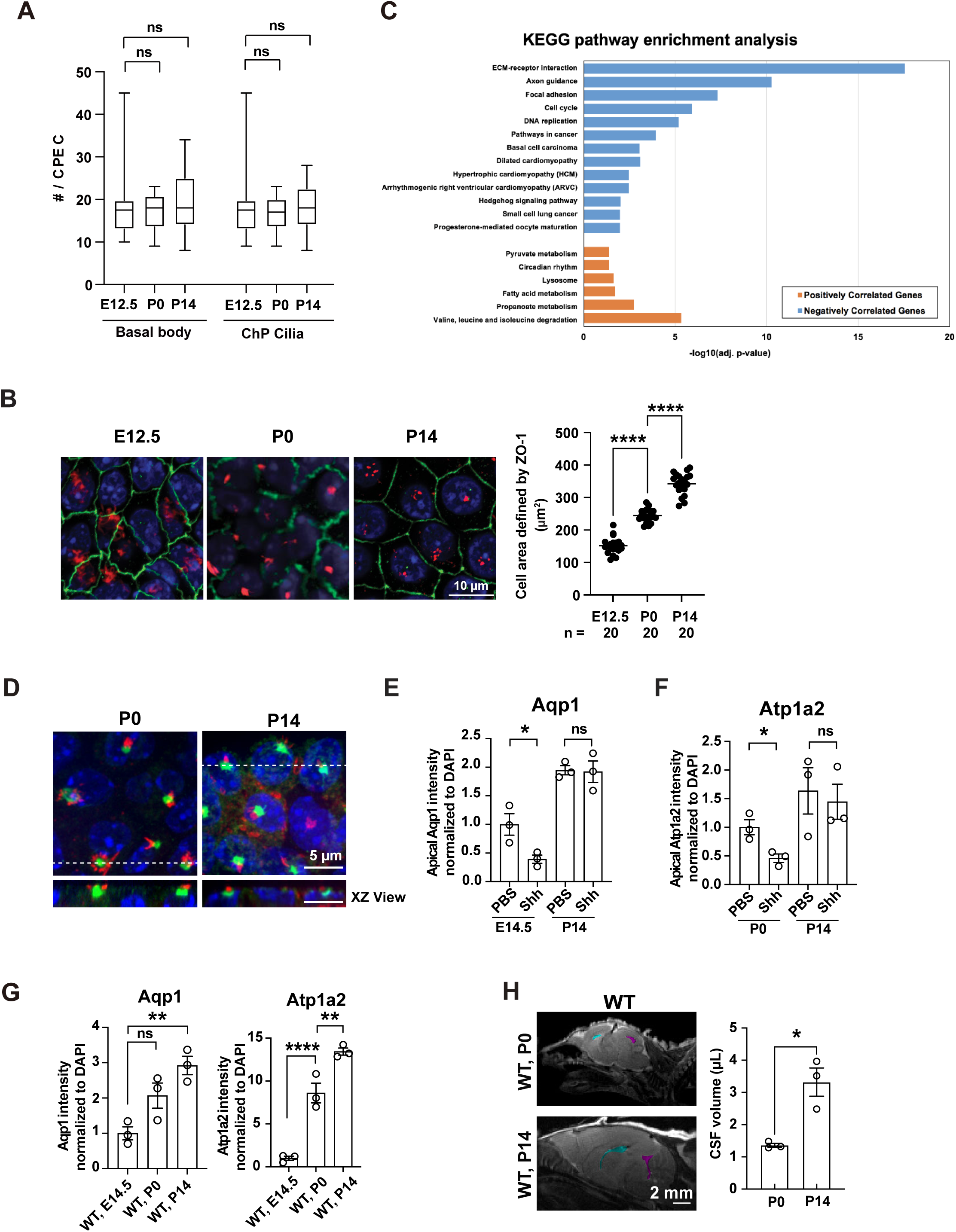
A dynamic change in ciliary length attenuates choroid plexus Shh signaling in development. **A**. The number of cilia and basal bodies per CPEC remains constant during choroid plexus development. FIB-SEM analyses demonstrate that an average CPEC harbors 17-19 cilia or basal bodies per cell in E14.5, P0, and P14 choroid plexus. n.s.not significant; unpaired two-tailed Student’s t-test. **B.** CPECs increase in size during normal development. The cell size of E12.5, P0, and P14 choroid plexus was quantified using cell areas defined by ZO-1 immunostaining. E12.5 vs. P0 (n=20), \*\*\*\**P* < 0.0001, t = 10.96, df = 57; P0 vs. P14 (n=20), \*\*\**P* = 0.0004, t = 11.59, df = 57, one-way ANOVA and Sidak multiple comparisons tests. **C.** Developmentally downregulated genes in choroid plexus are enriched for Shh pathway components. The transcriptomes of E12.5, P0, P14, and 6-month-old choroid plexus were compared using RNA-seq, and the genes with a significant developmental increase (orange) or decline (blue) were subjected to KEGG pathway enrichment analyses. **D.** XY and XZ view of Smo location in P0 and P14 WT ChP from Fig. 4F. **E, F.** Developing choroid plexus attenuates the Shh inhibition on Aqp1 and Atp1a2 expression. E14.5, P0, and P14 choroid plexus explants were treated with Shh (a Shh N-terminus protein, 10nM) for 24 hours before apical Aqp1 or Atp1a2 levels were quantified using immunofluorescence. **E.** Apical Aqp1: E14.5 PBS vs. E14.5 Shh, \**P* = 0.0438, t = 2.908, df = 4; P14 PBS vs. P14 Shh, n.s. *P* = 0.9087, t = 0.1221, df = 4. **F.** Apical Atp1a2 (P0 and P14), P0 PBS vs. P0 Shh (n=3), \**P* = 0.0497, t = 2.781, df = 4; P14 PBS vs. P14 Shh (n=3), n.s. *P* = 0.6788, t = 0.4457, df = 4; nested unpaired two-tailed Student’s t-test. **G.** Aqp1 and Atp1a2 exhibit an increased expression during choroid plexus development. E14.5, P0, and P14 choroid plexus were subjected to immunofluorescence staining of Aqp1 and Atp1a2, and the total fluorescence signals were quantified. Total Aqp1: E14.5 vs. P0 (n=3), n.s. *P* = 0.0993, t = 2.728, df = 6; E14.5 vs. P14 (n=3), \*\**P* = 0.0081, t = 4.905, df = 6. Total Atp1a2: E14.5 vs. P0 (n=3), \*\*\**P* = 0.0006, t = 7.480, df = 6; P0 vs. P14 (n=3), \*\**P* = 0.0064, t = 4.743, df = 6. nested one-way ANOVA and Sidak multiple comparisons tests. **H.** MRI measurement reveals a CSF volume increase during postnatal development. Representative sagittal MRI scans (left) and quantitative measurement of CSF volume (right) were shown for P0 and P14 wild-type mice. P0 vs. P14 (n=3), \**P* =0.0115, t = 4.426, df = 4; unpaired two-tailed Student’s t-test. Error bars, sem. Pseudocolor indicates the lateral (cyan) and the 4th (magenta) ventricle.

**Table S1.** Ultrastructural comparison among different cilia.

|  | **Primary cilia** | **Motile cilia<sup>a</sup>** | **Choroid plexus cilia** |
| --- | --- | --- | --- |
| **Cell type** | nearly all cell types | airway epithelial cells,<br>ependymal epithelial<br>cells, fallopian tube<br>epithelial cells, sperm | CPECs |
| **Cilia number / cell** | 1 per cell | hundreds per cell or<br>1 per cell (sperm) | 8-28 cilia per cell<br>(average ~18 cilia / CPEC) |
| **Axoneme**<br>**microtubule**<br>**configuration** | 9+0 | 9+2 | 9+0 |
| **DA # / basal body** | 9 / basal body | 9 / basal body | 9 / basal body |
| **SDA # / basal**<br>**body** | 9 / basal body | 1 / basal body | 1-9 / basal body |
| **SDA arrangement**<br>**on basal body** | symmetrical<br>arrangement |  | random arrangement |
| **Ciliary attachment** | daughter centriole | rootlet | n.d. |
| **Basal foot**<br>**directionality** | N/A | yes | no (1SDA-basal body) |

n.d.: not detected

**Table S2.**
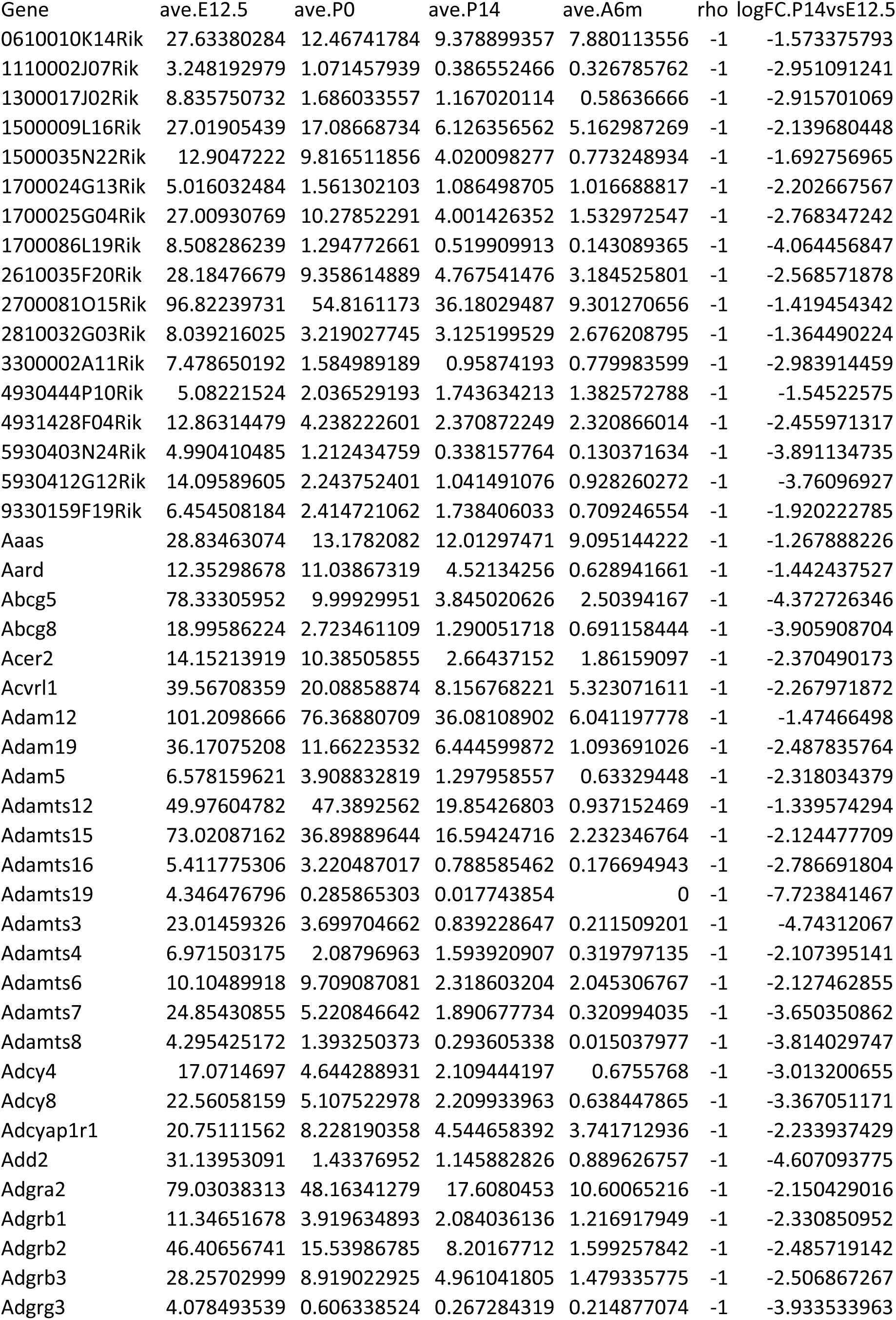
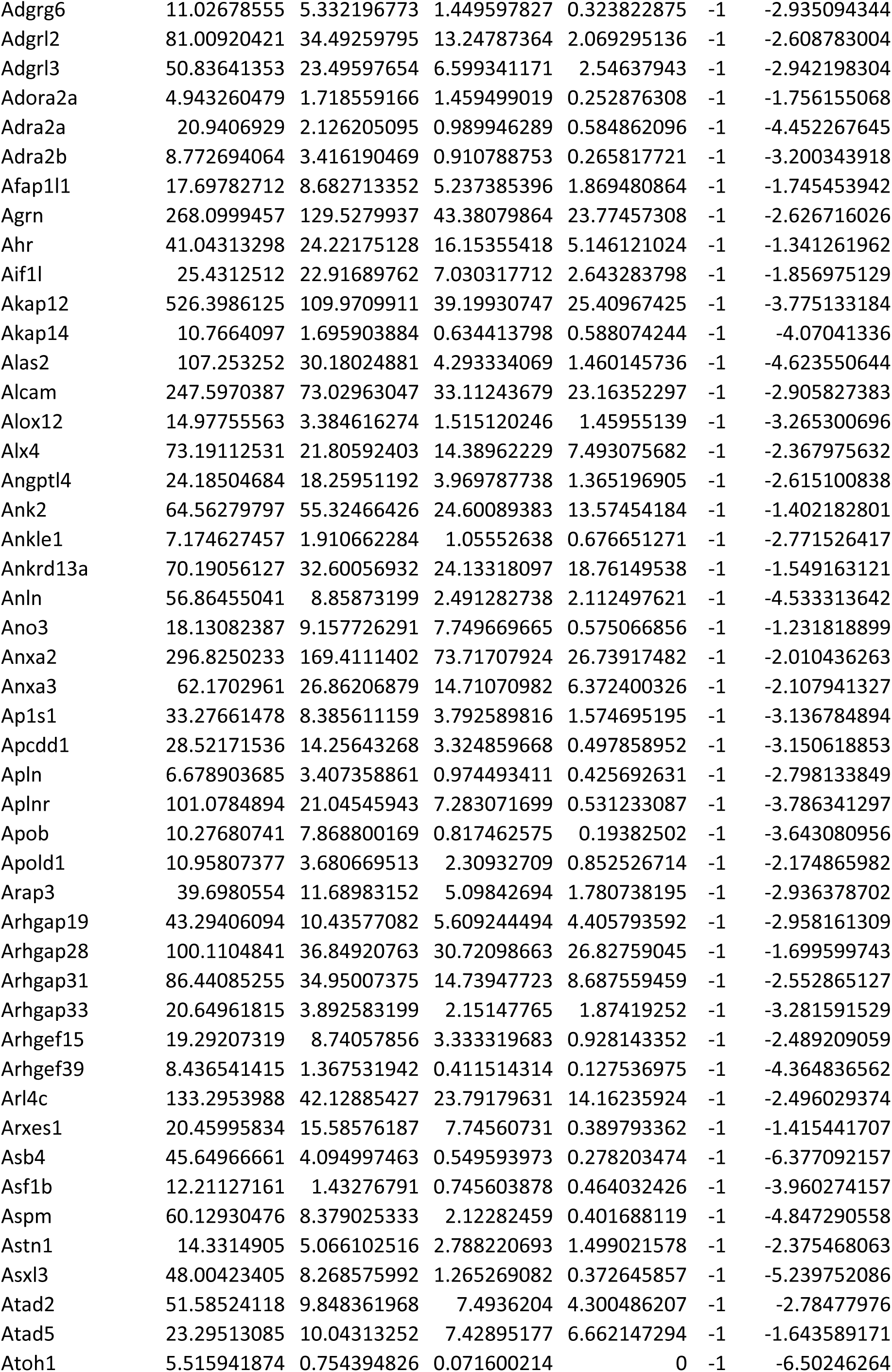
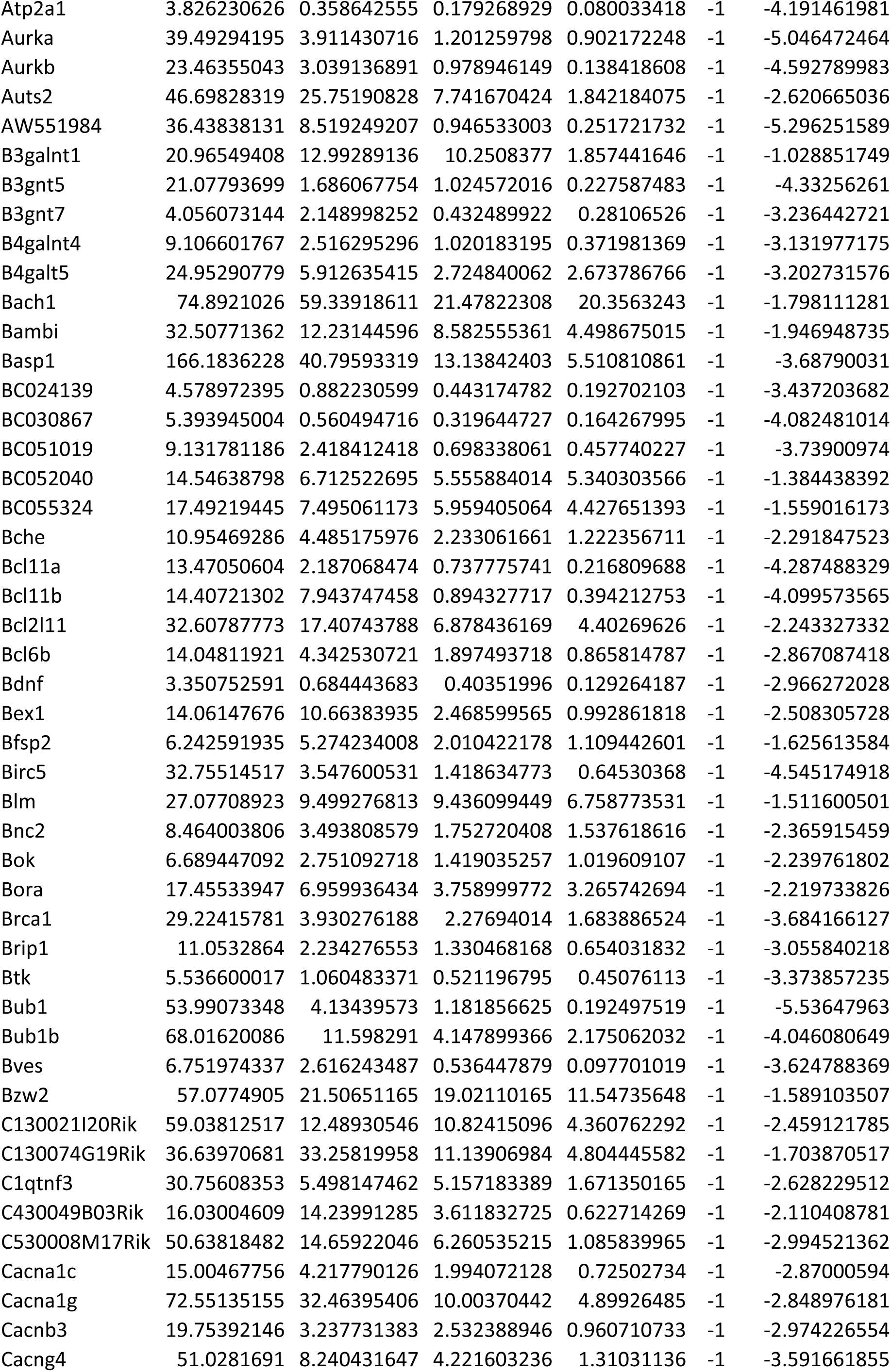
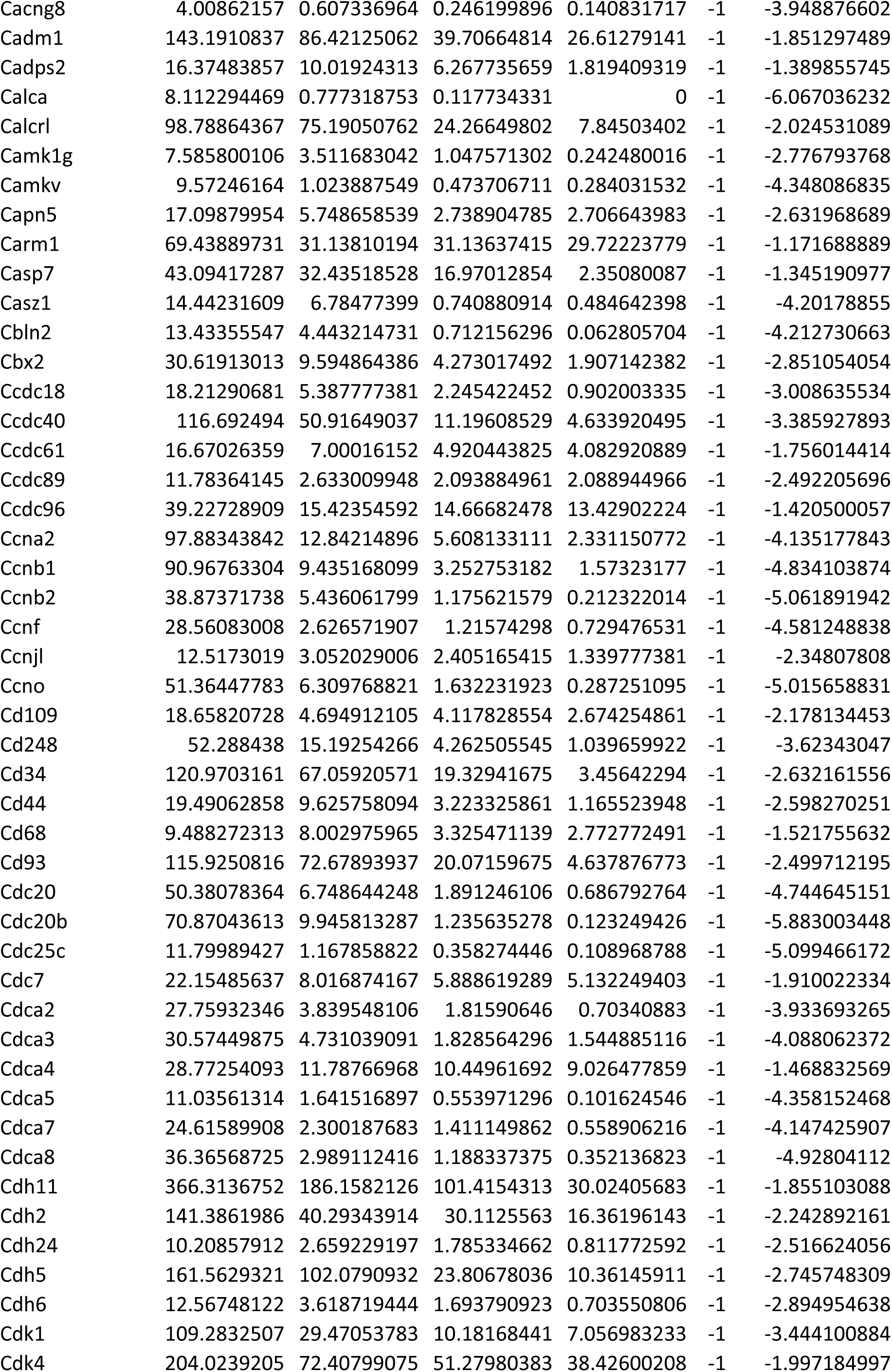
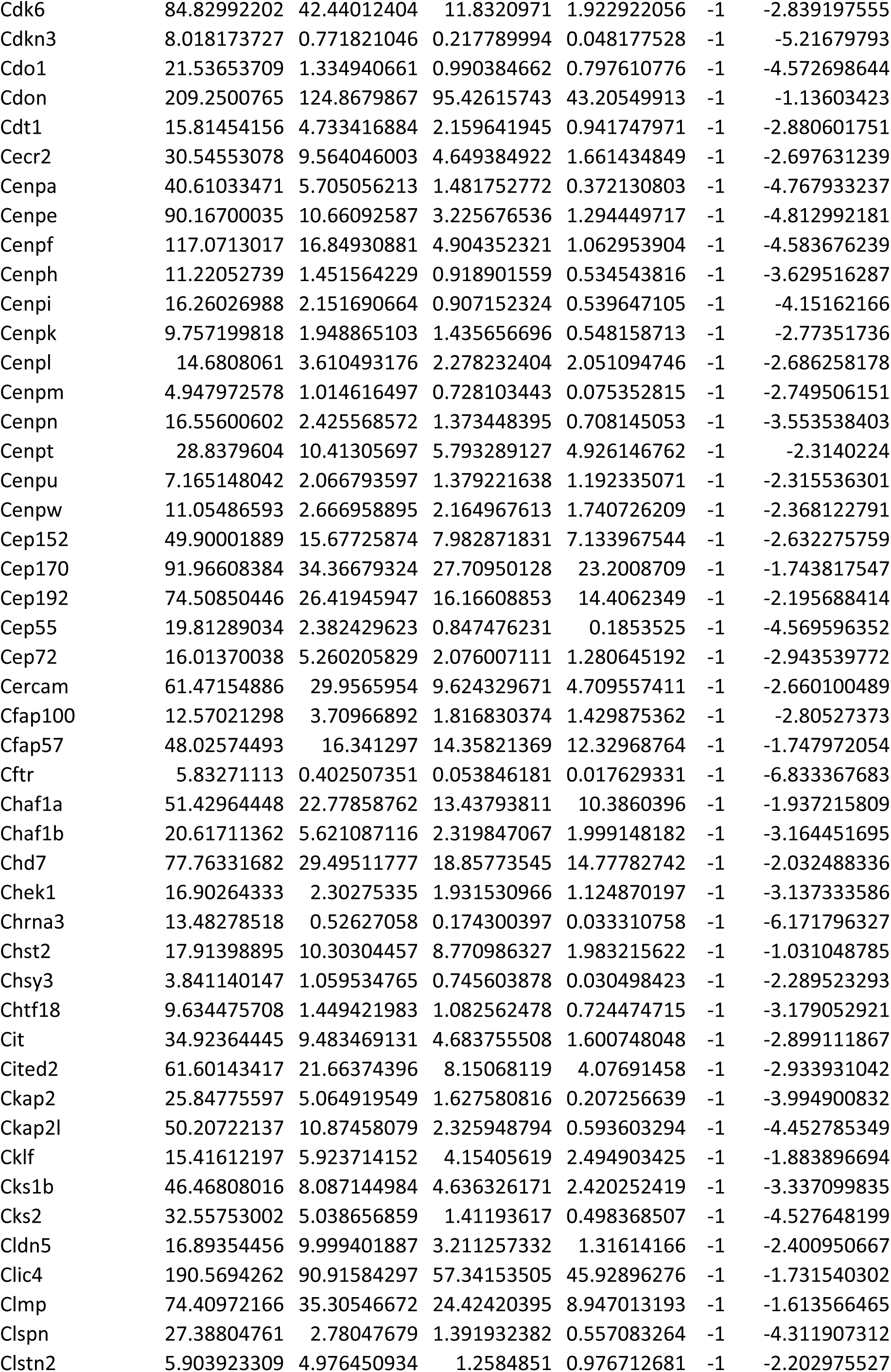
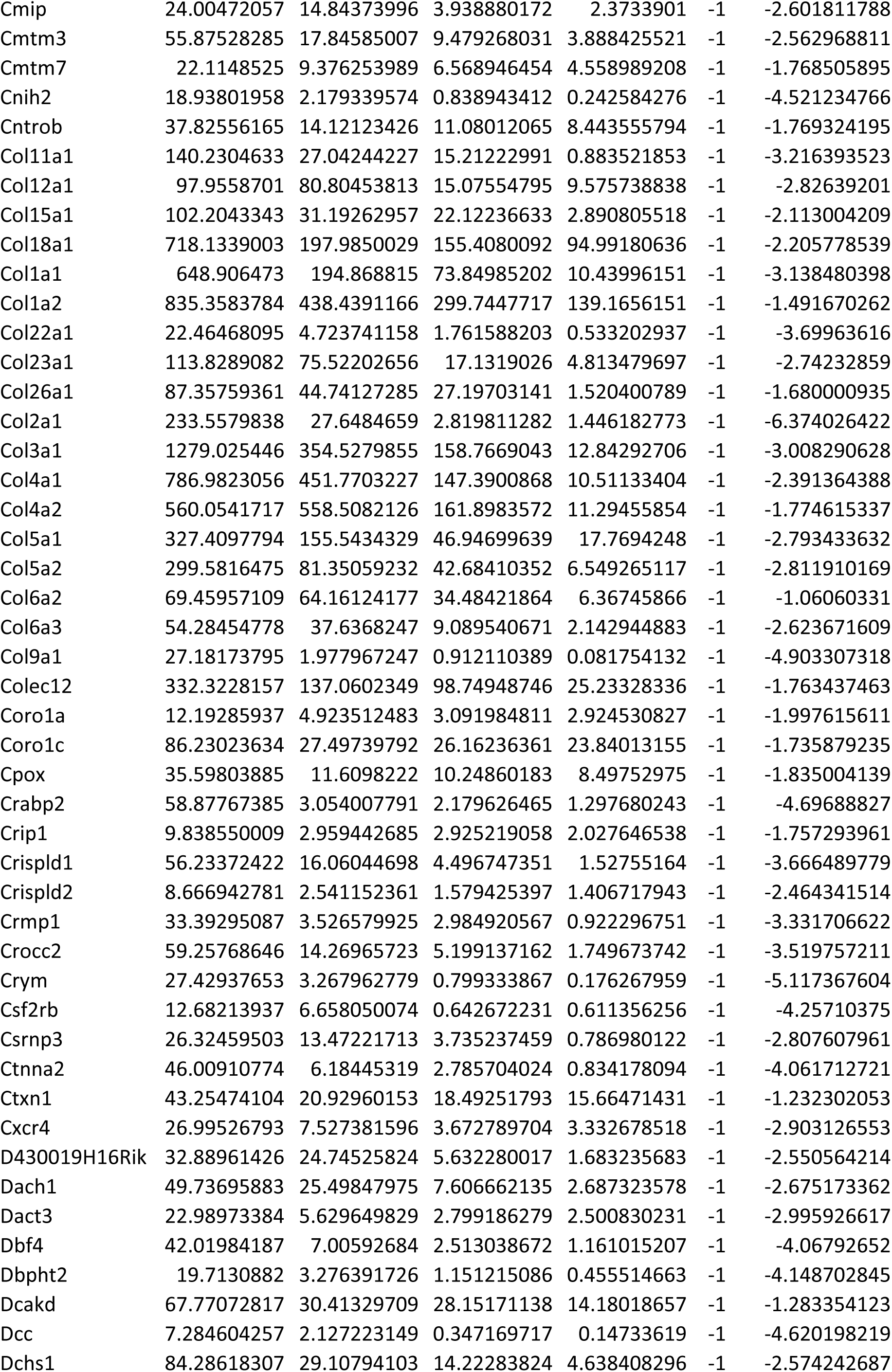
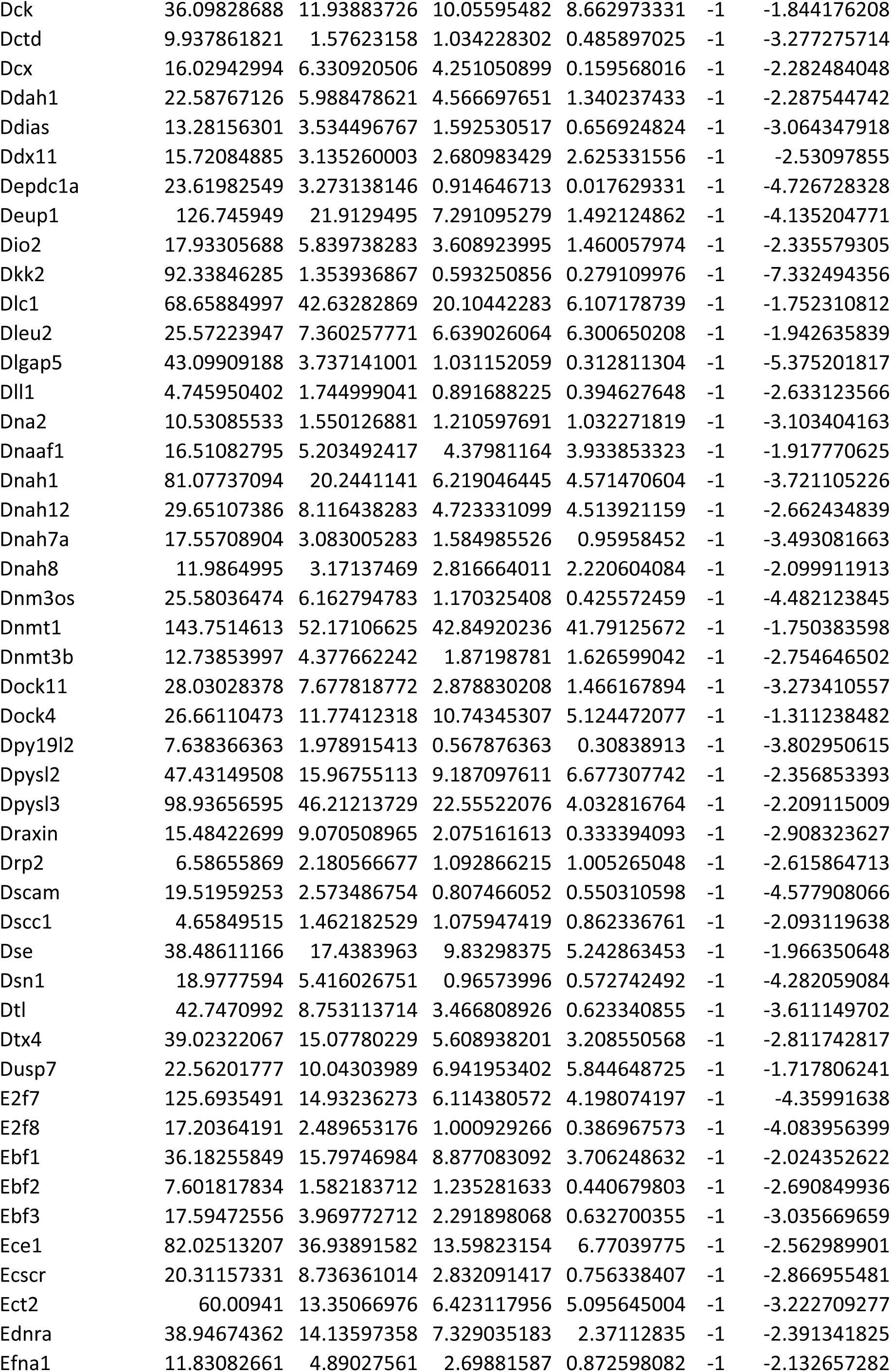
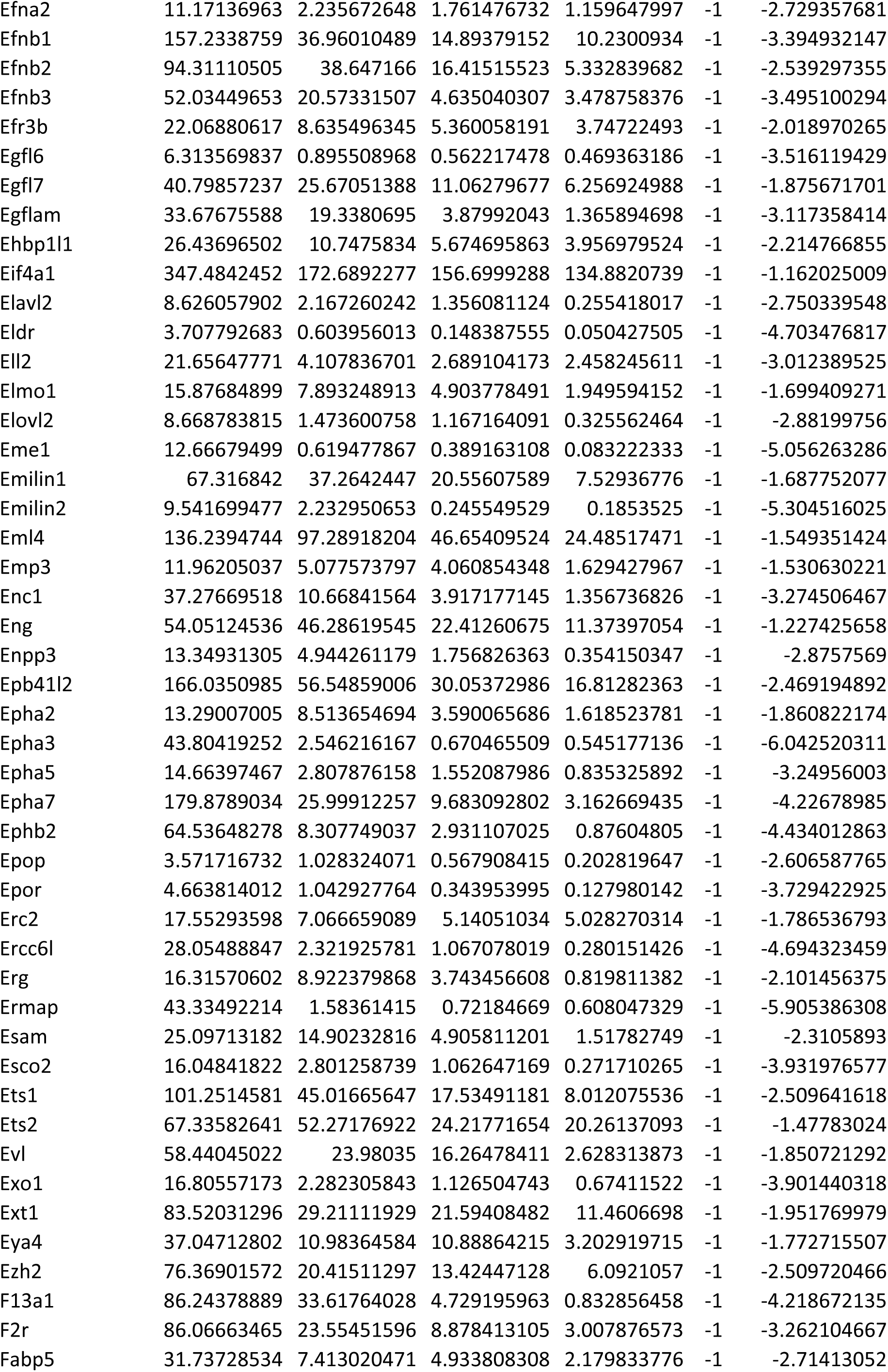
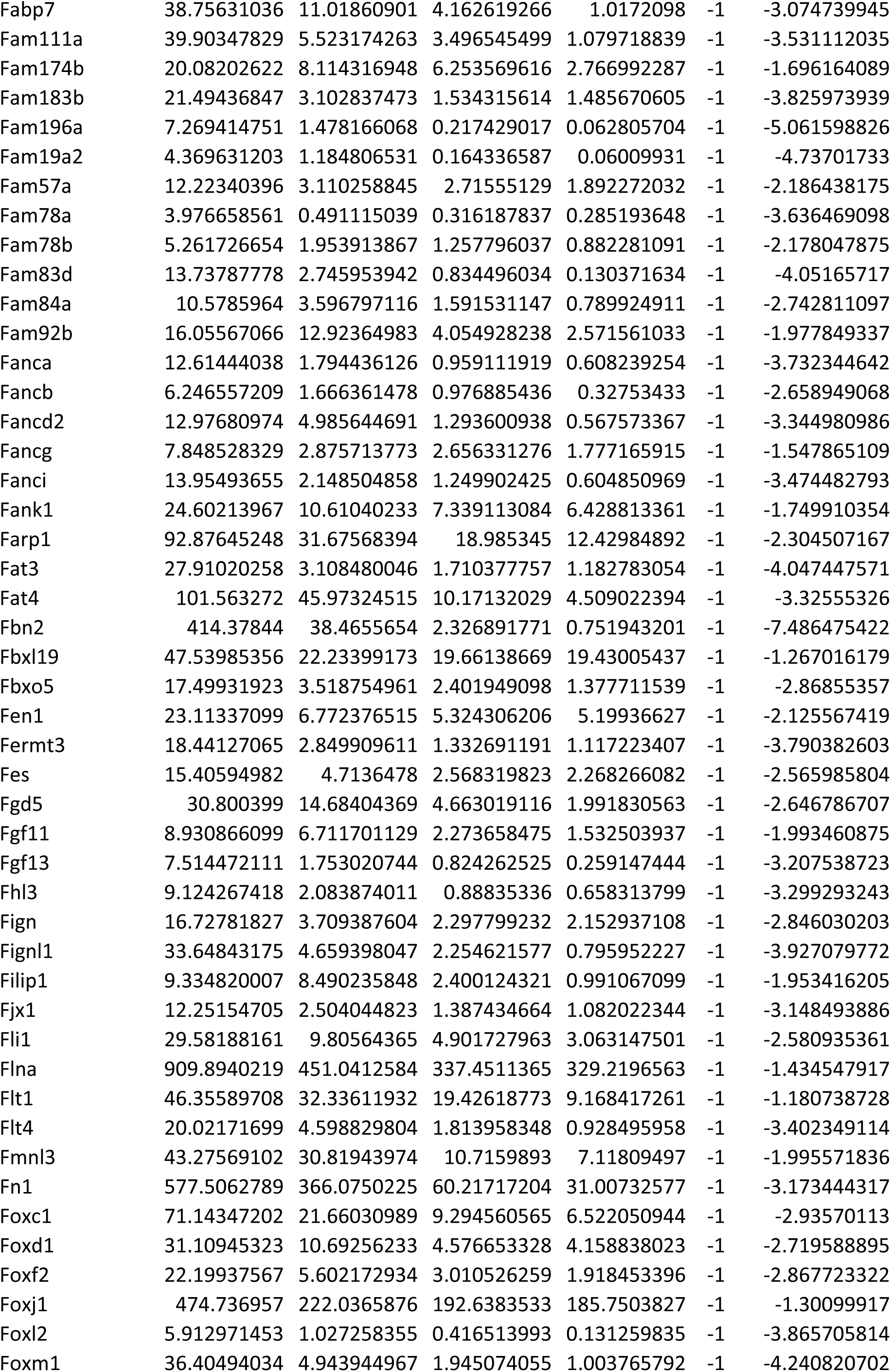
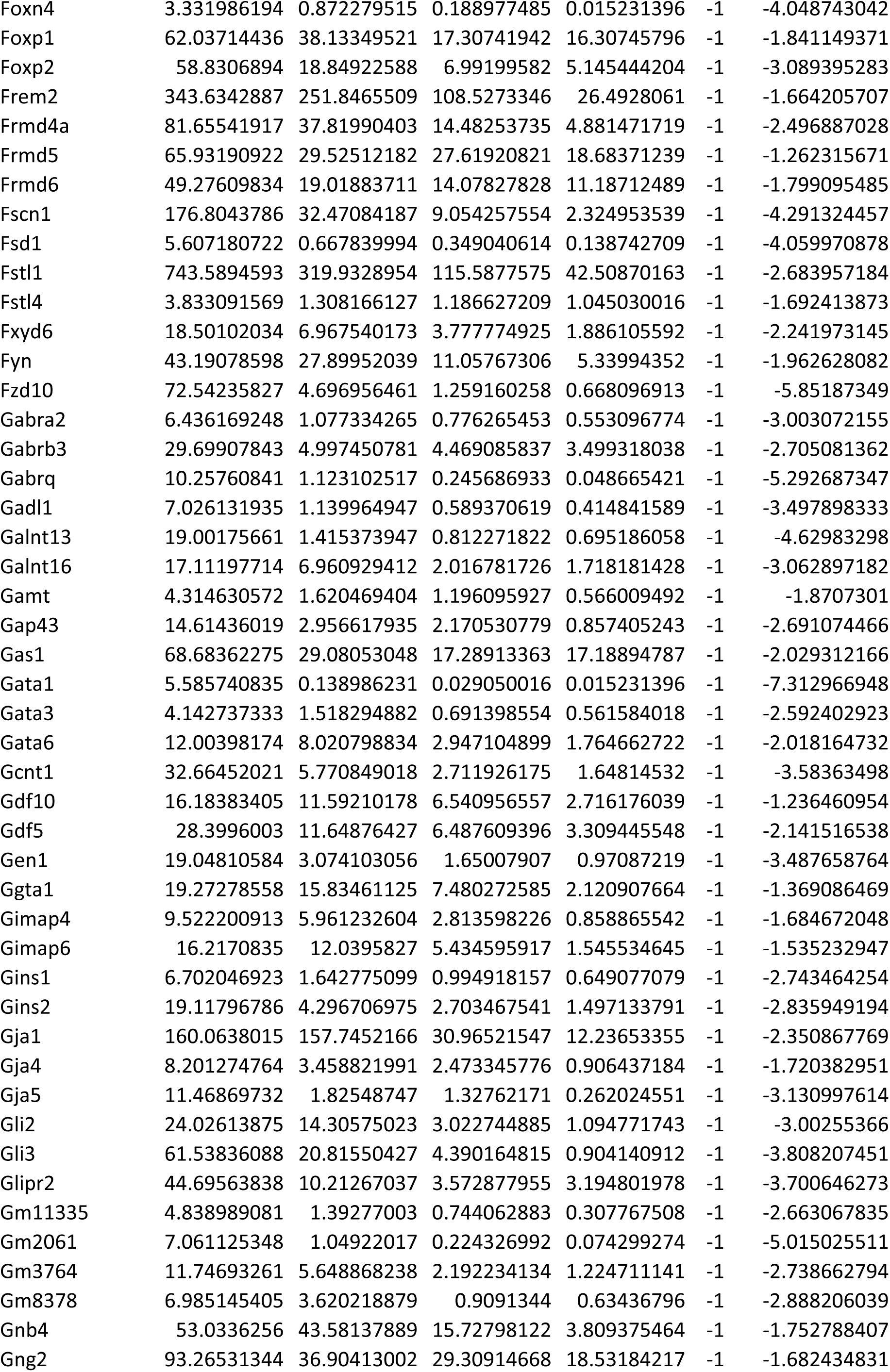
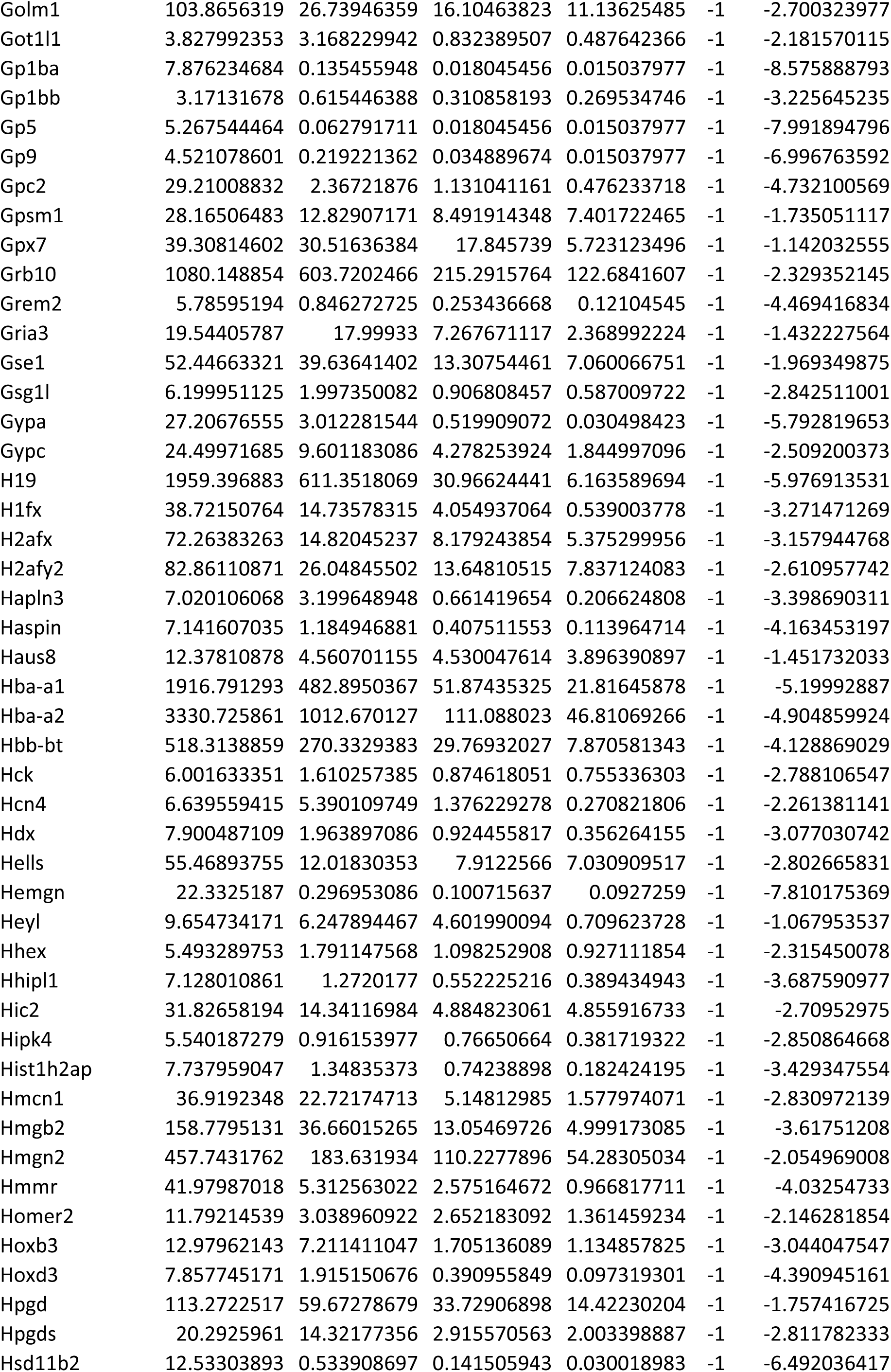
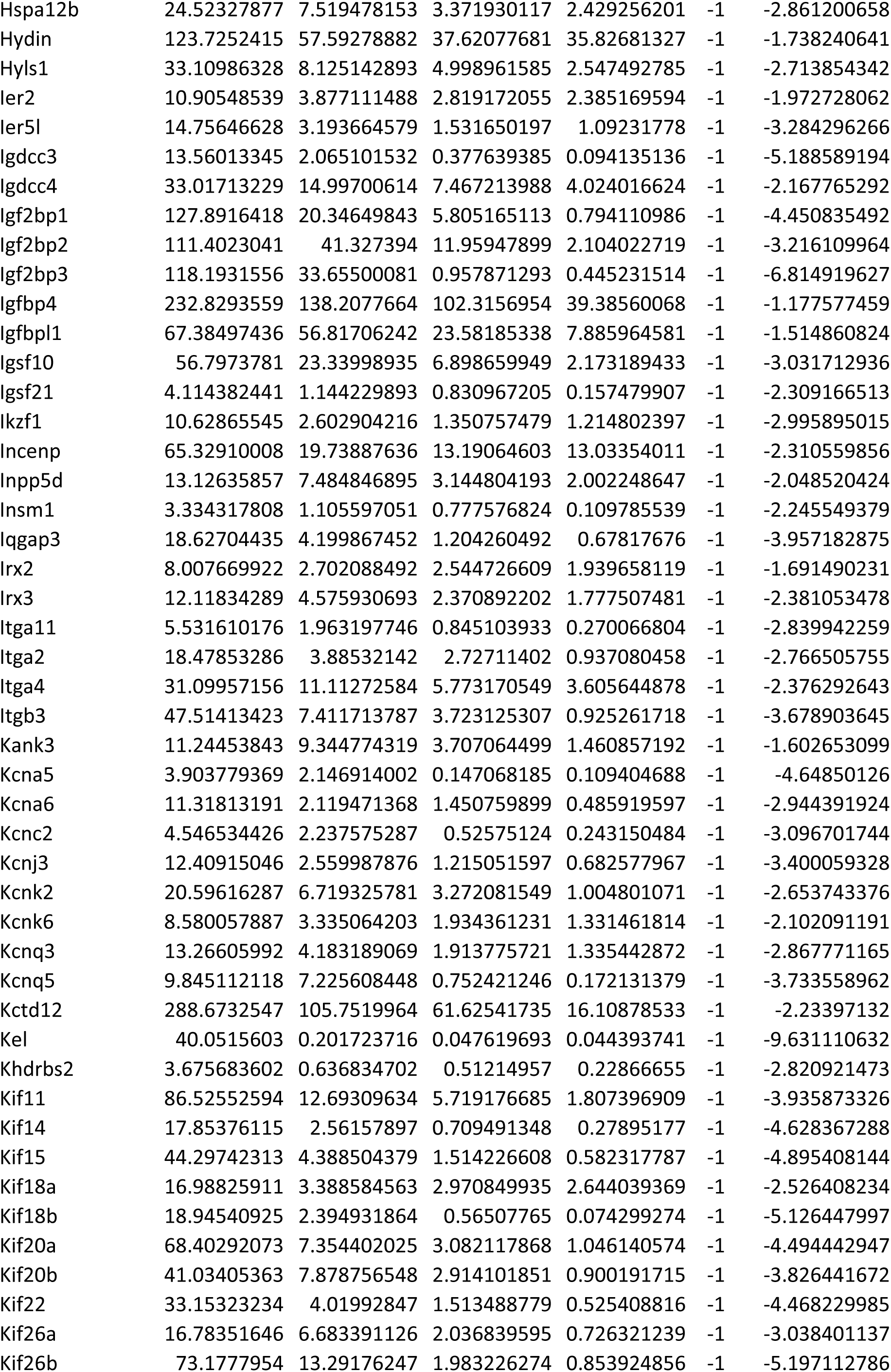
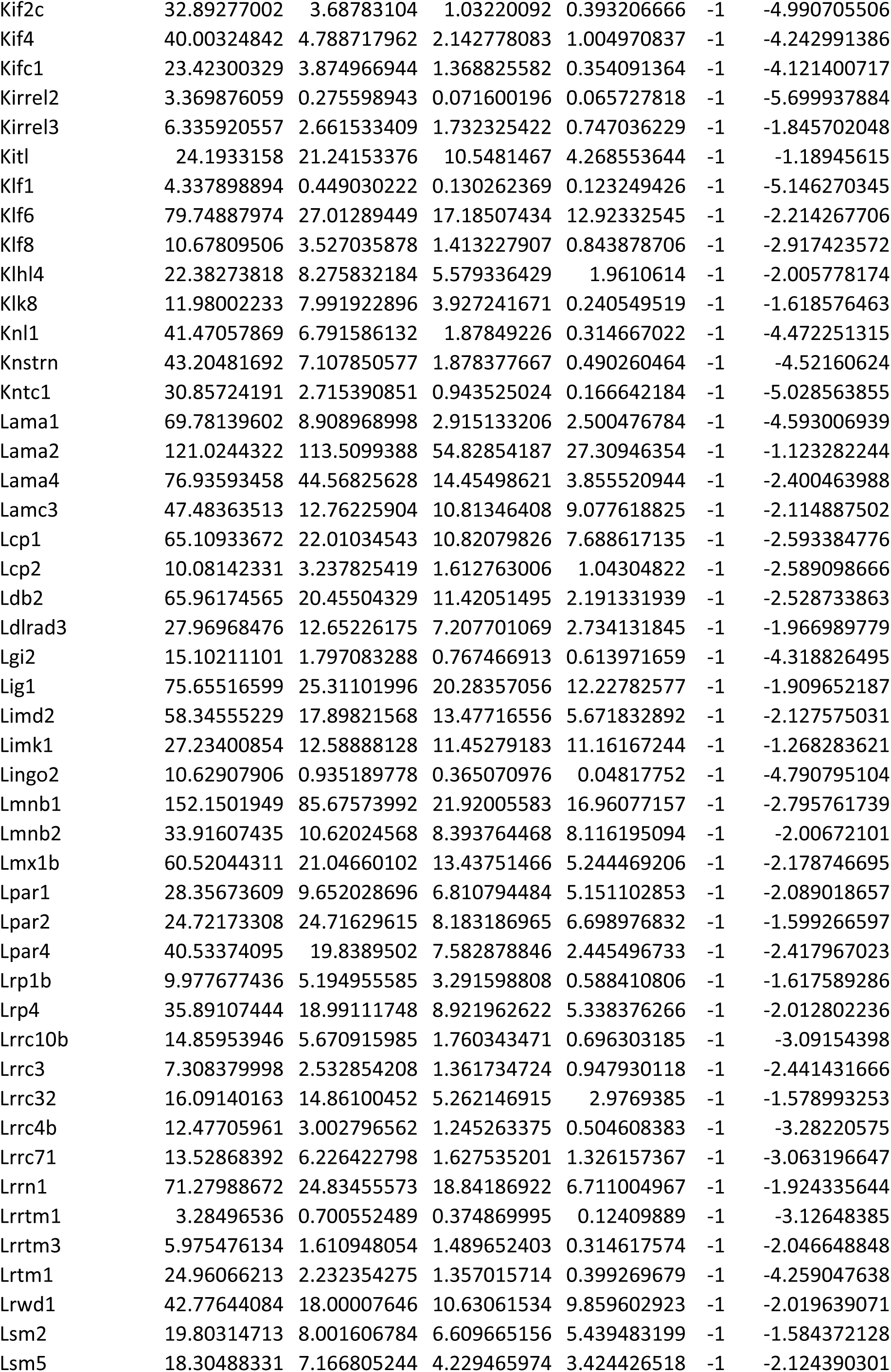
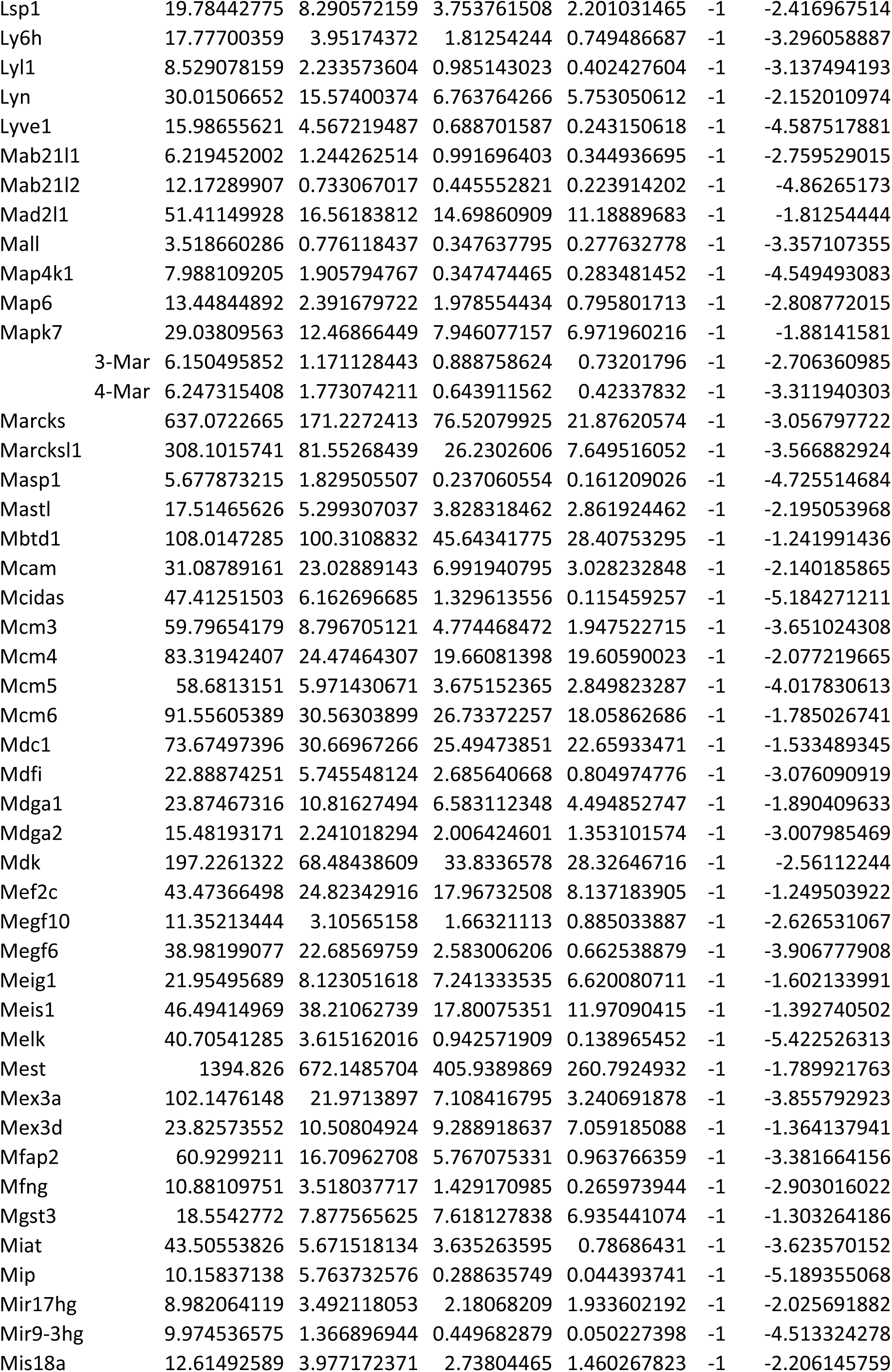
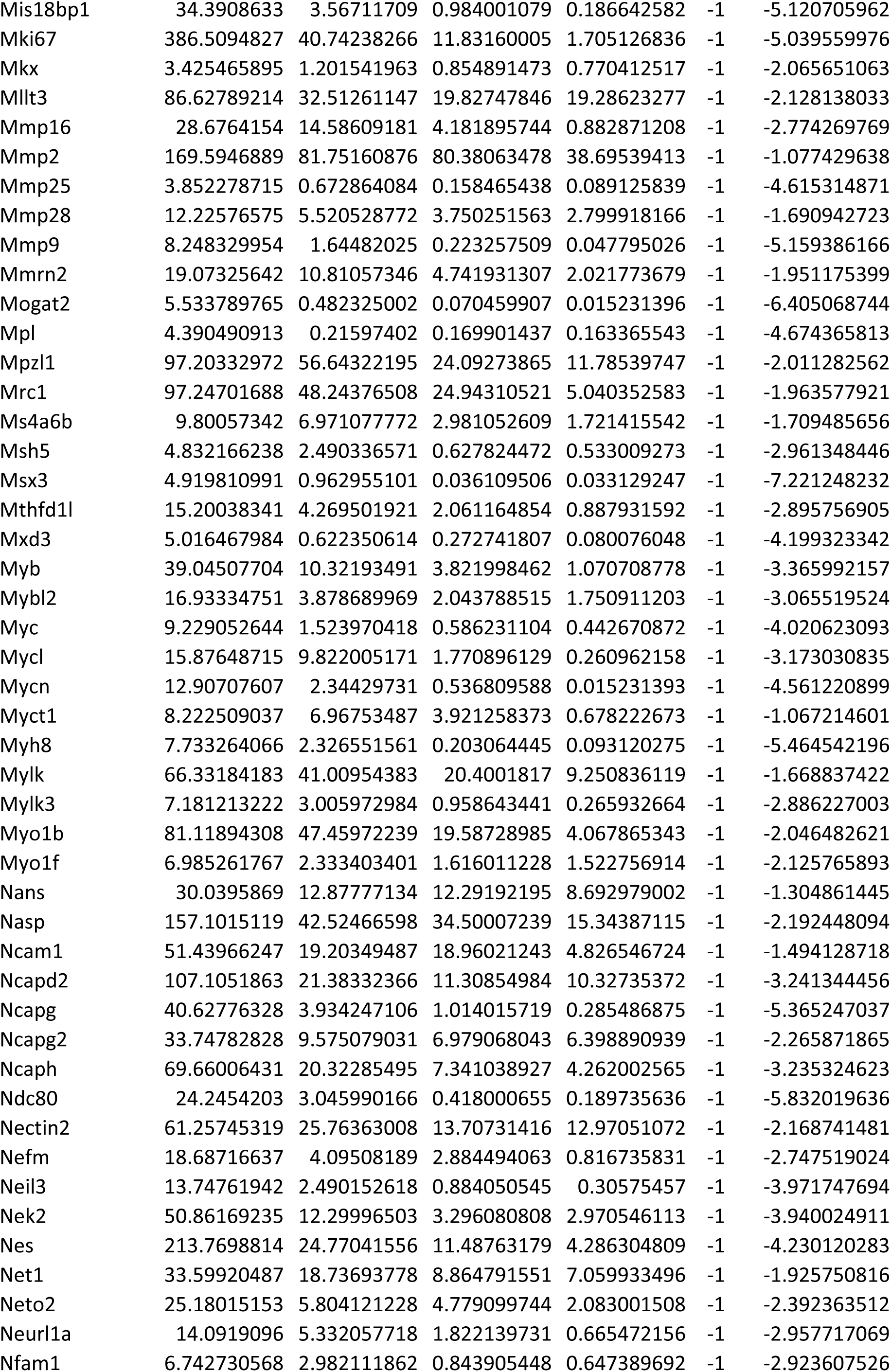
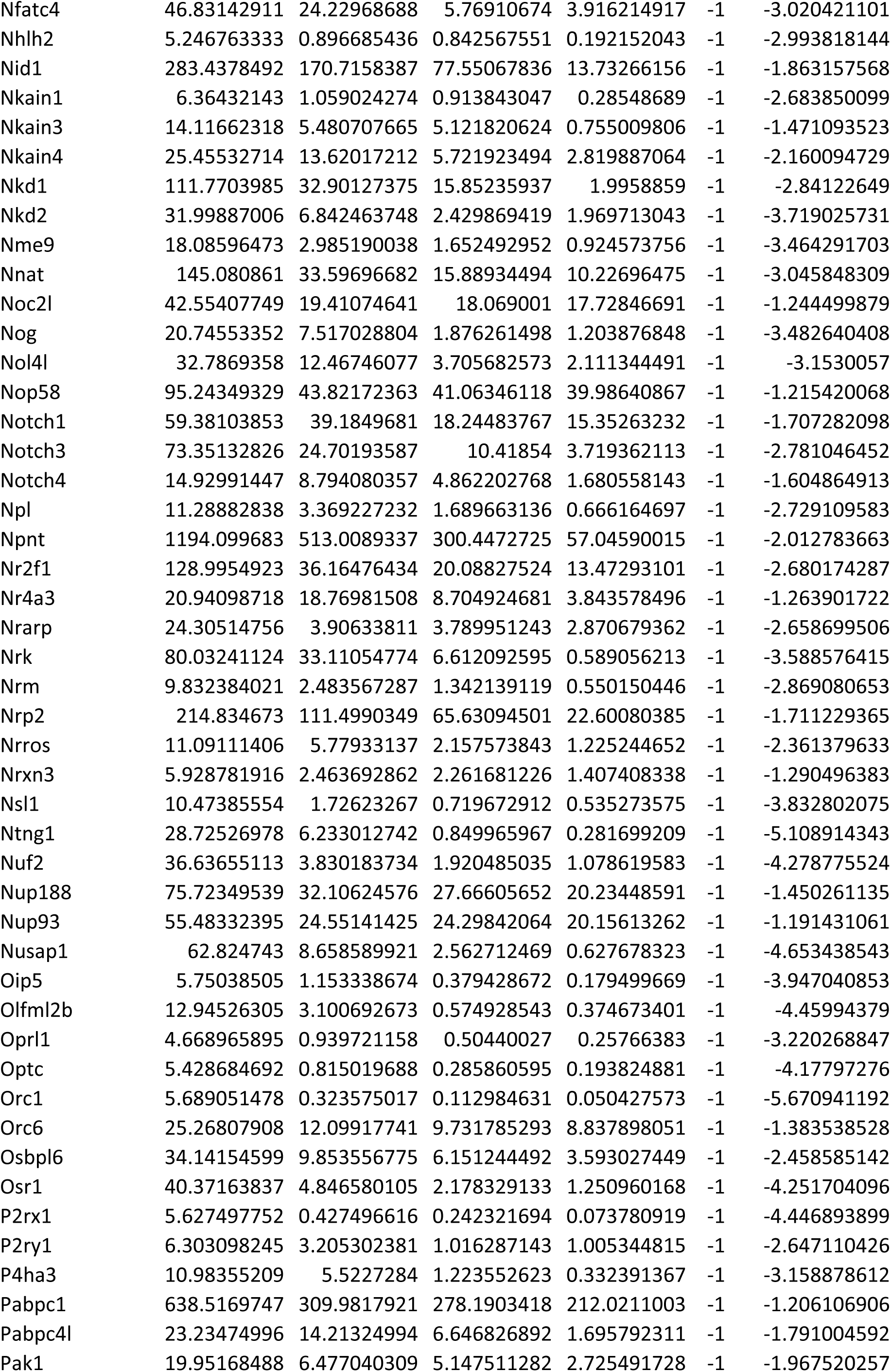
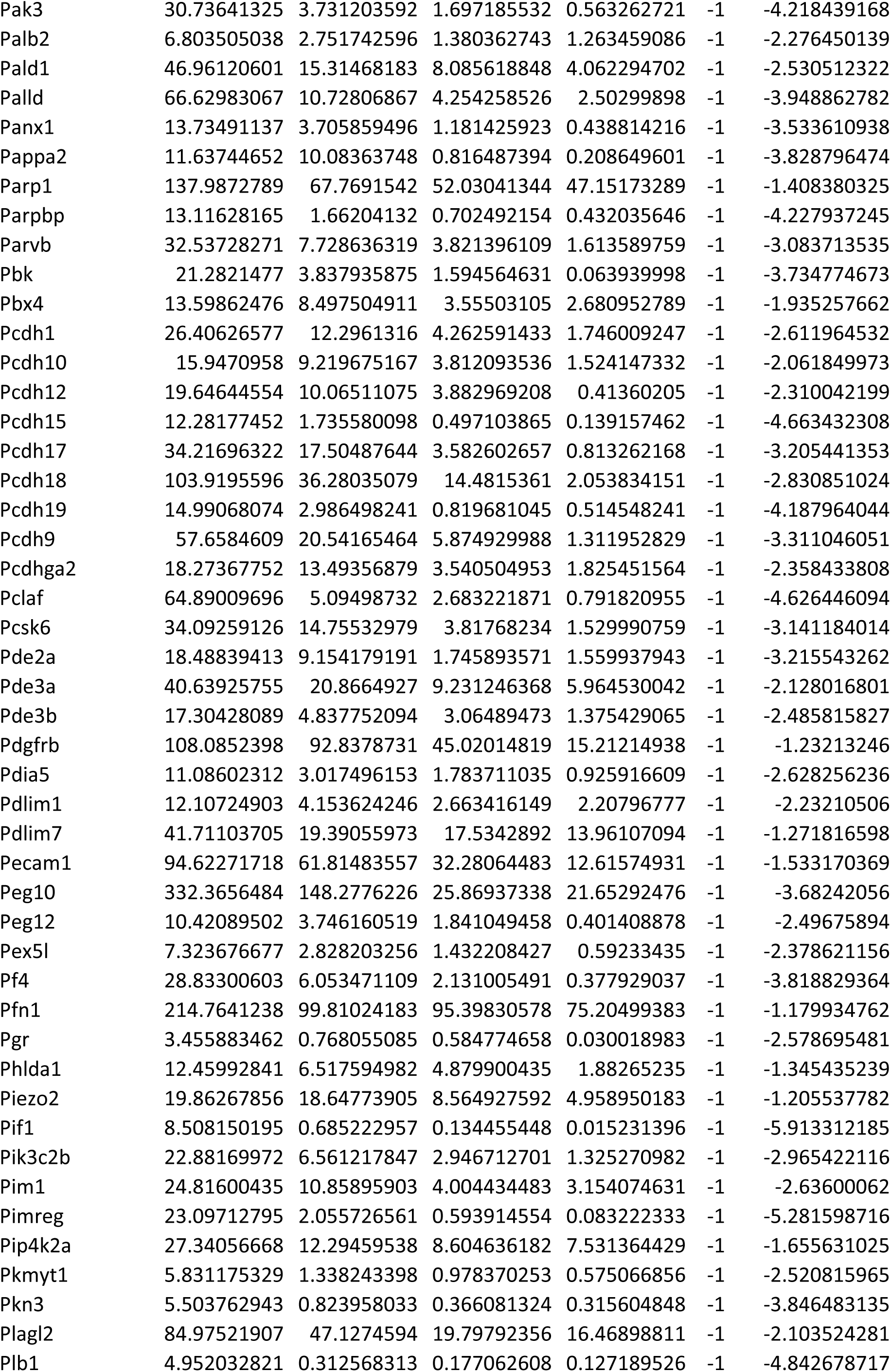
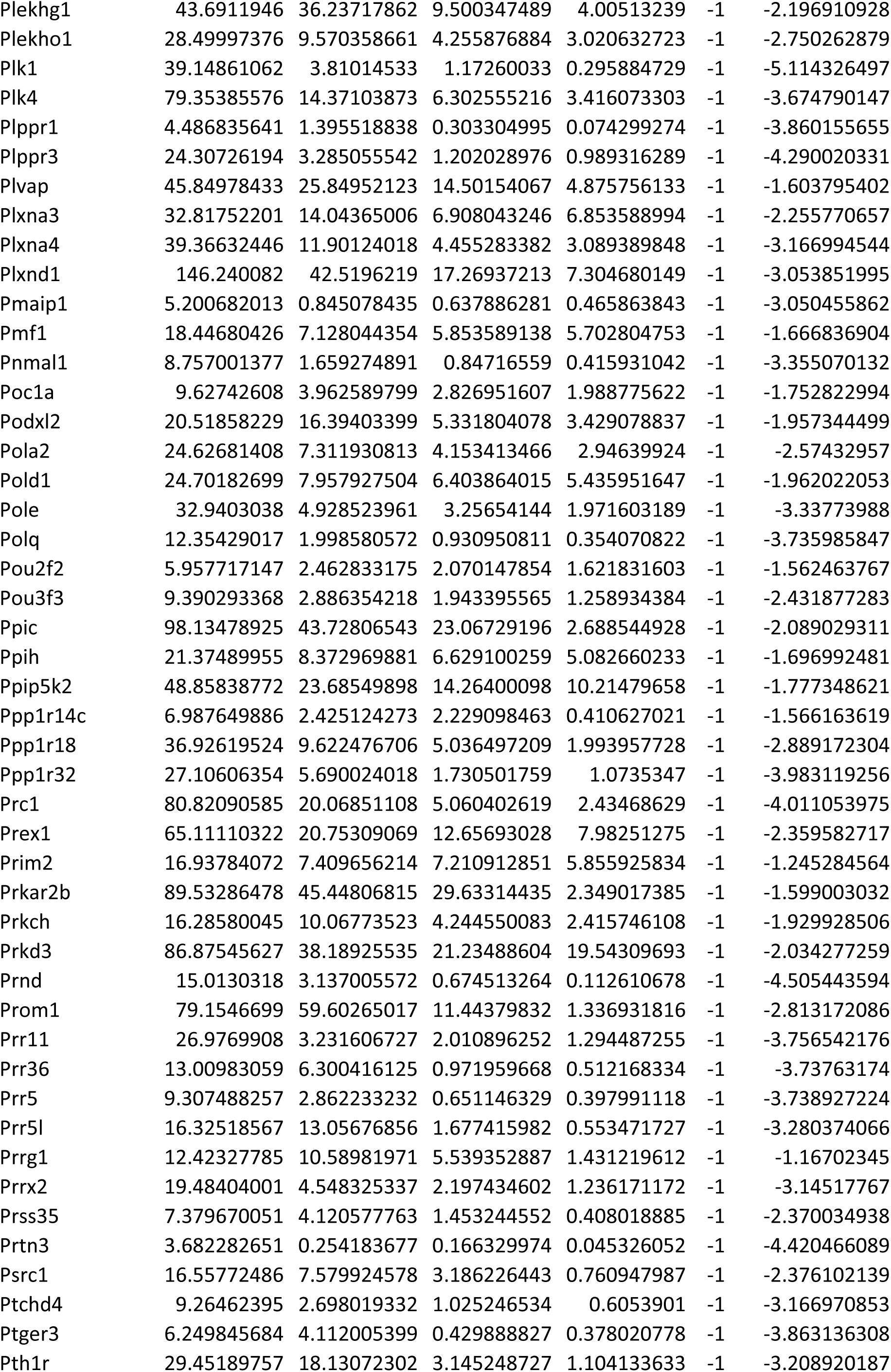
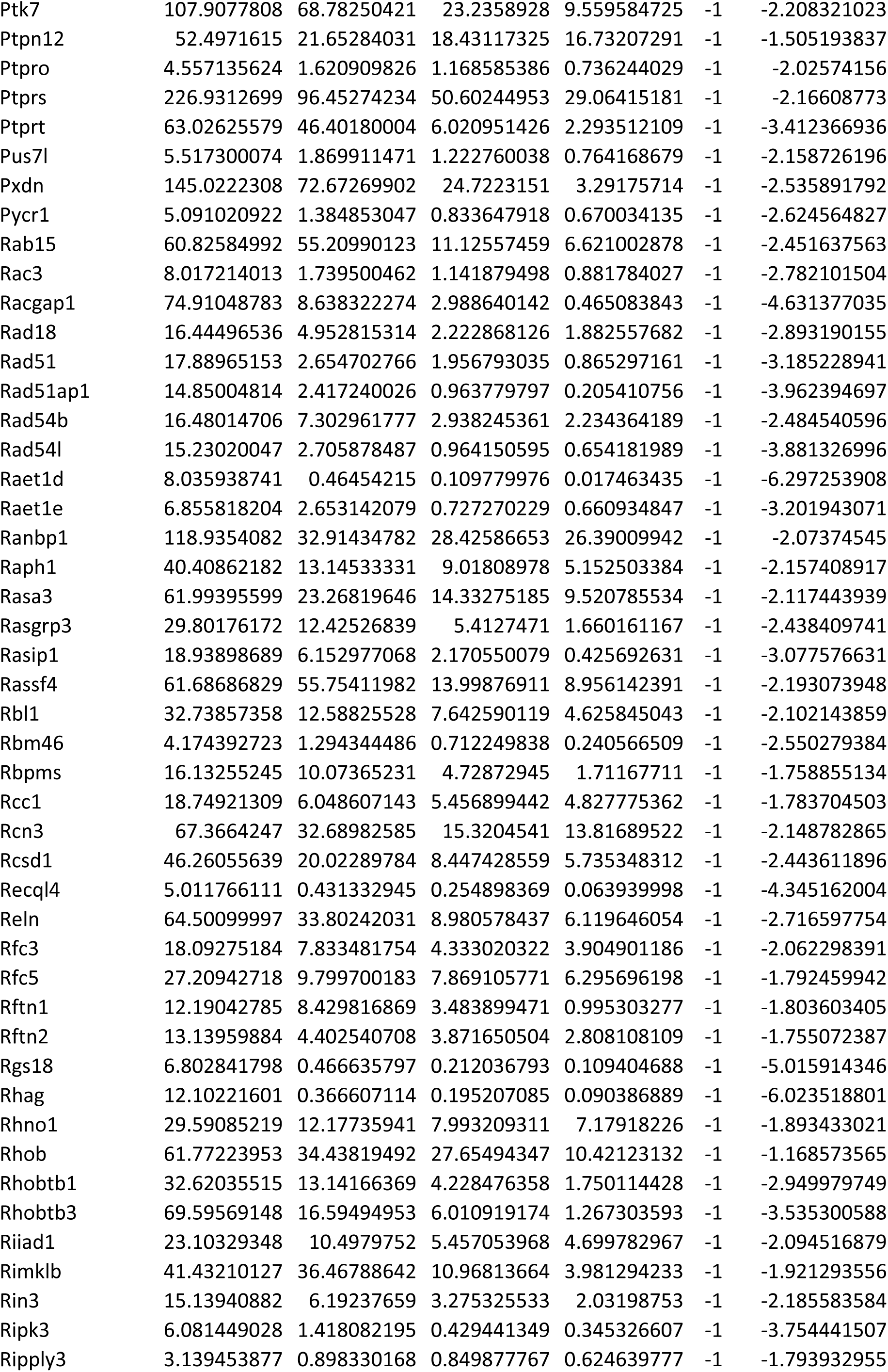
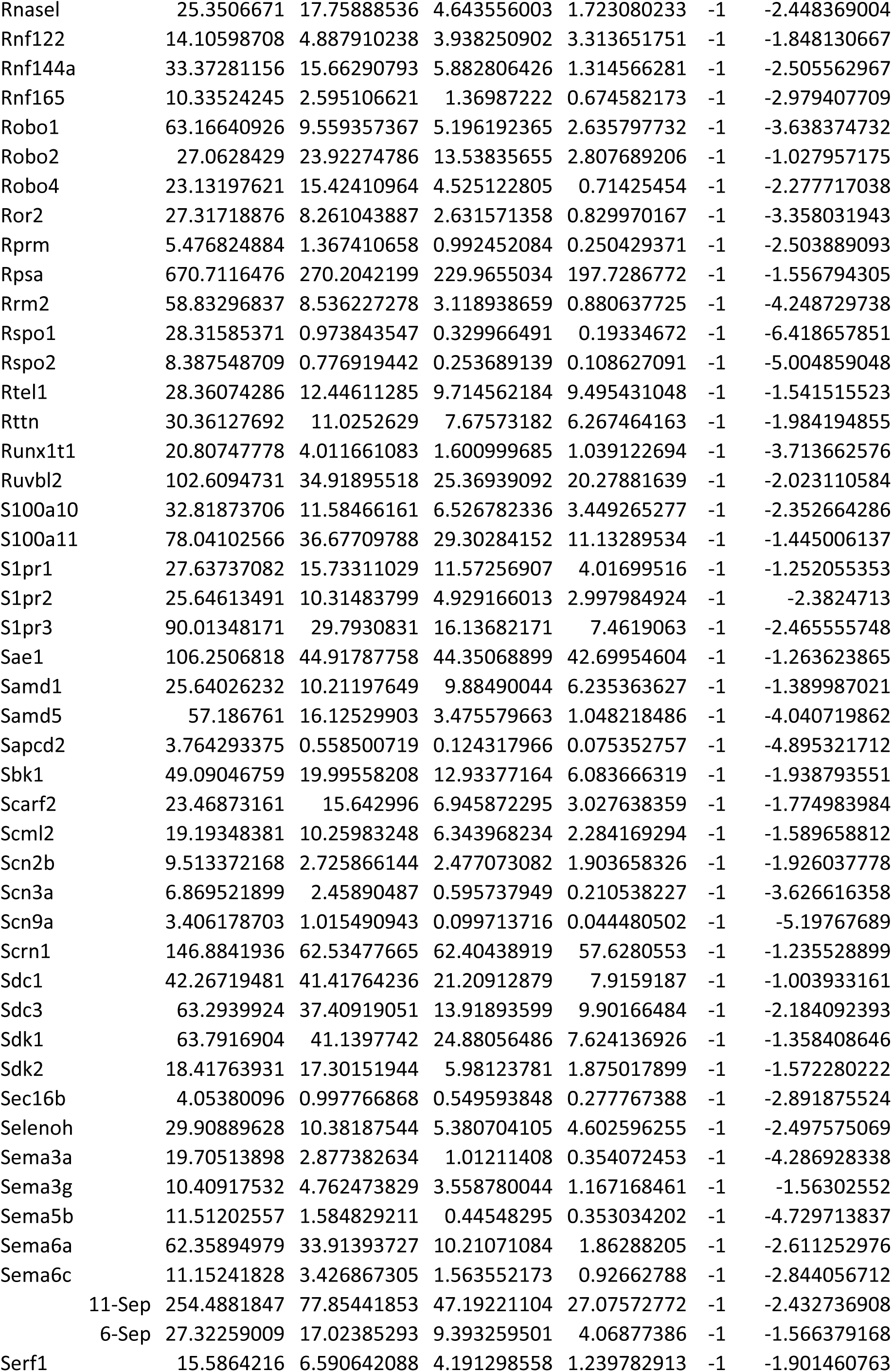
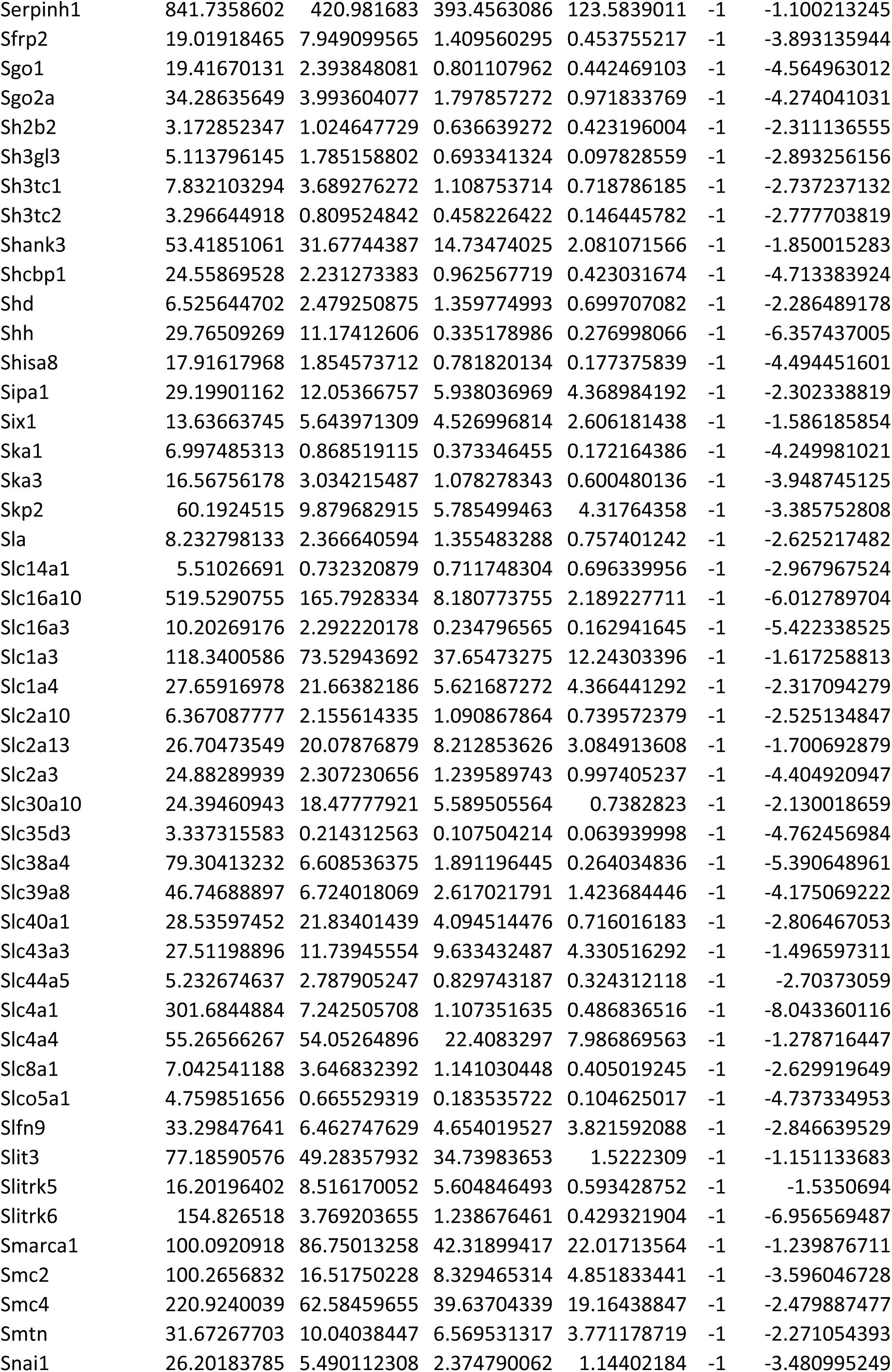
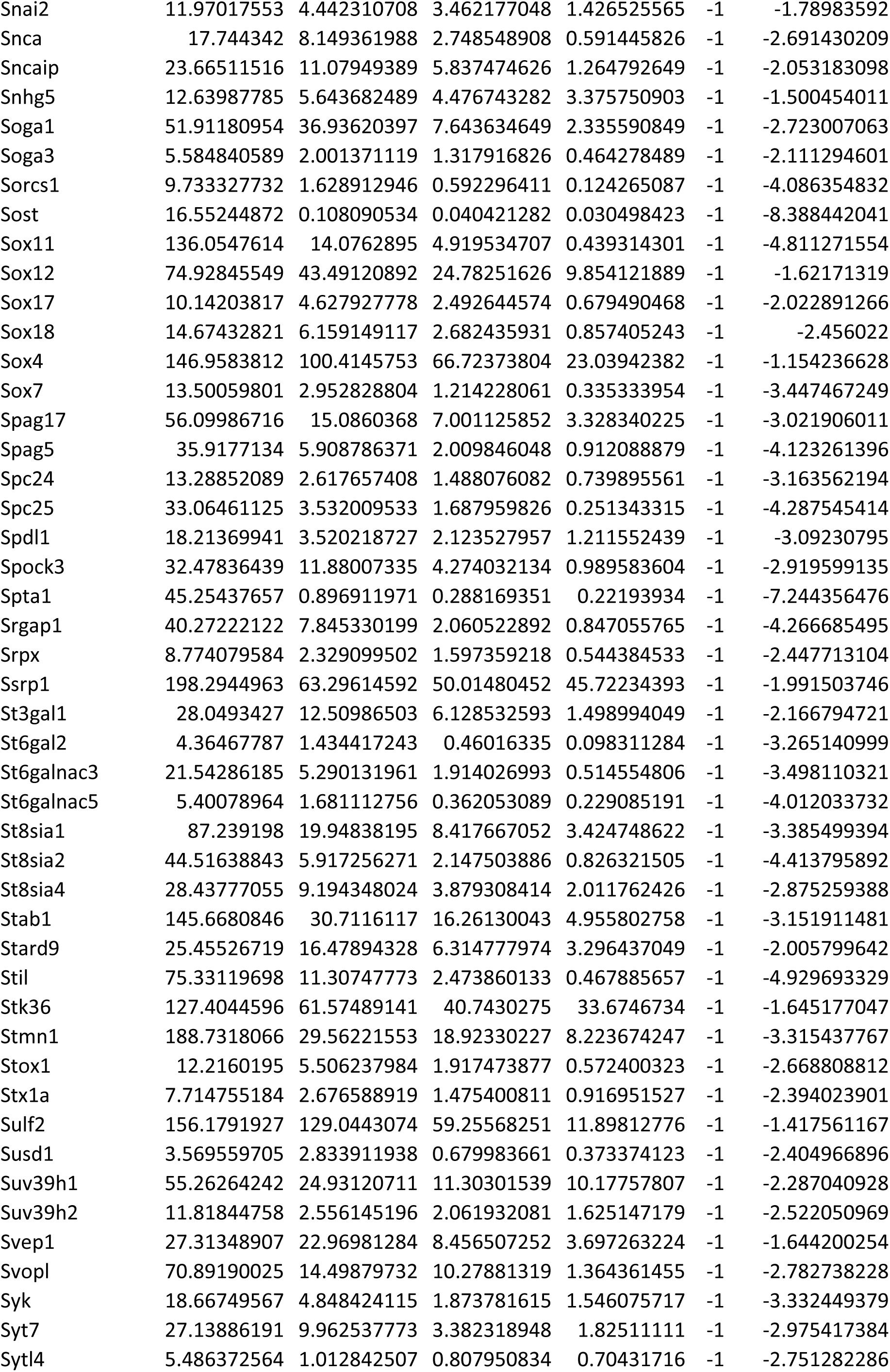
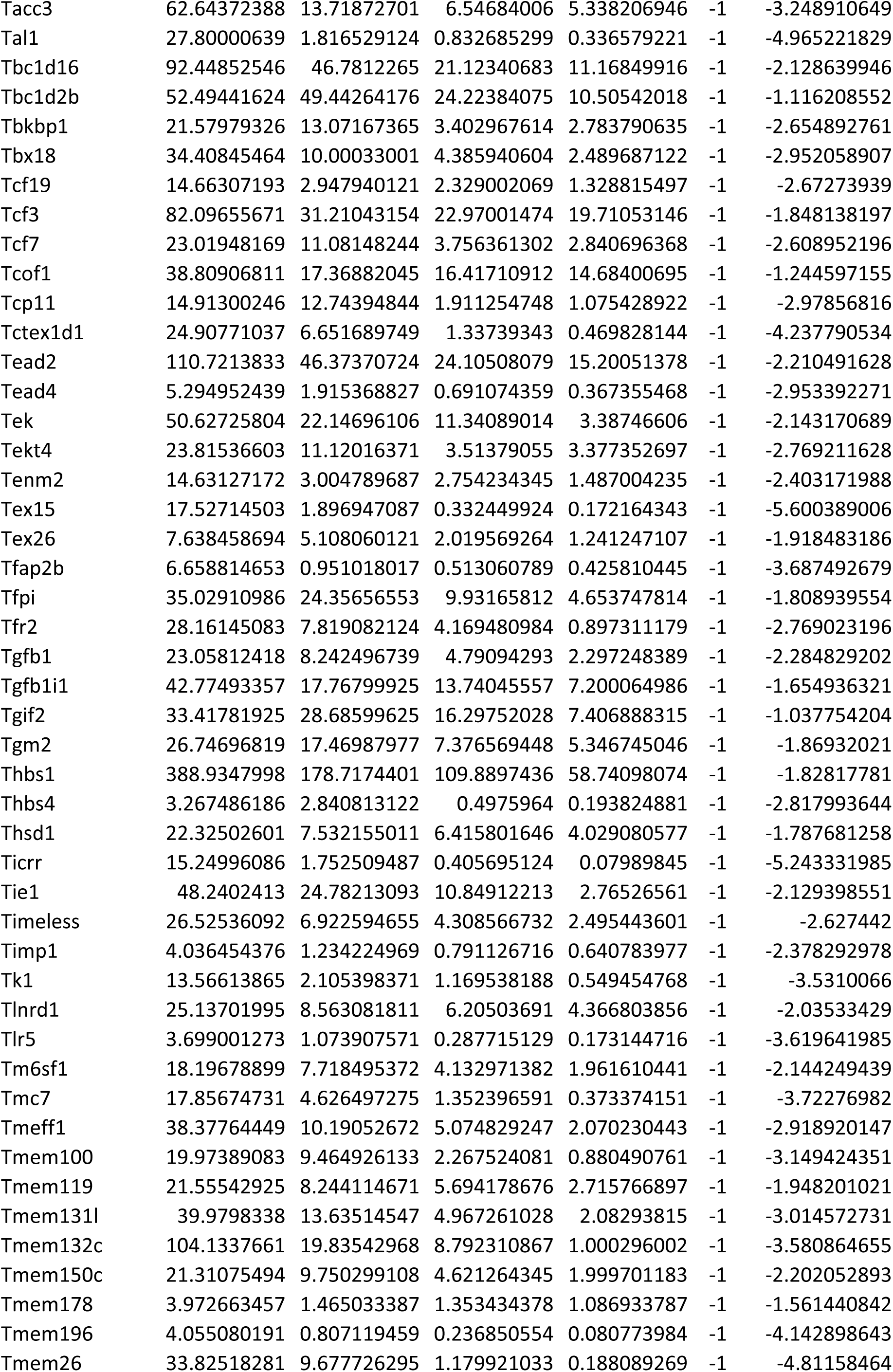
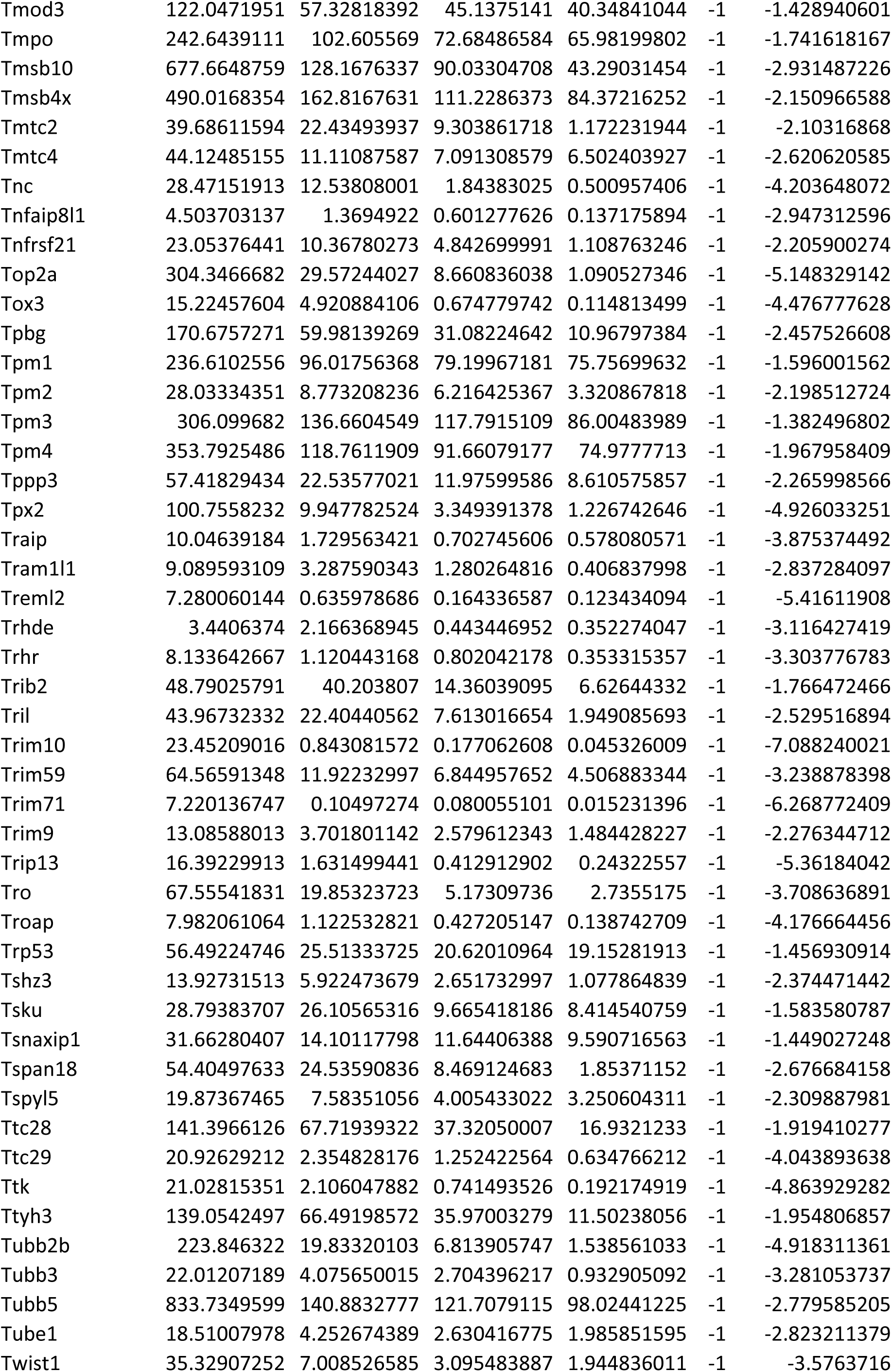
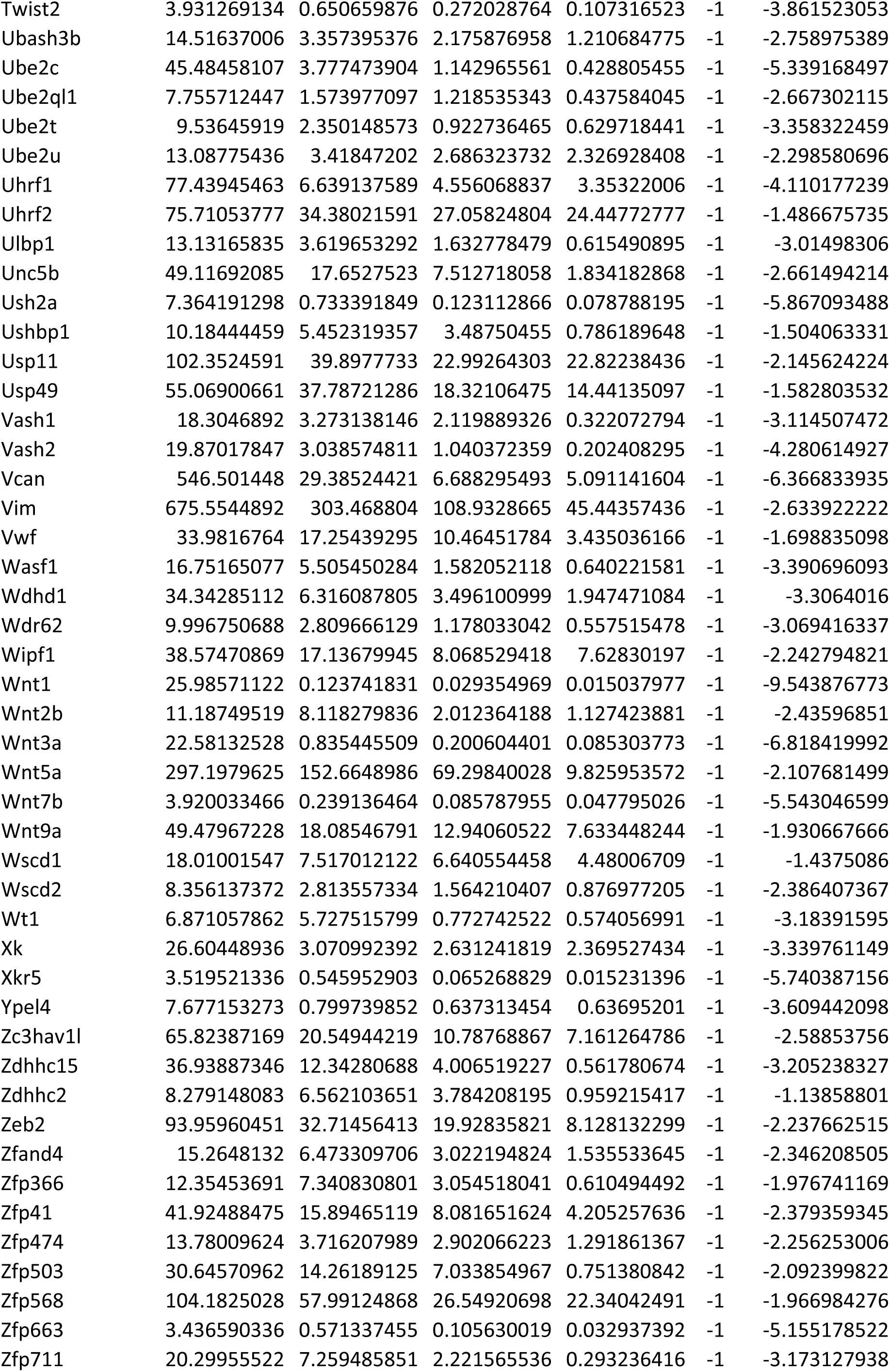
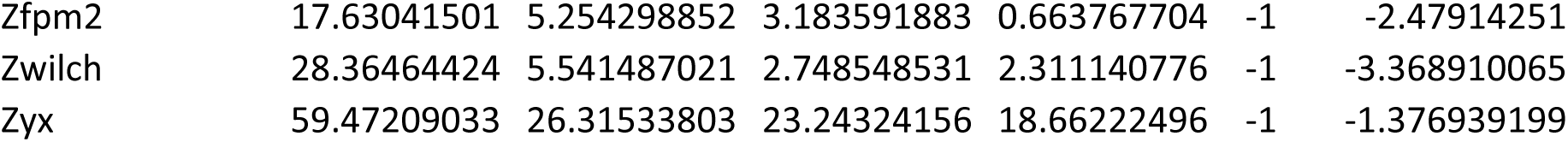
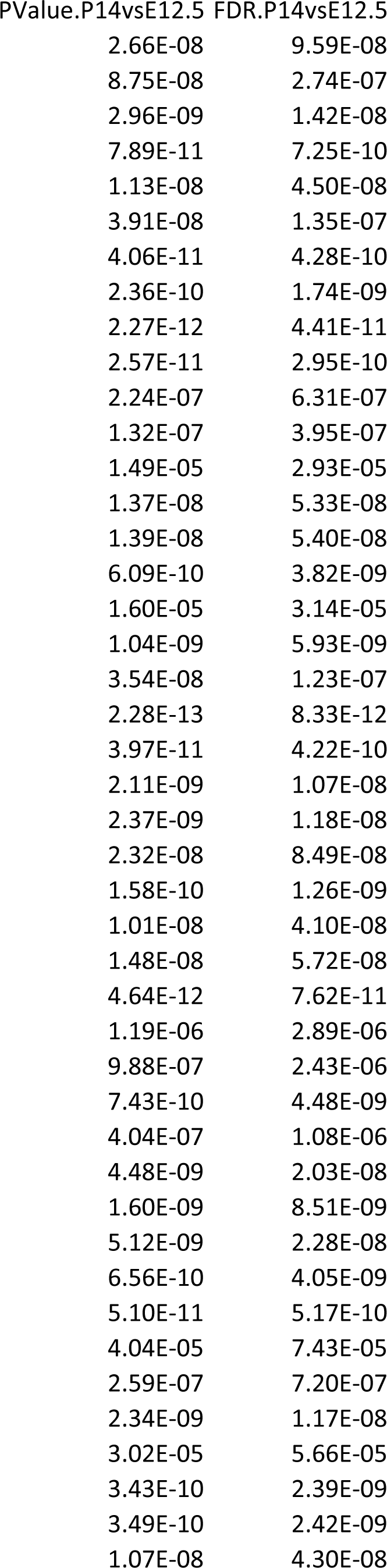
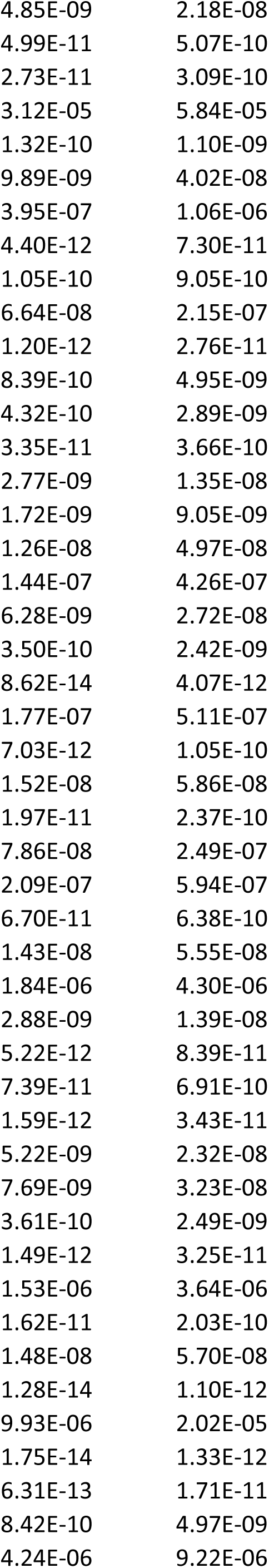
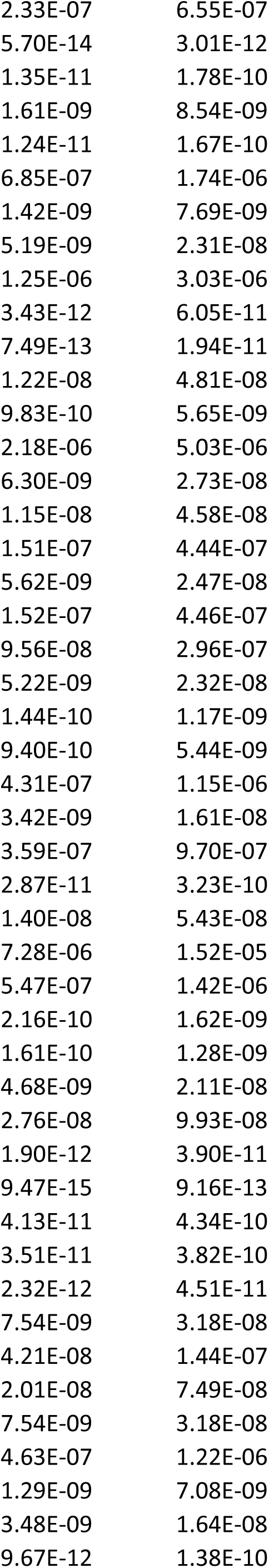
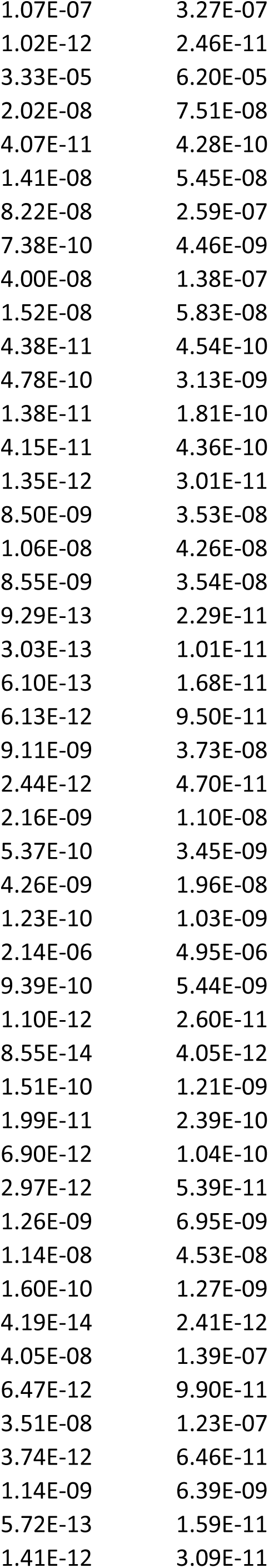
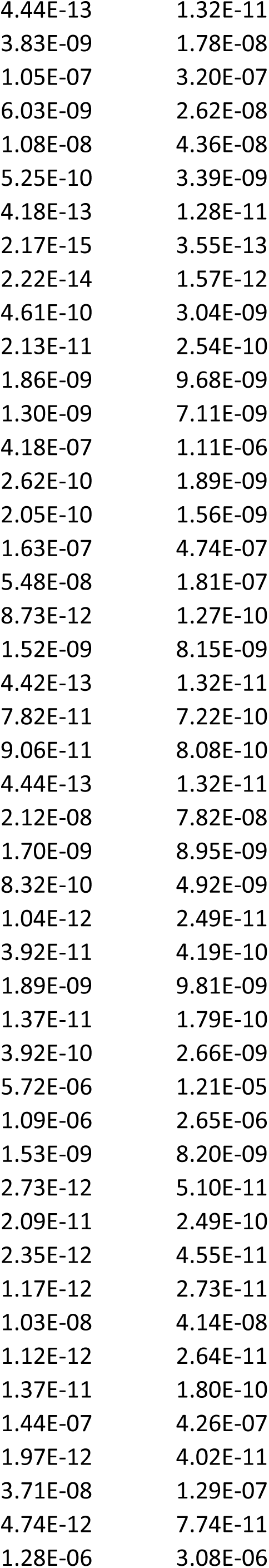
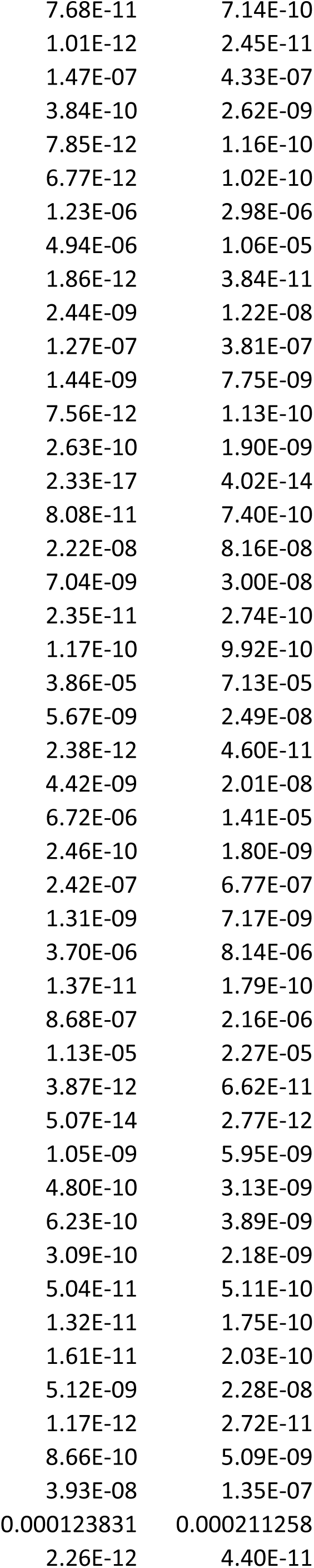
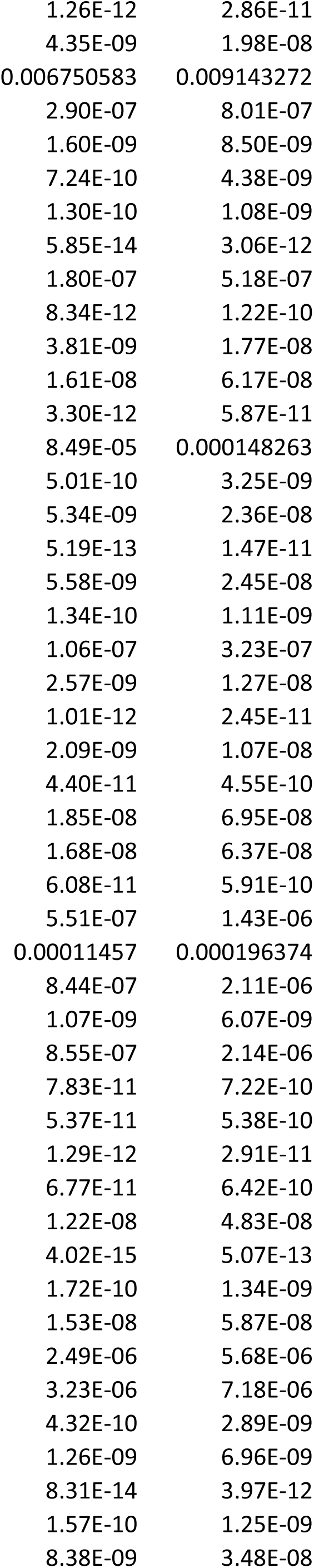
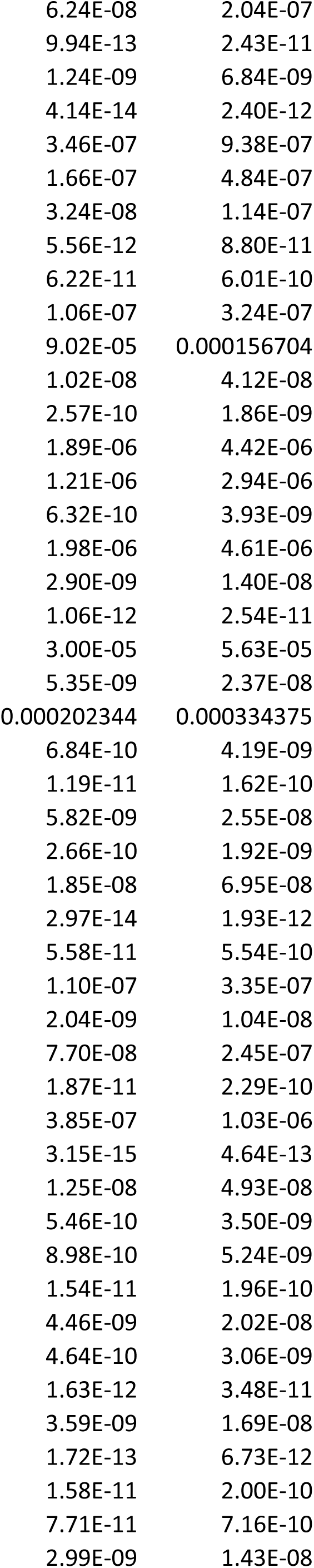
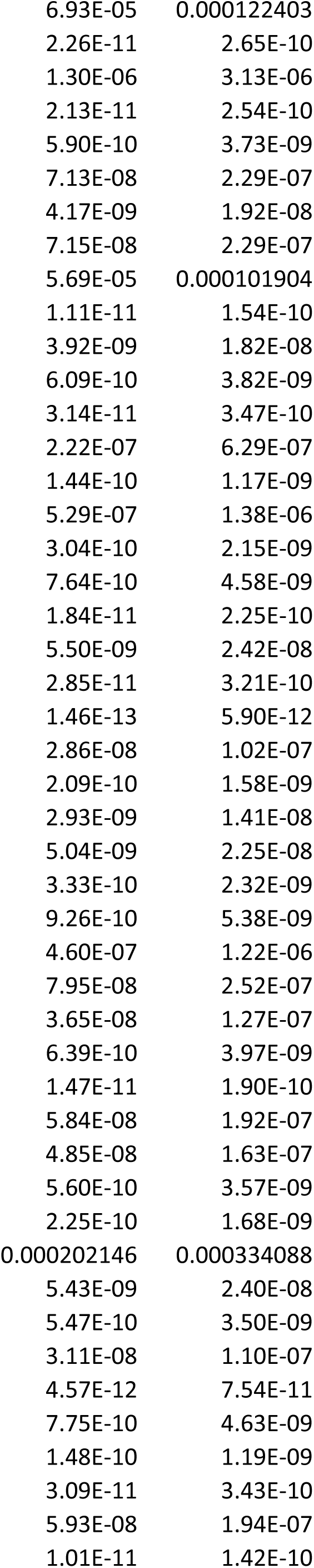
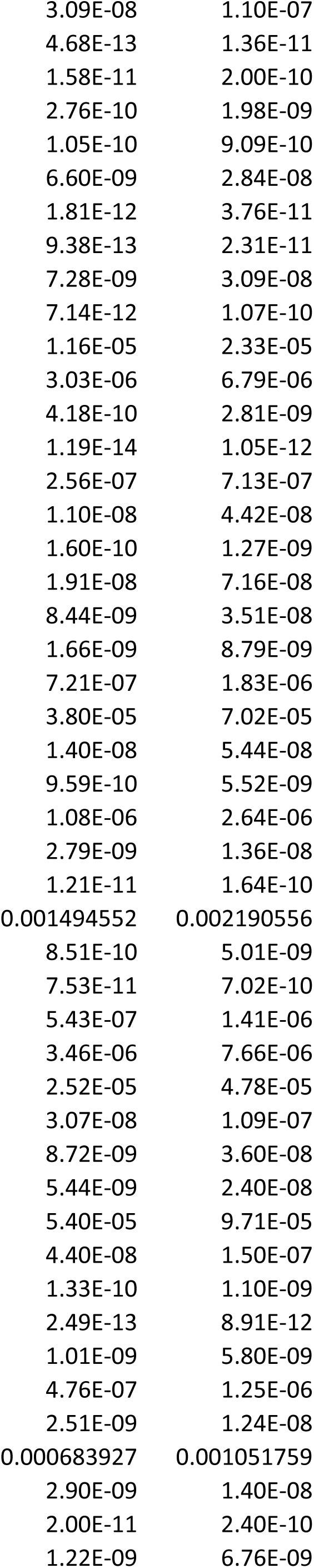
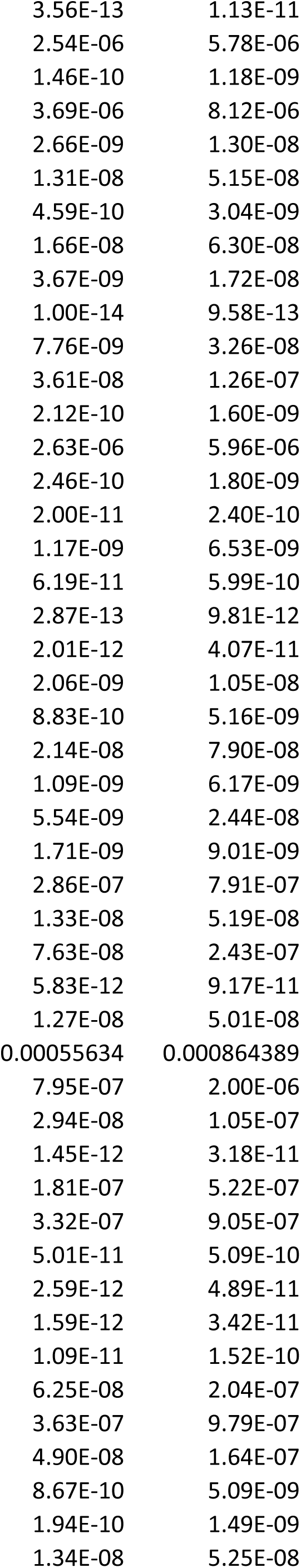
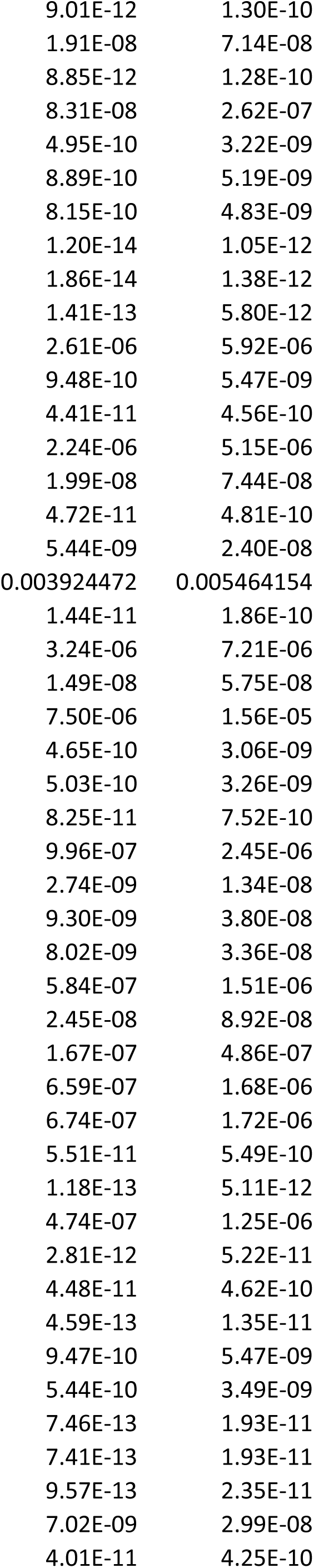
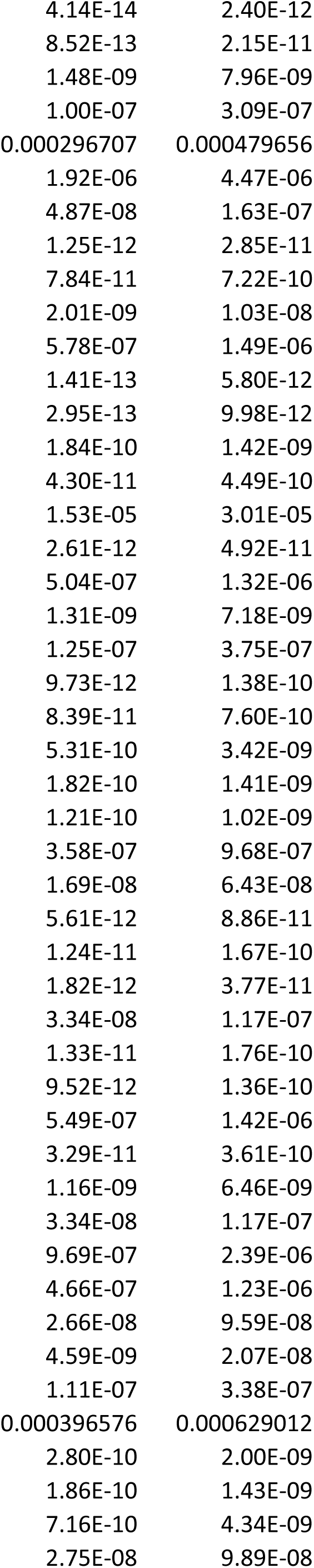
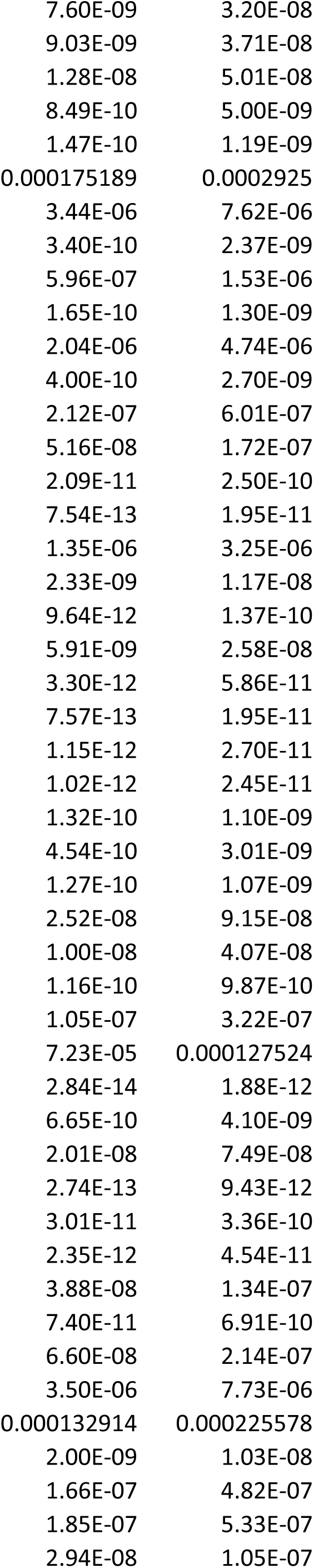
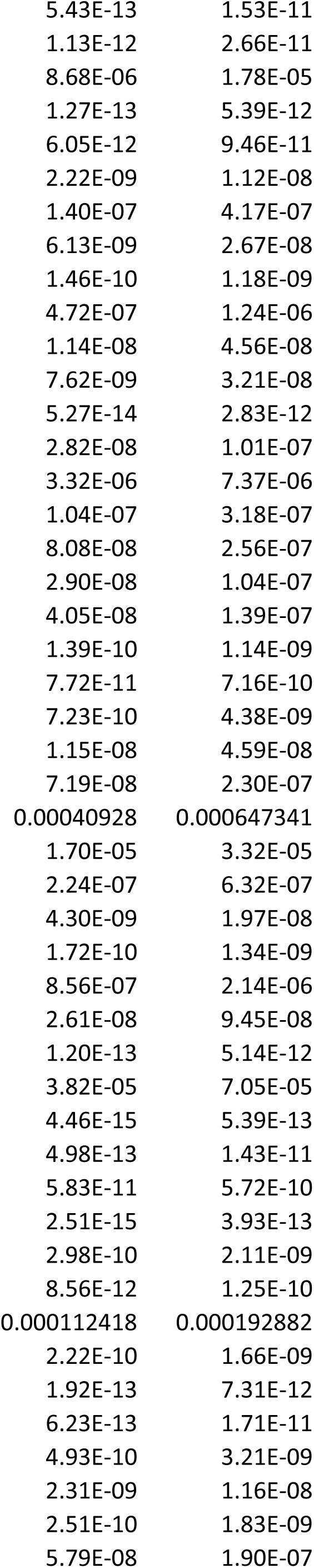
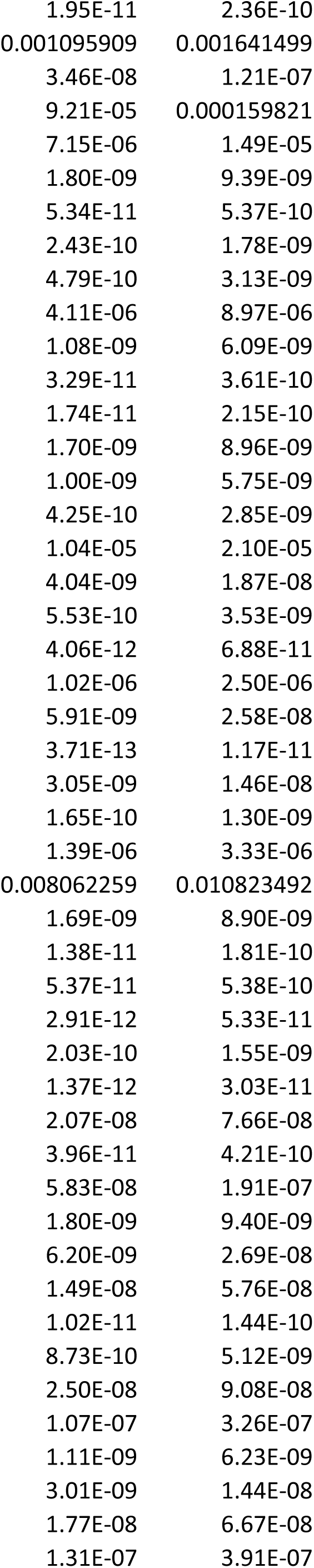
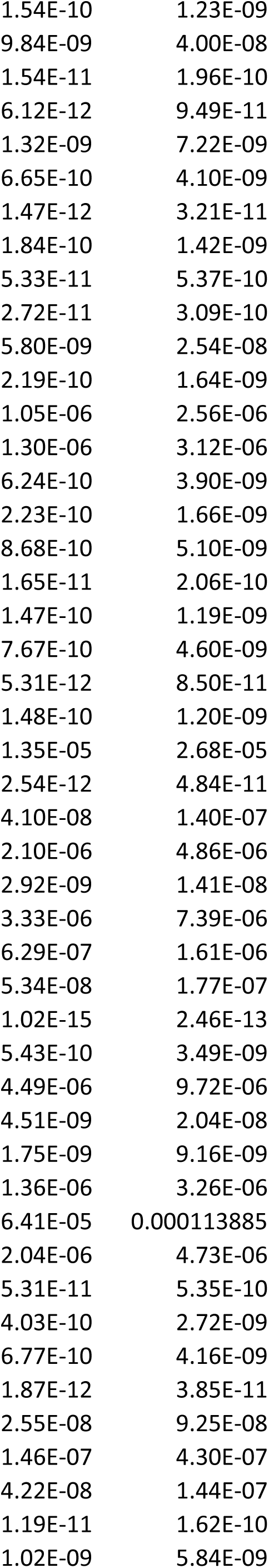
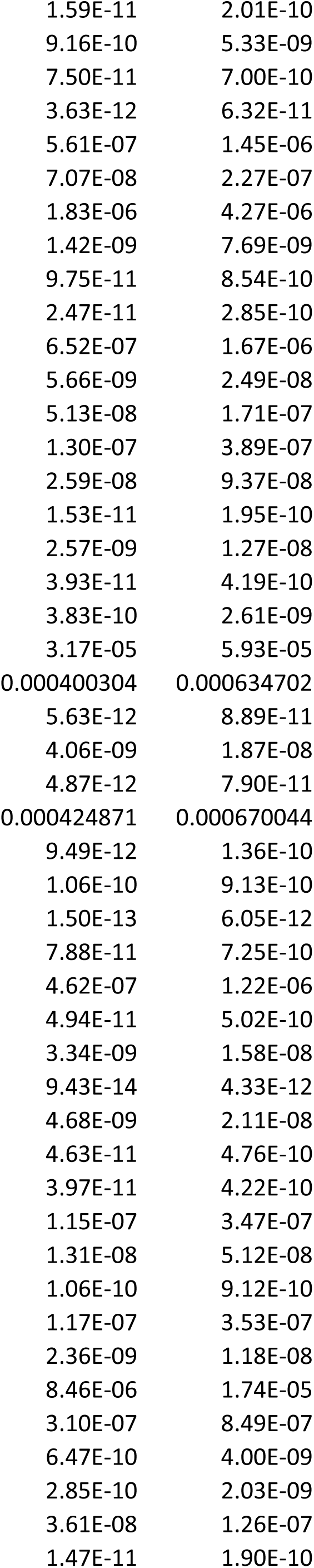
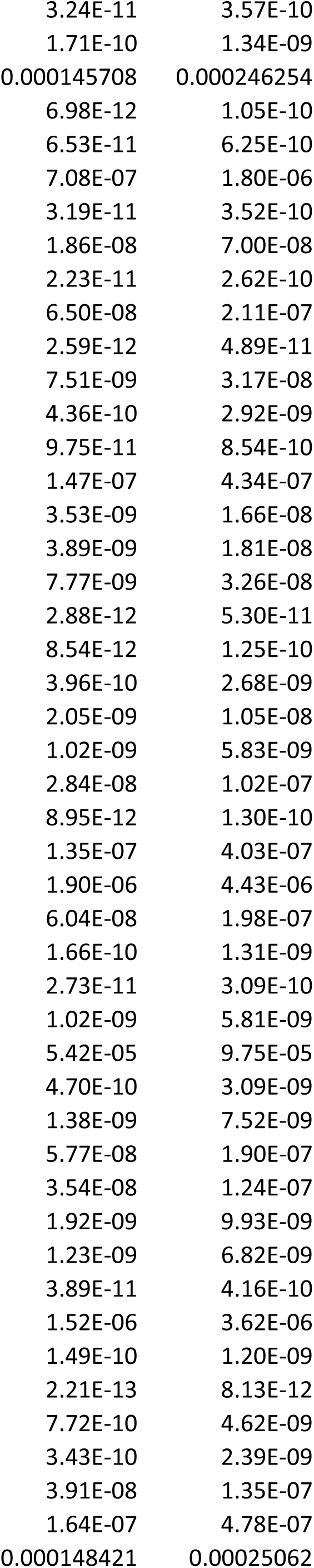
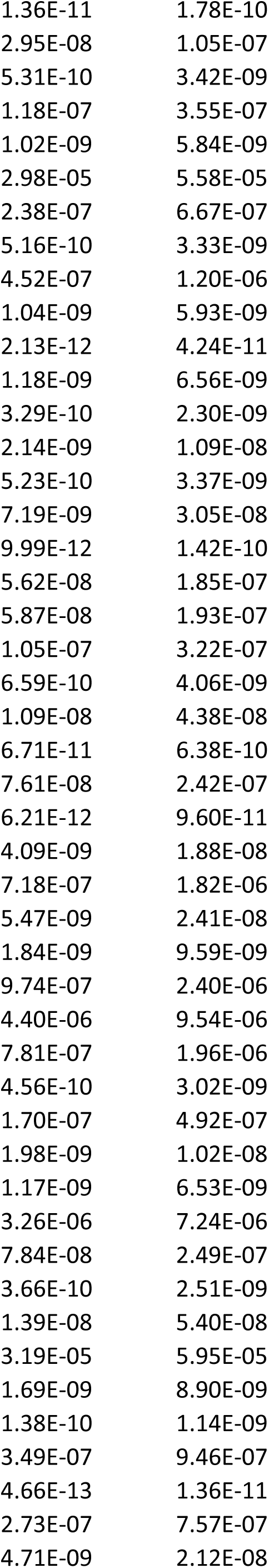
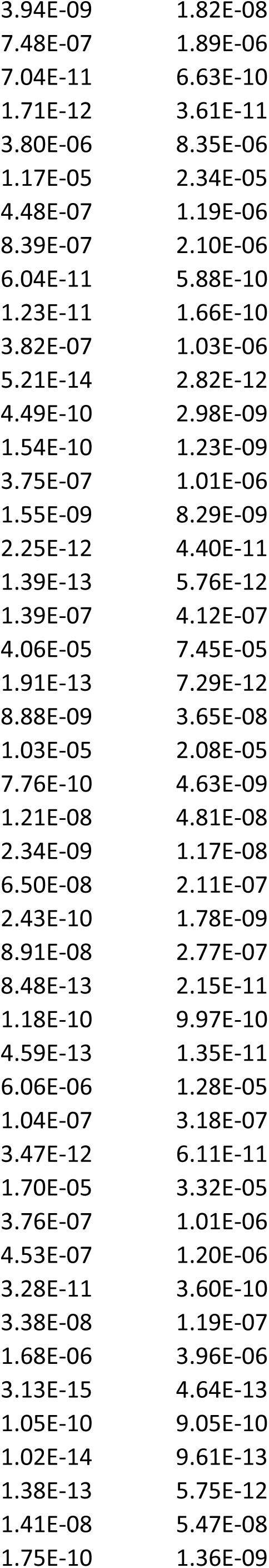
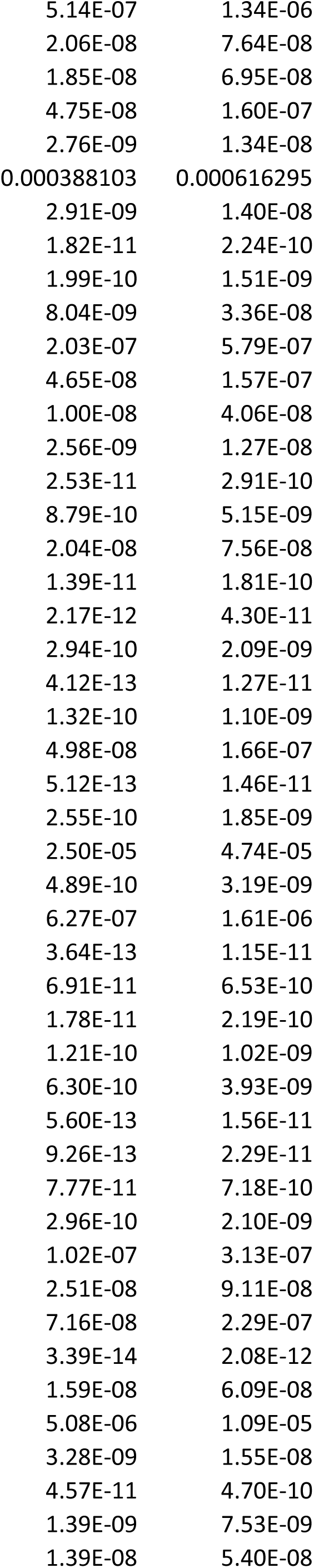
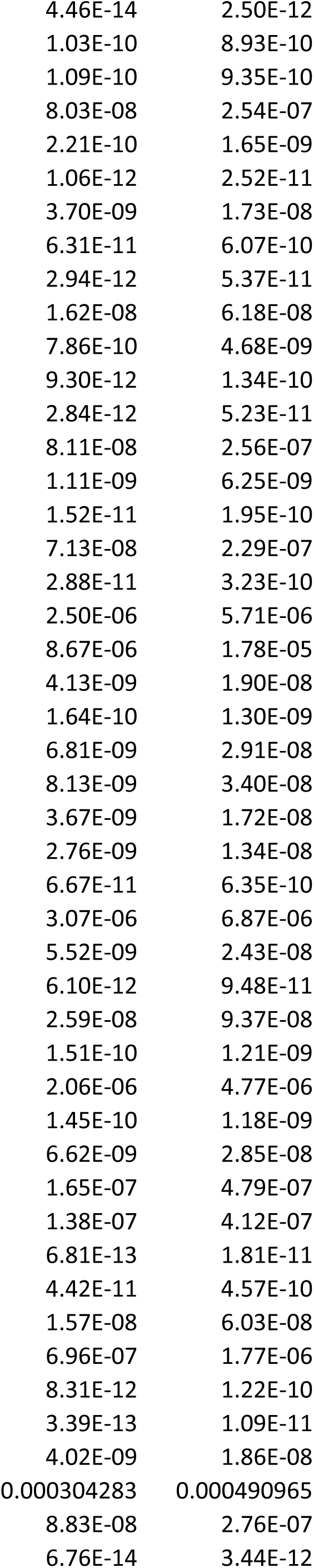
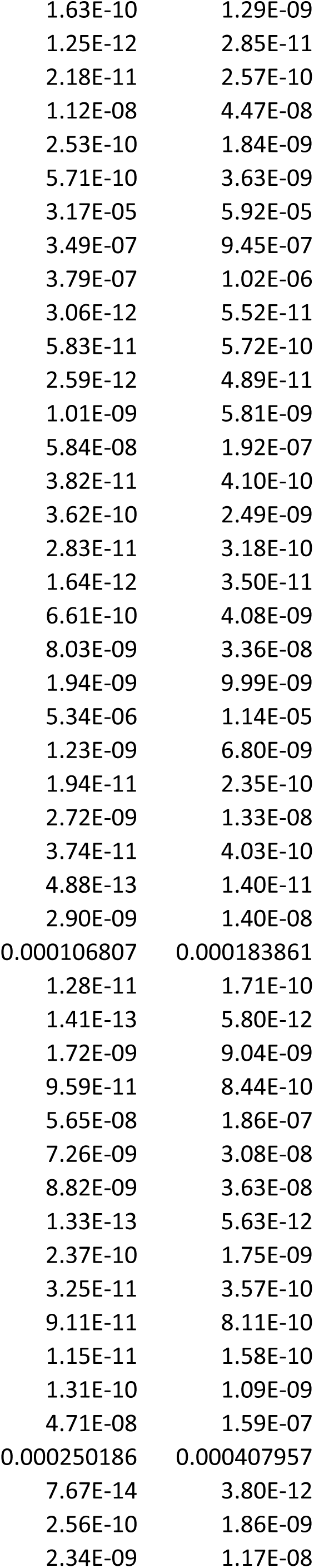
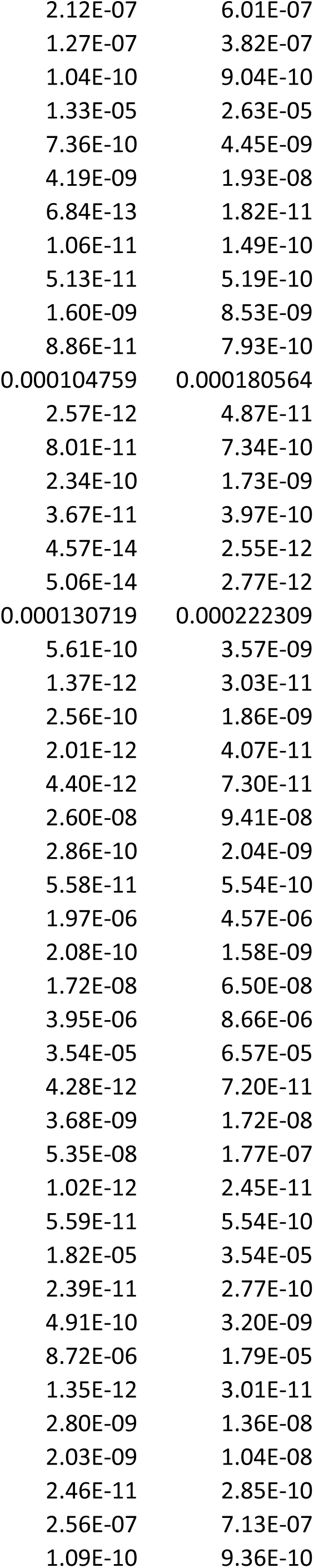
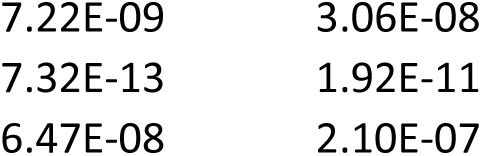
Gene expression during choroid plexus development. Table S2.1 Developmentally downregulated genes during choroid plexus development (E12.5, P0, P14, 6 months) PValue.P14vsE12.5 FDR.P14vsE12.5

**Table S2.**
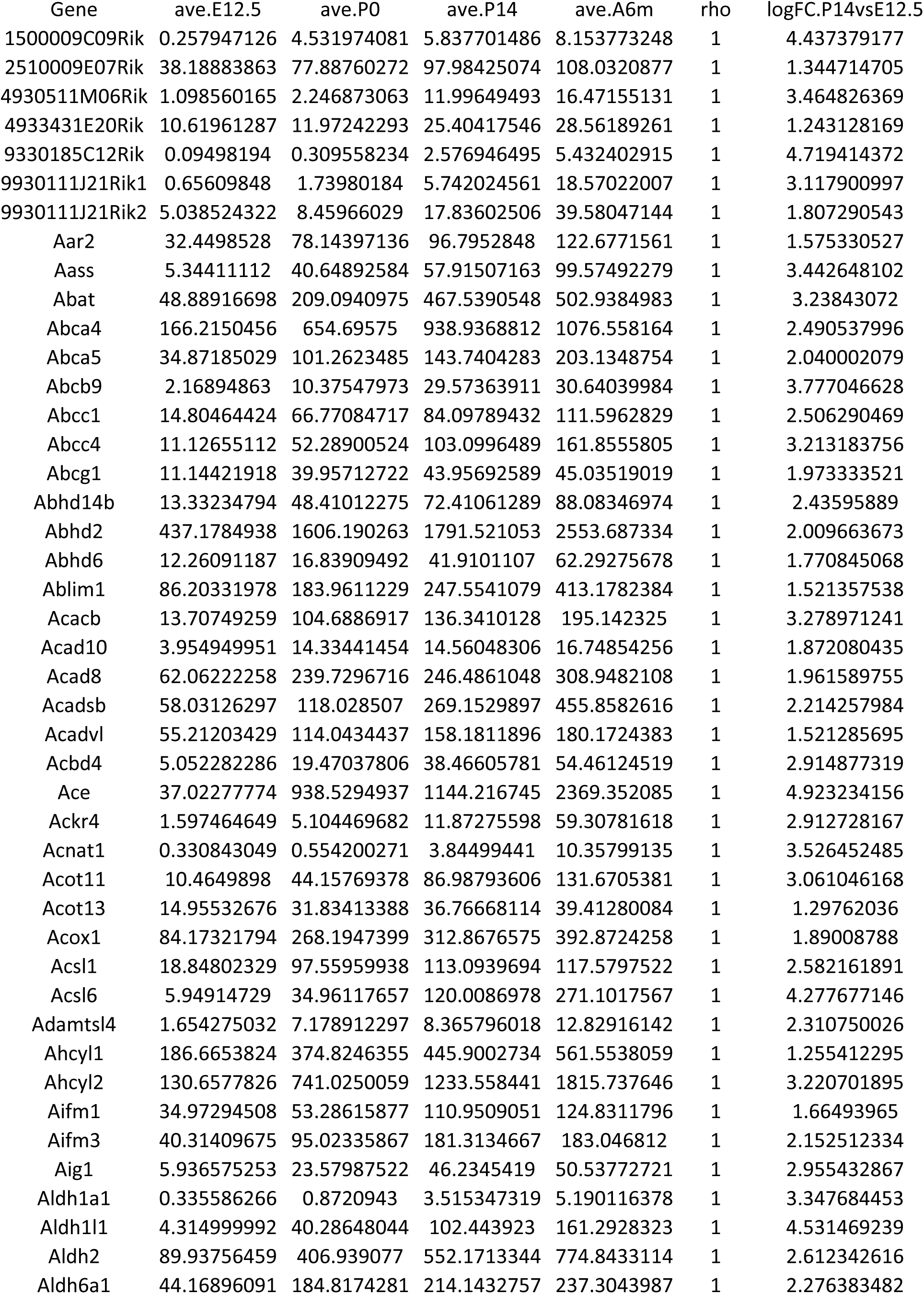
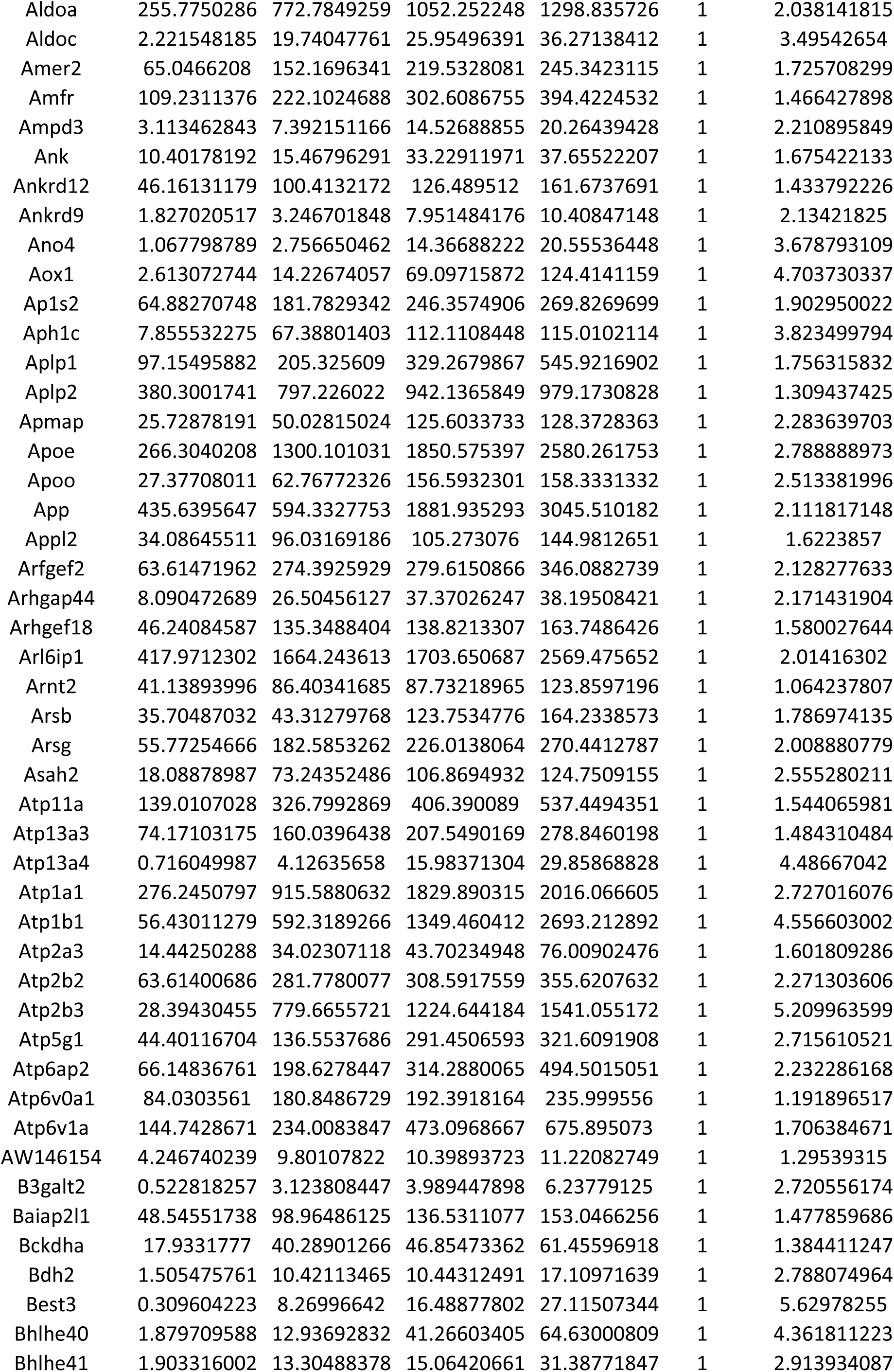
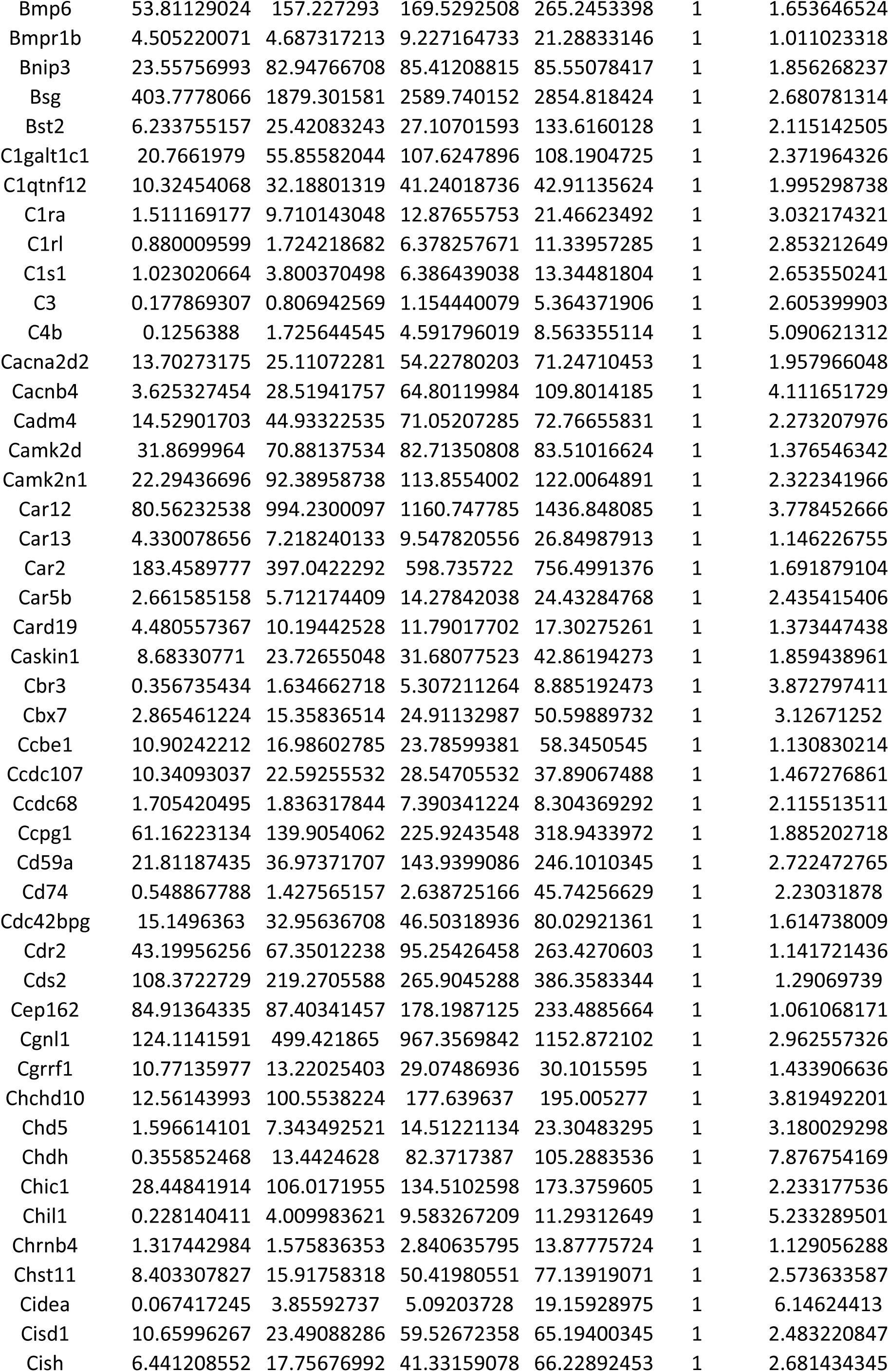
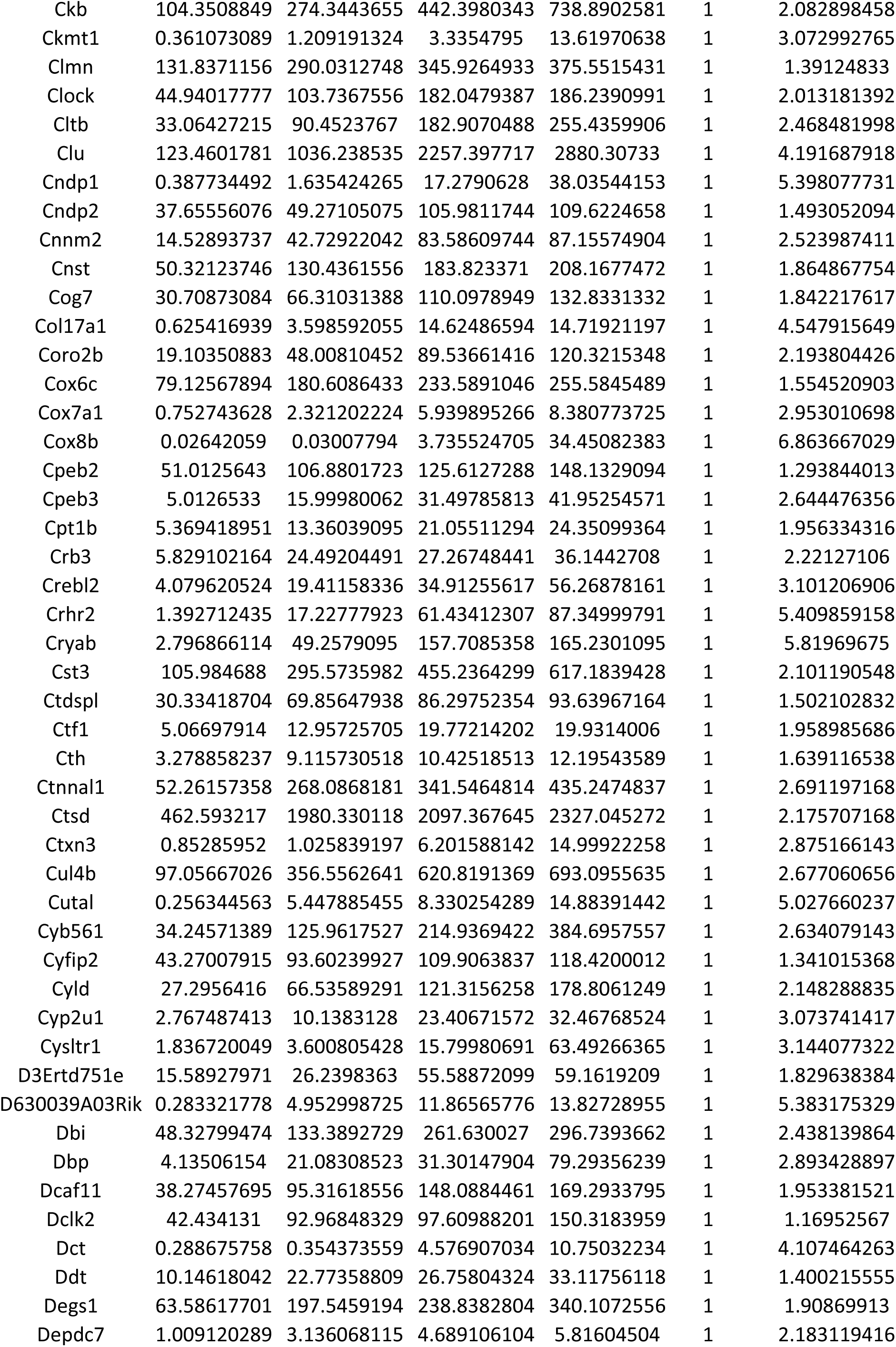
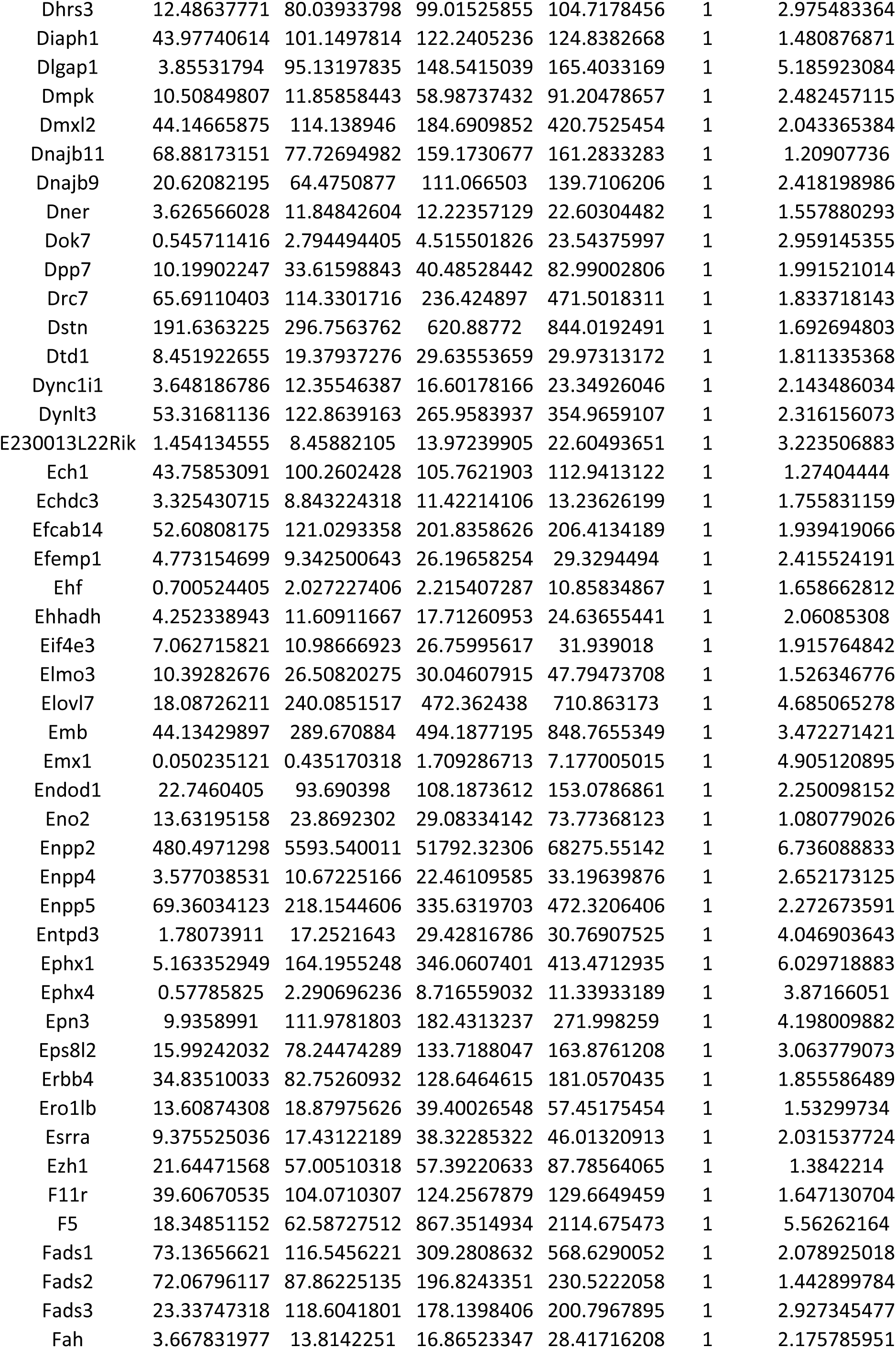
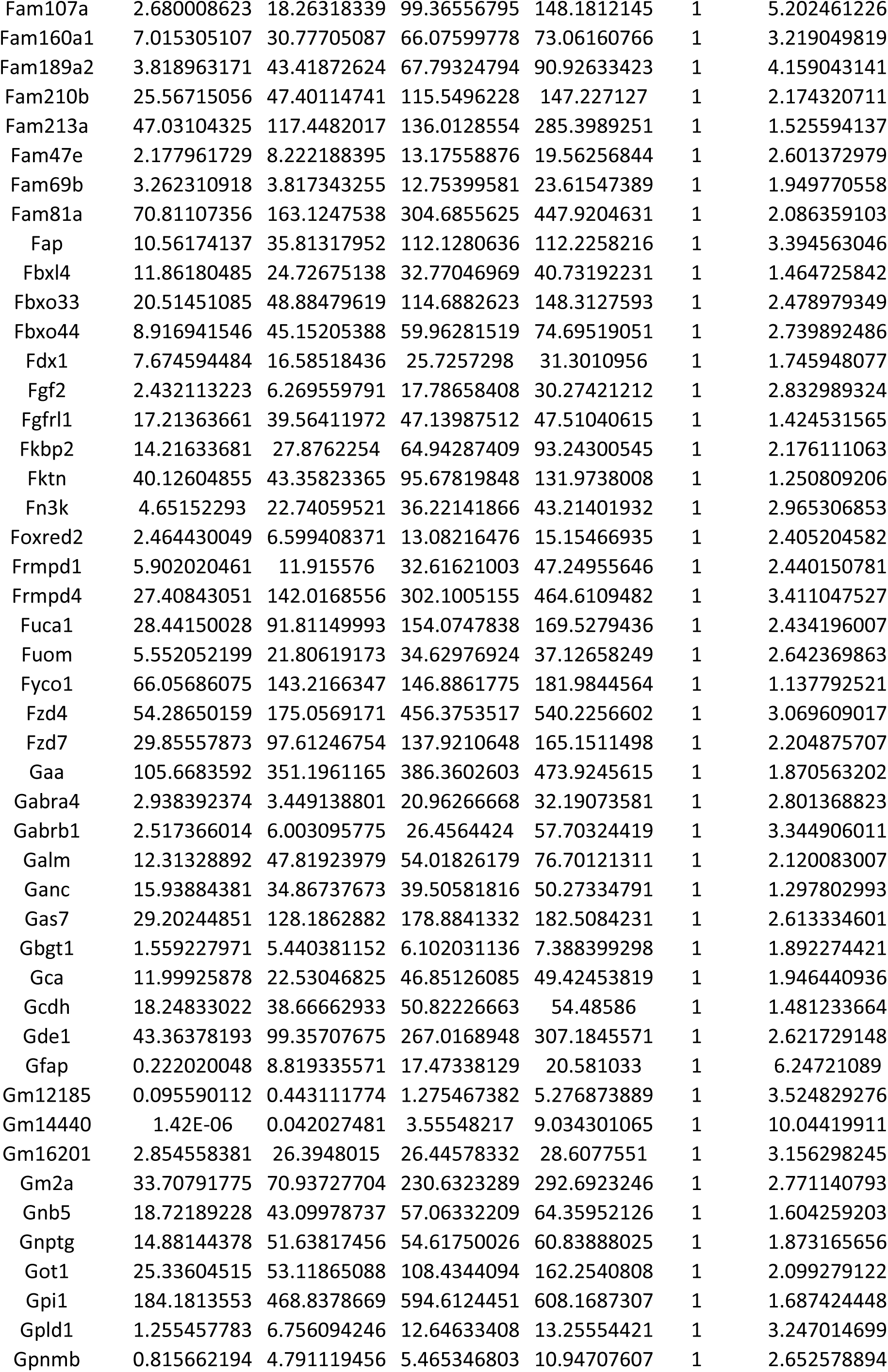
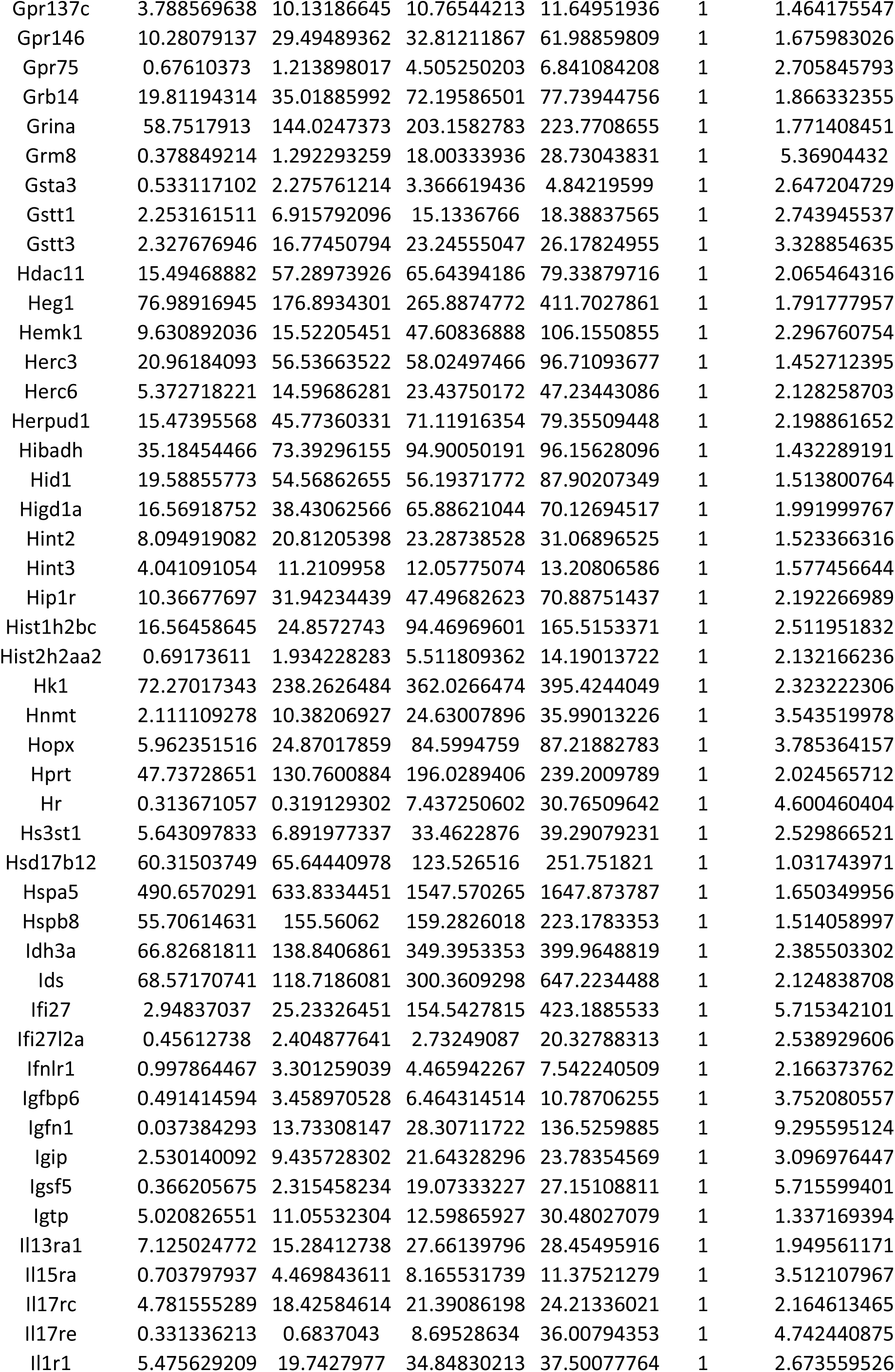
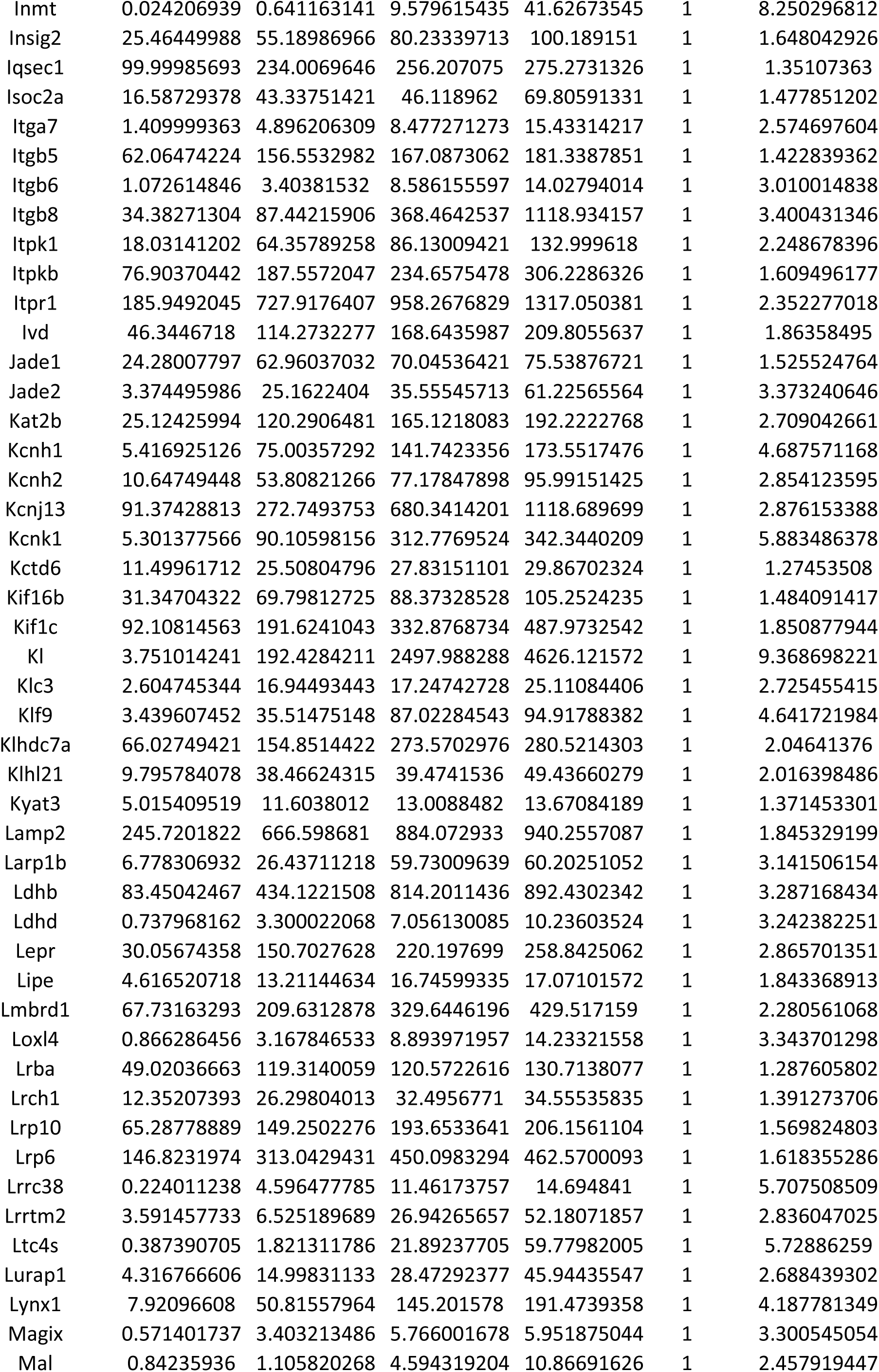
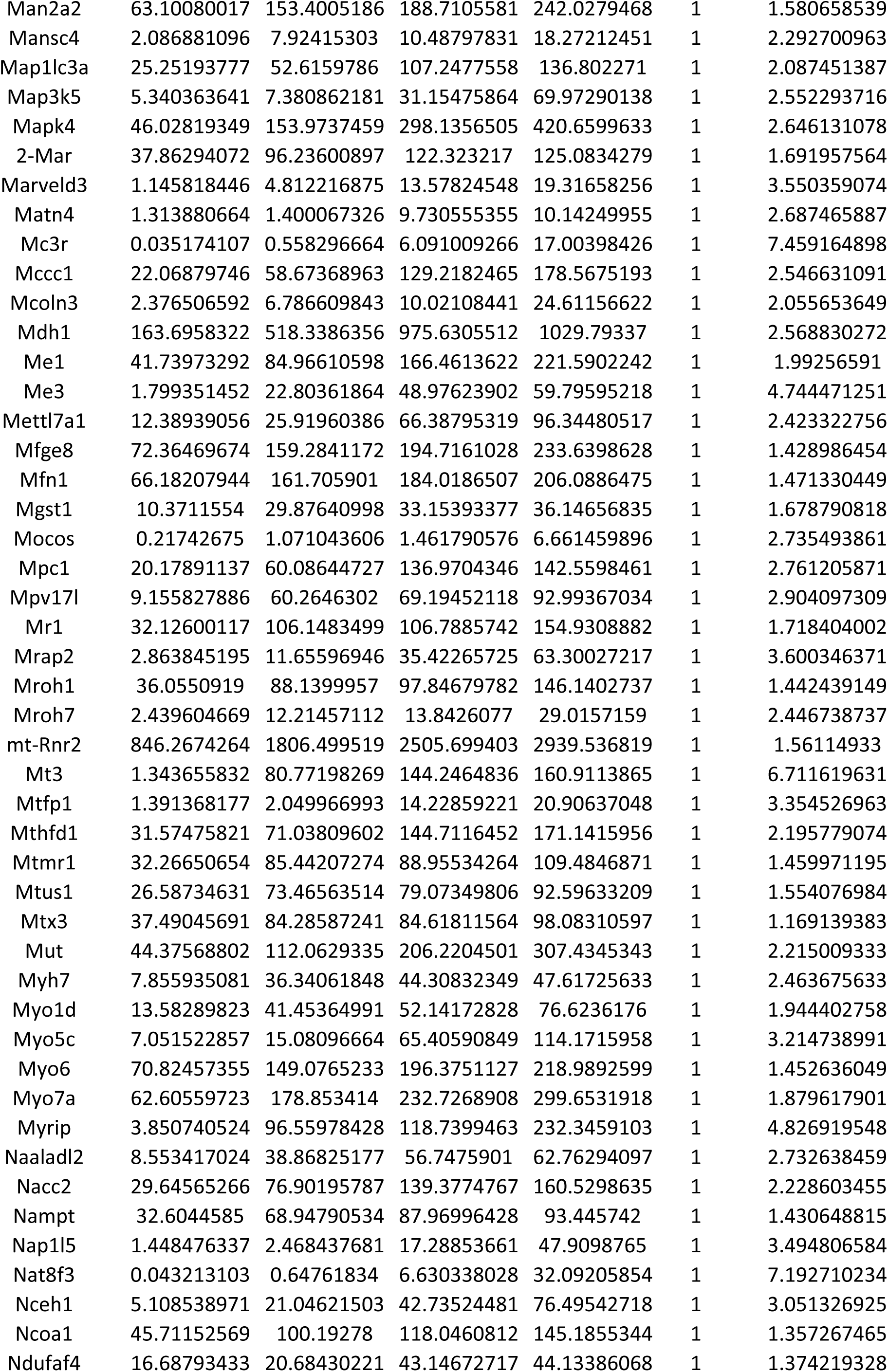
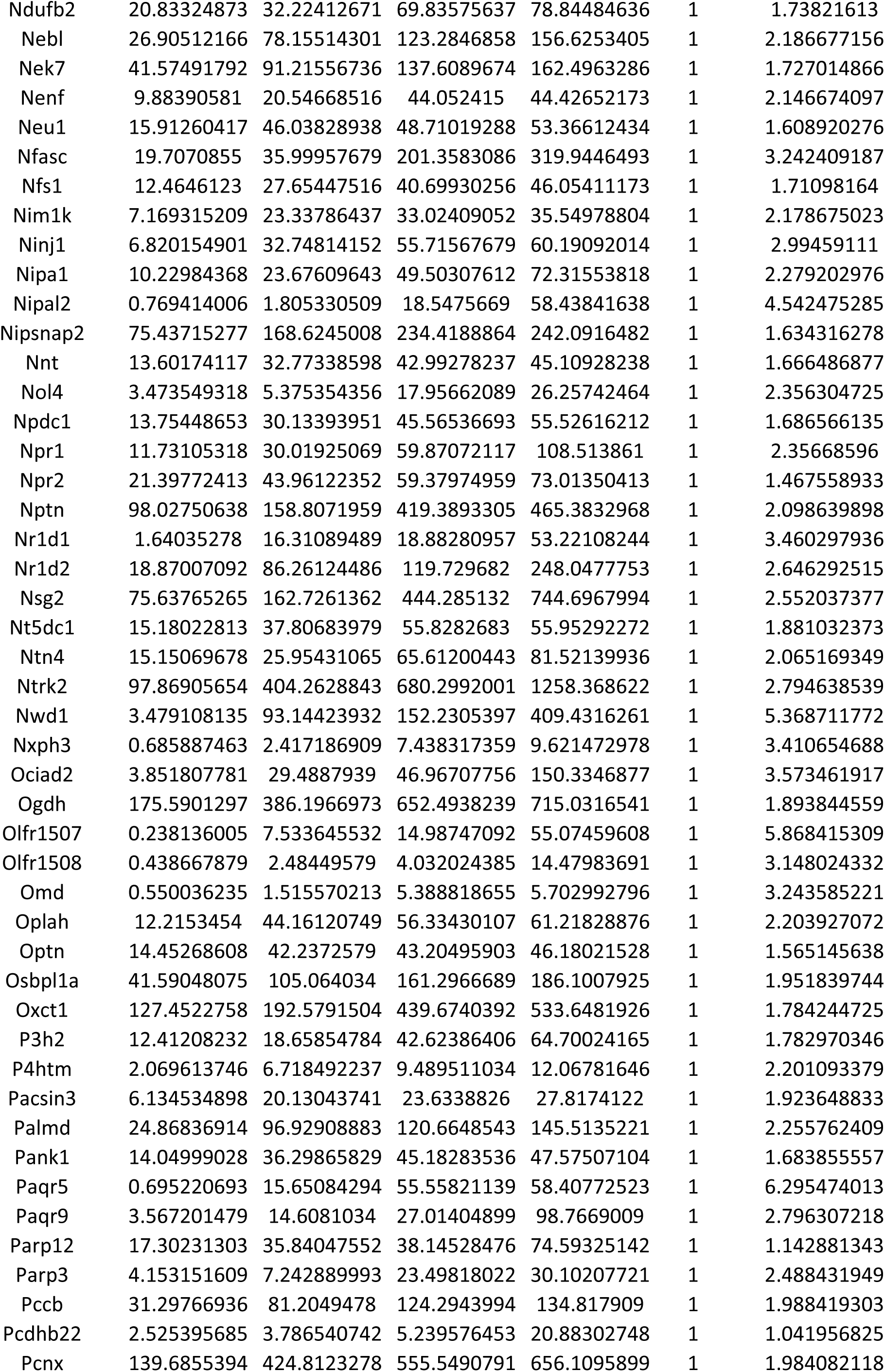
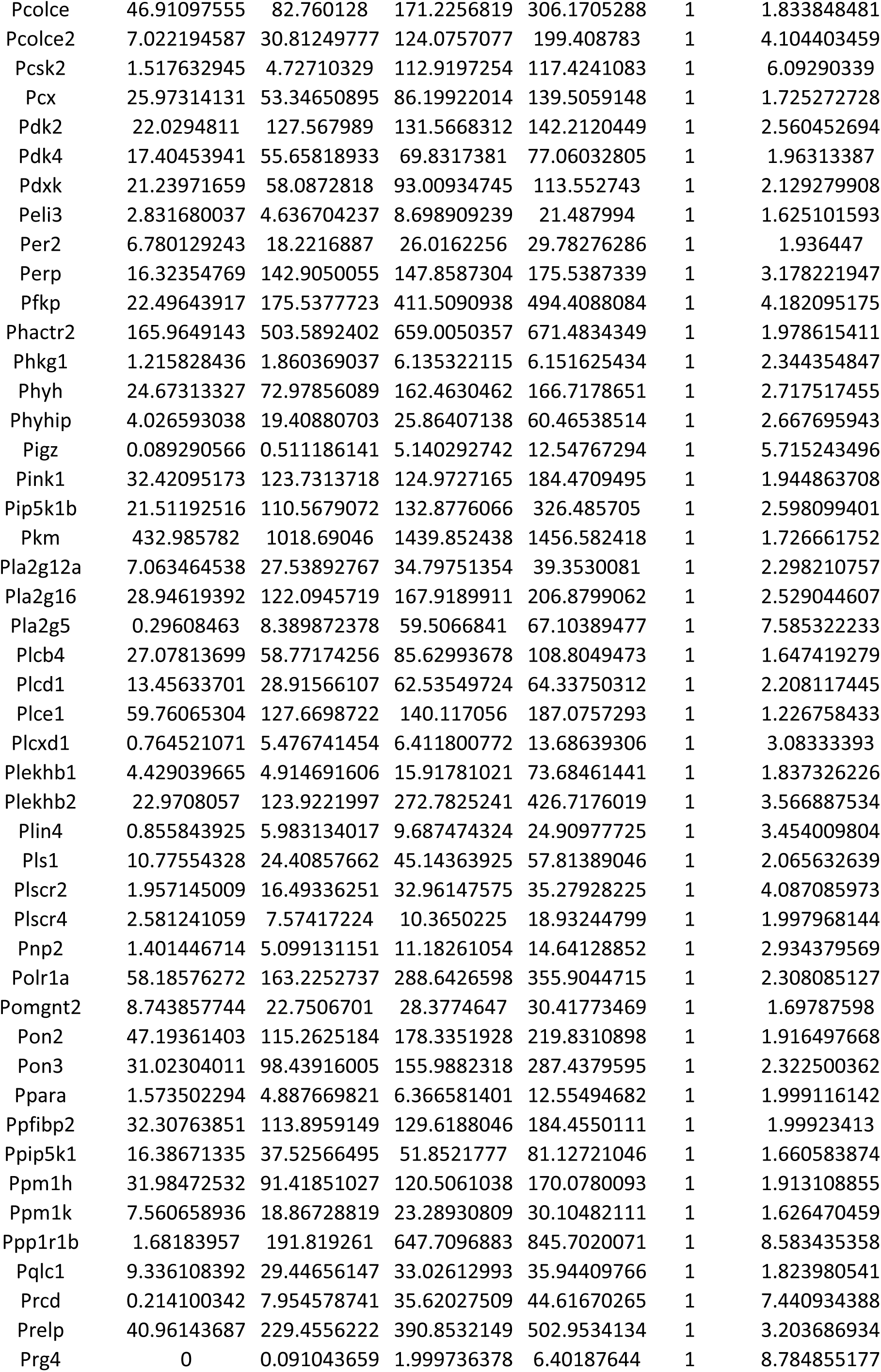
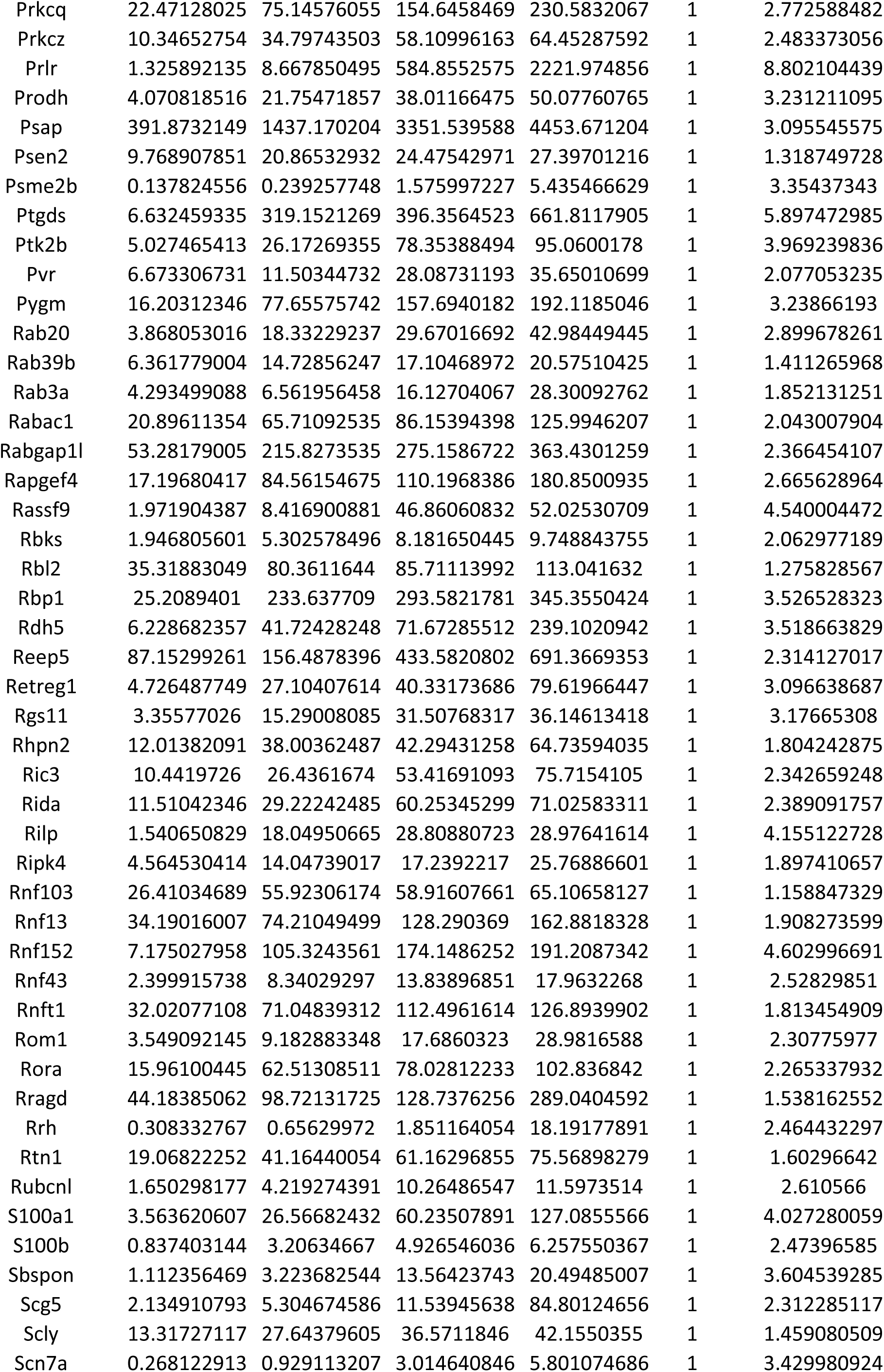
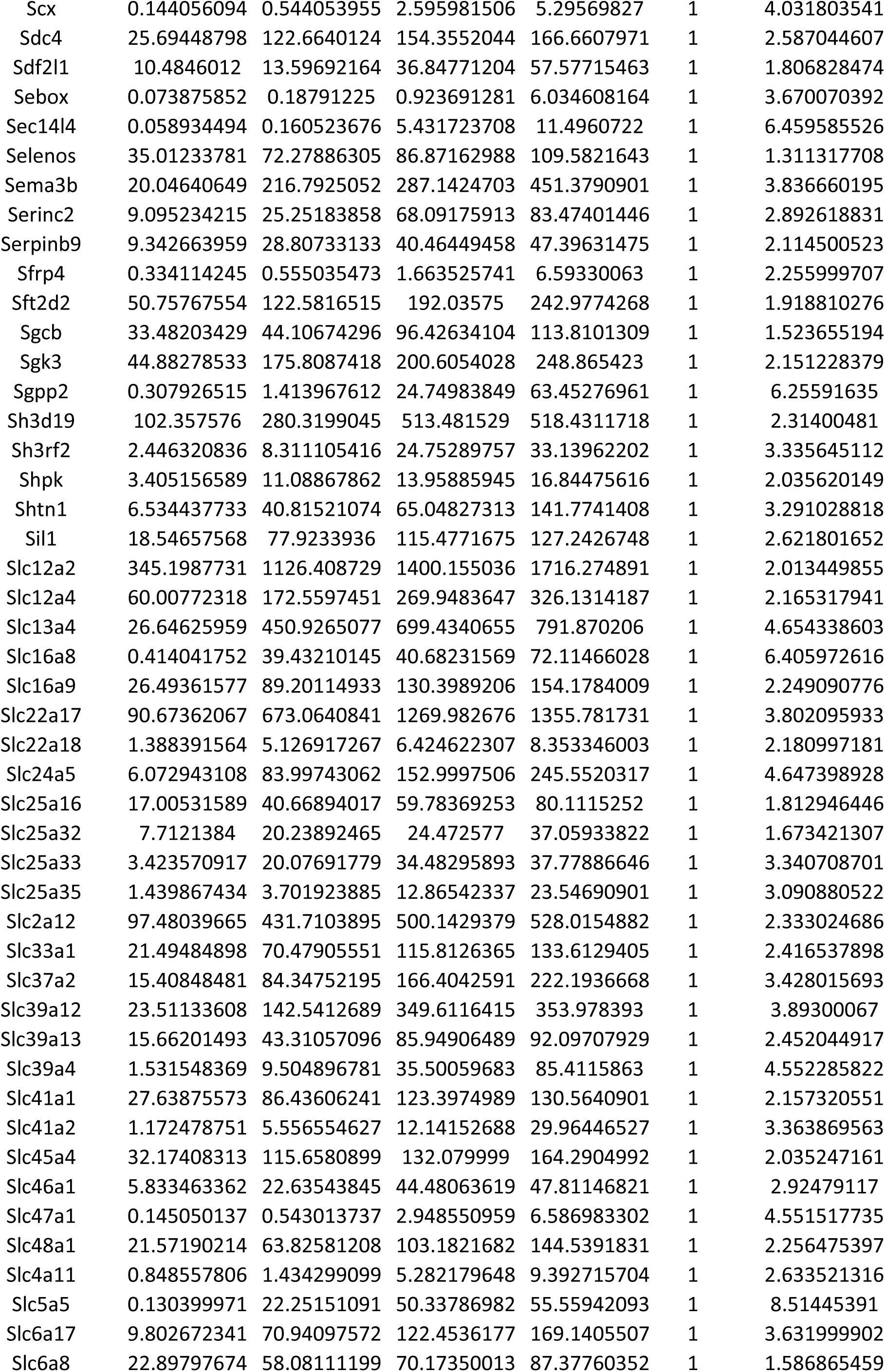
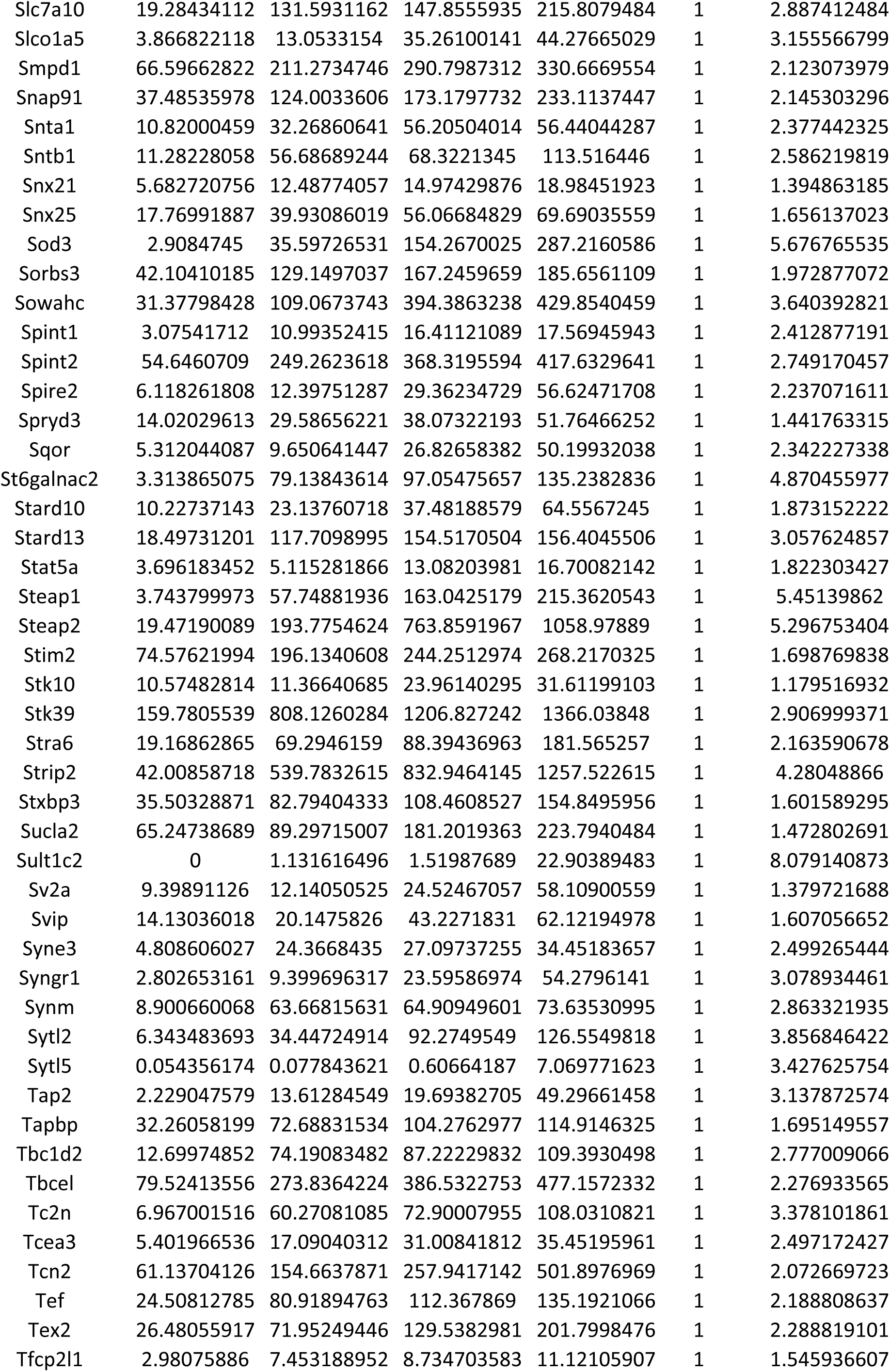
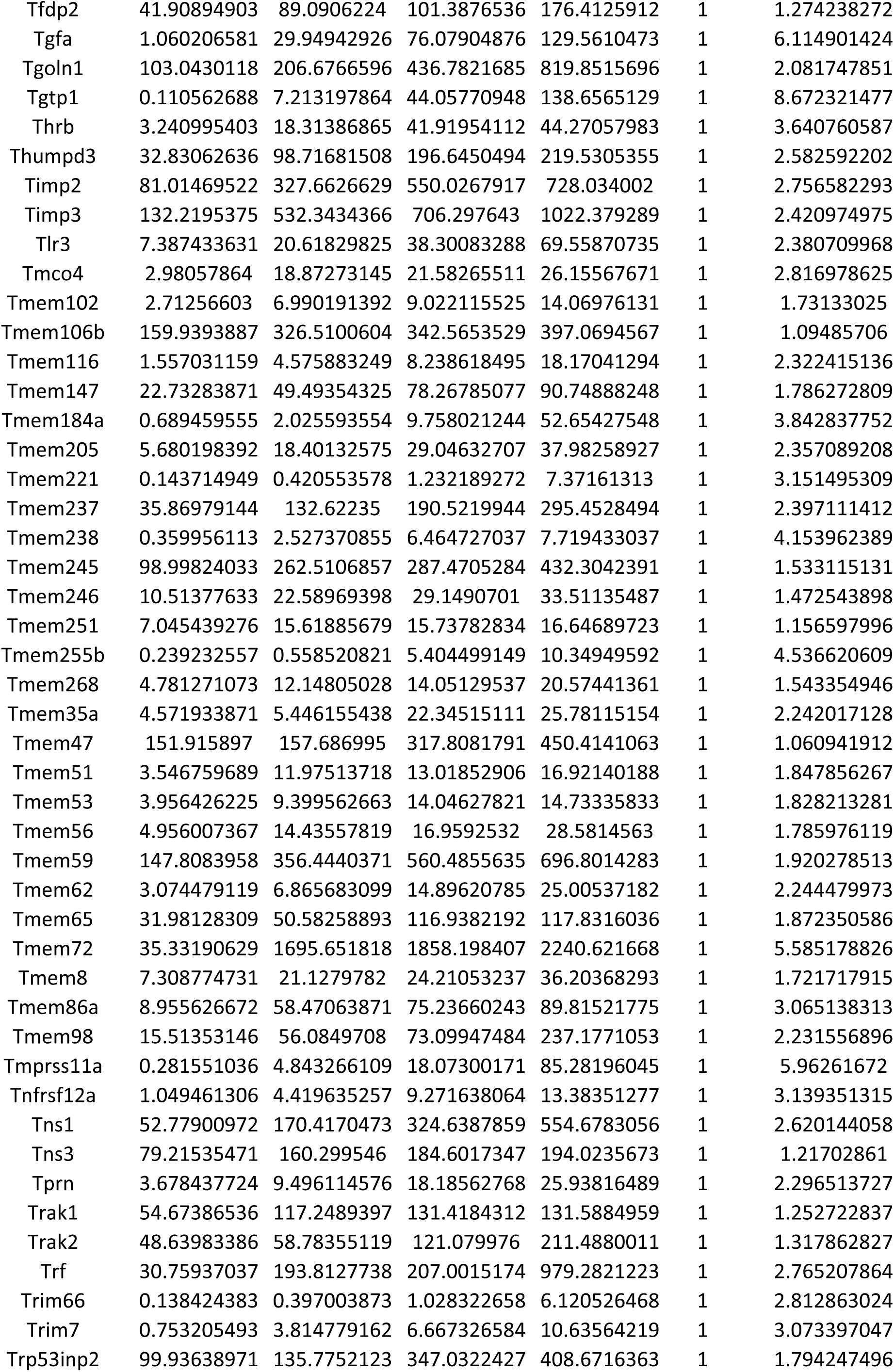
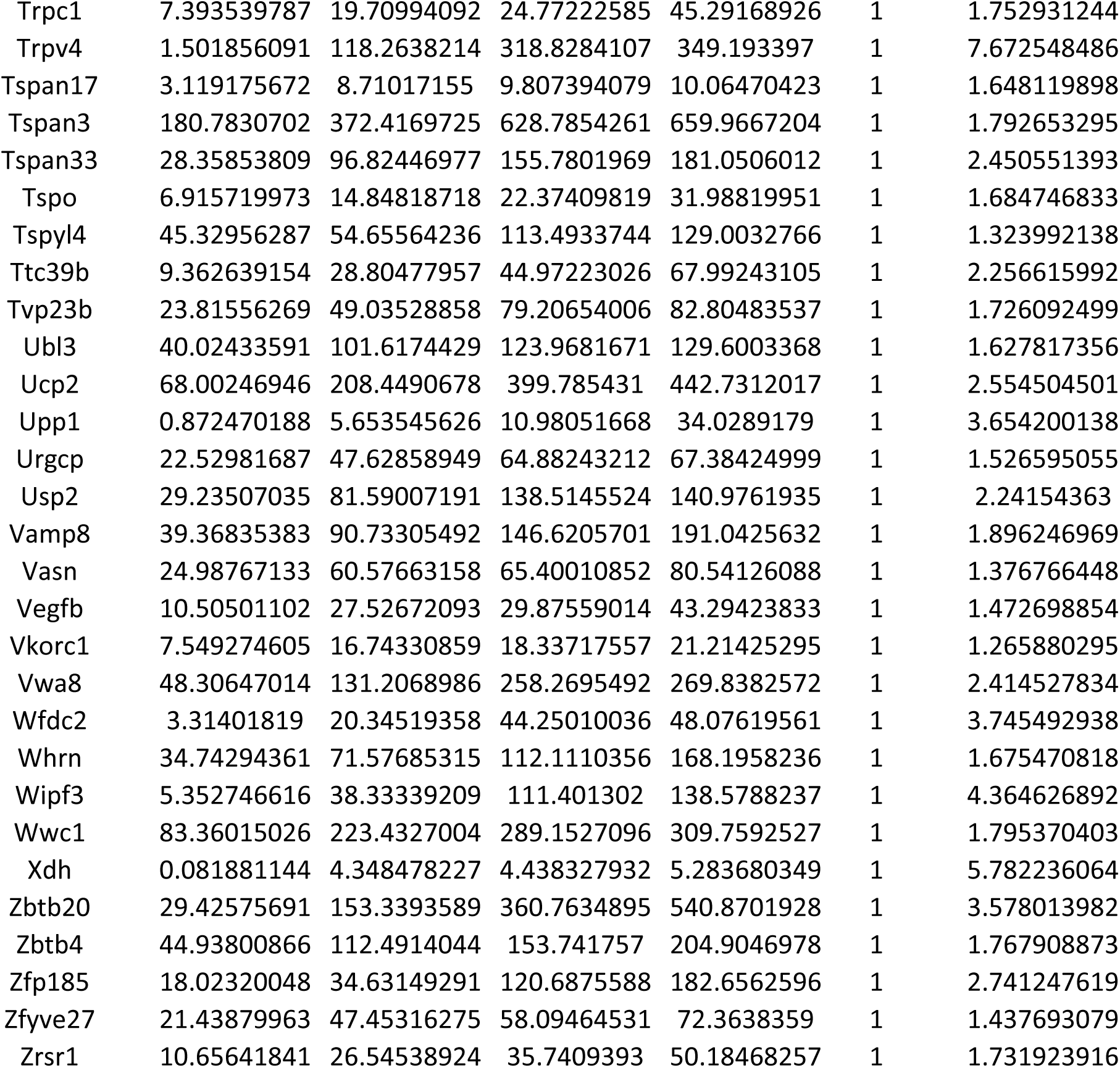
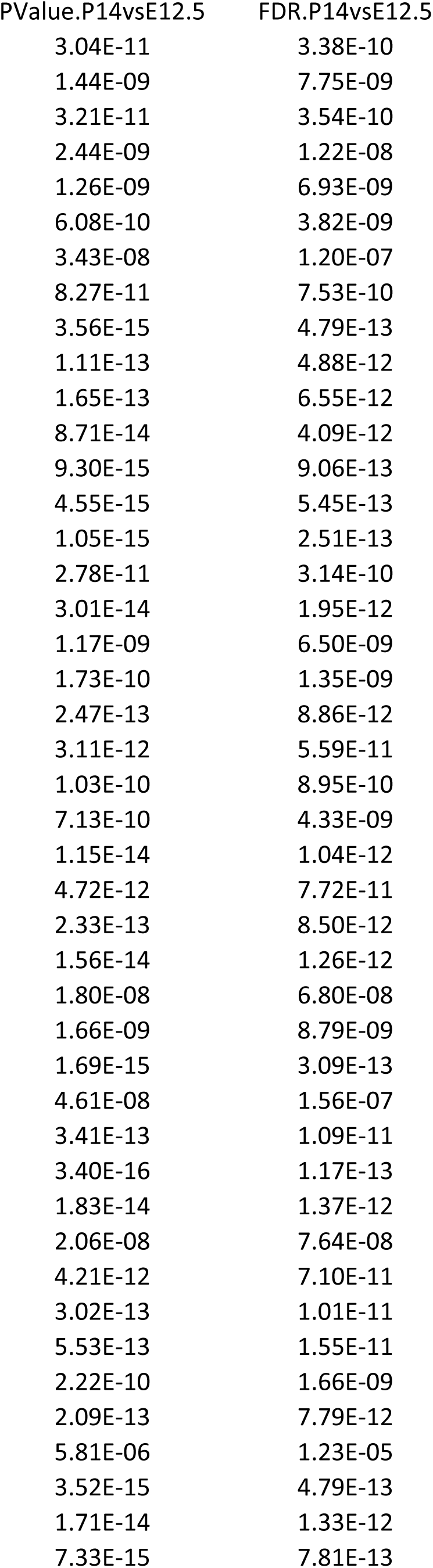
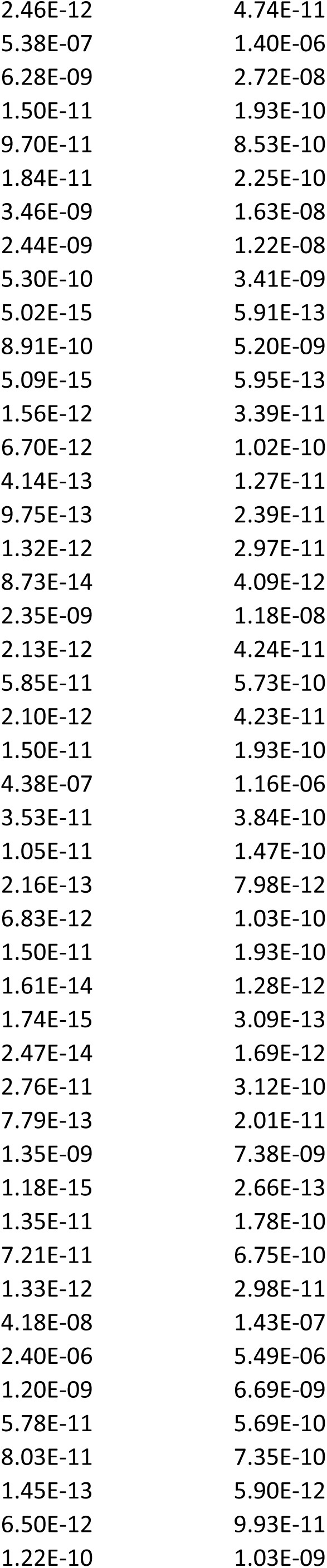
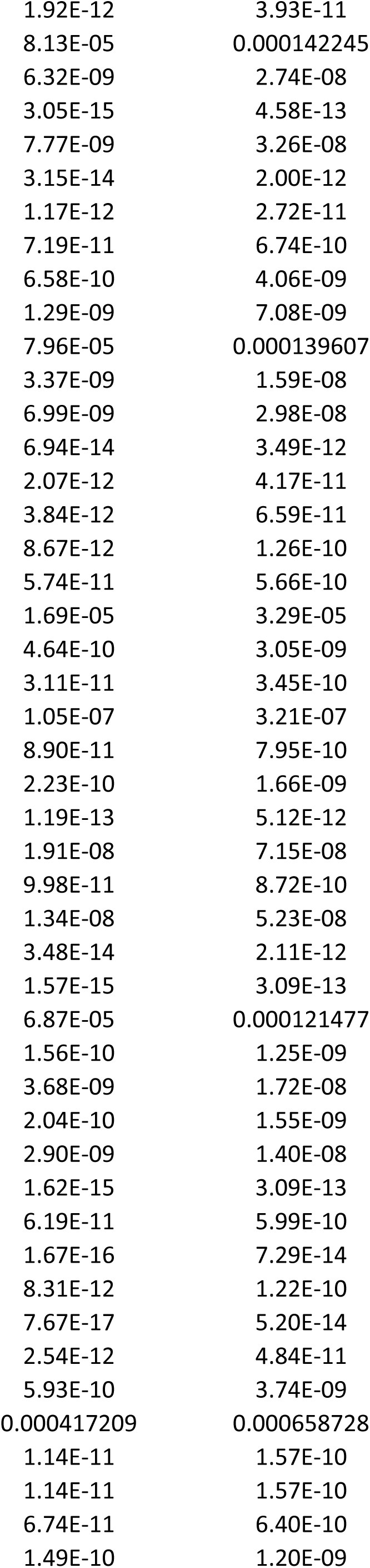
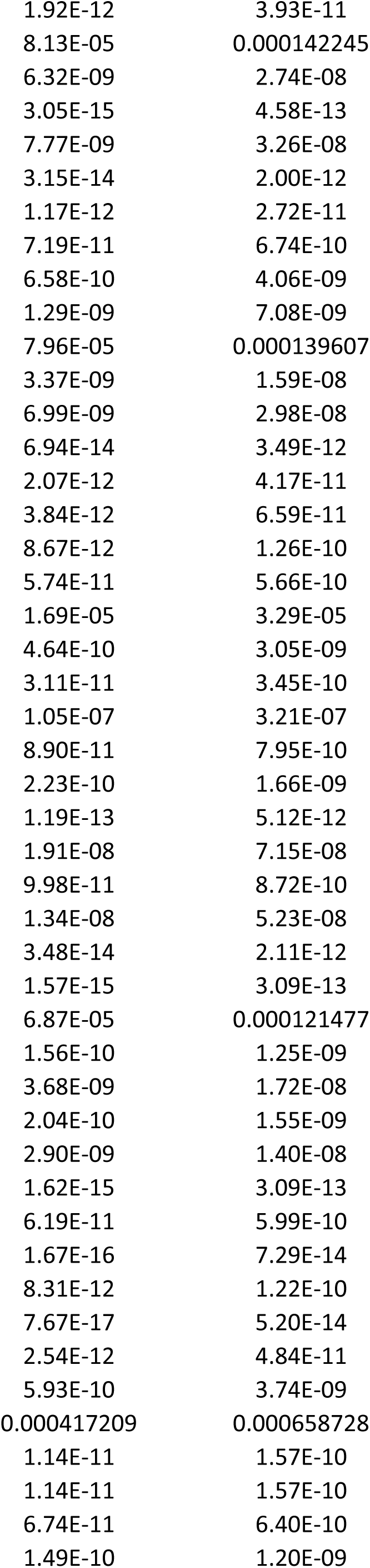
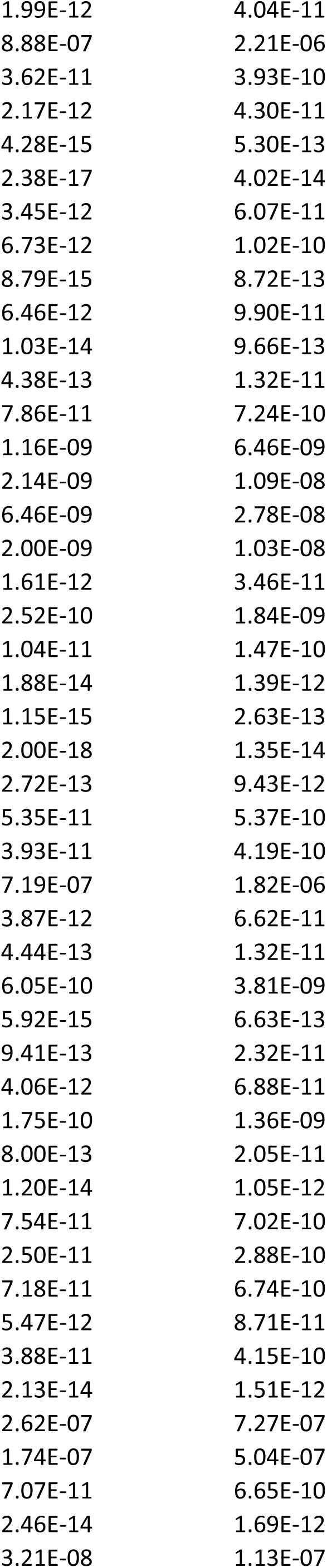
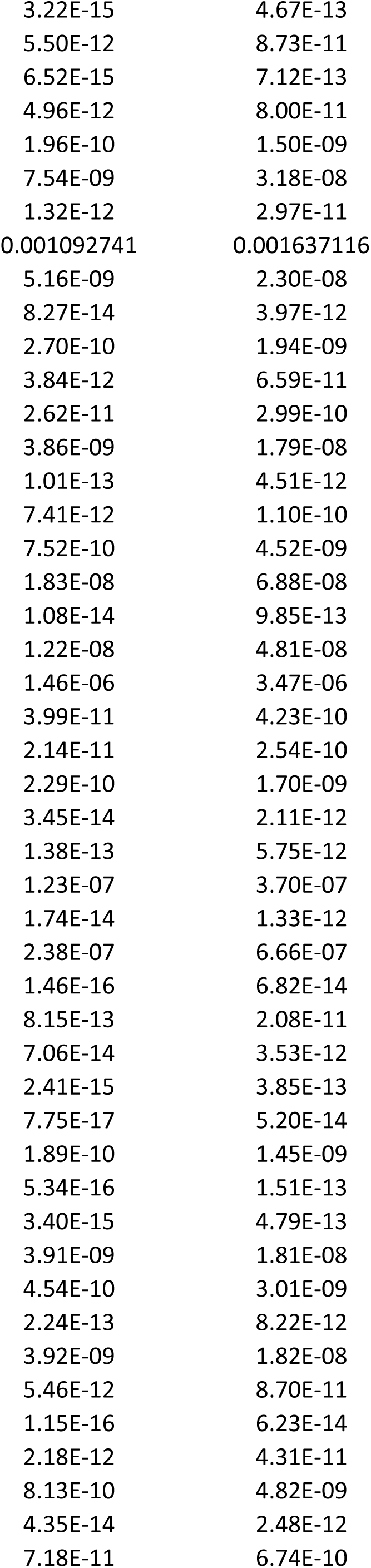
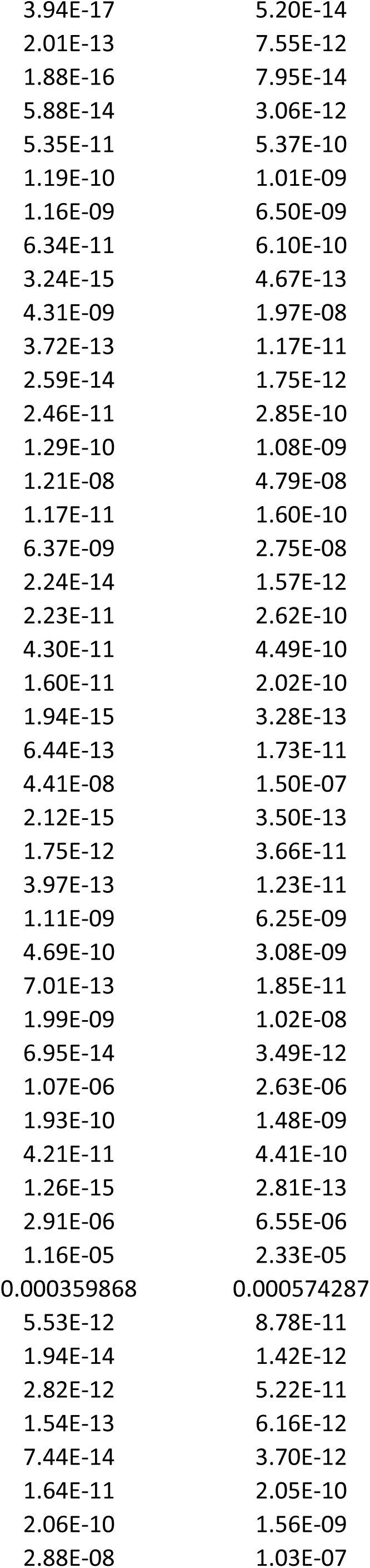
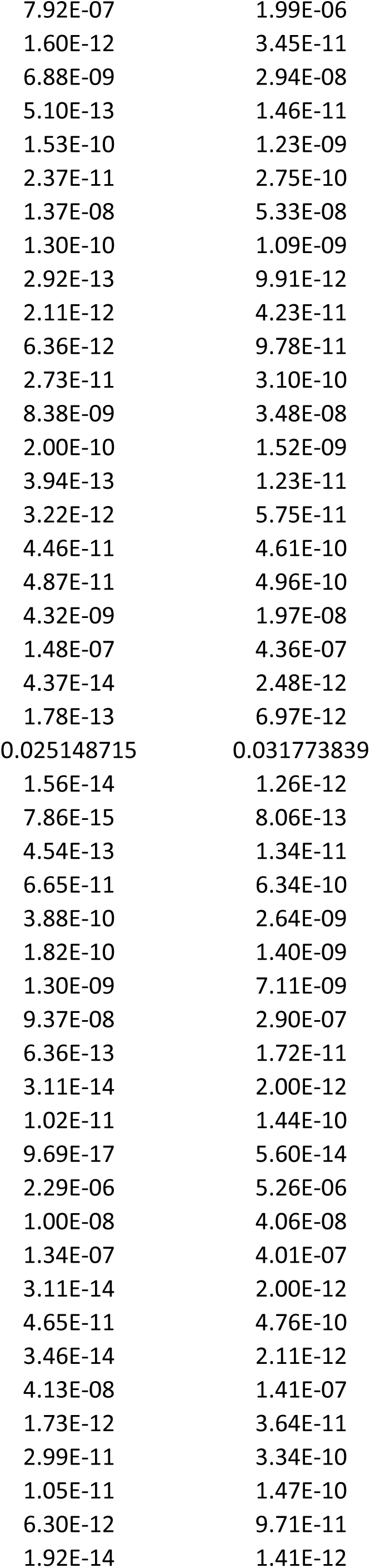
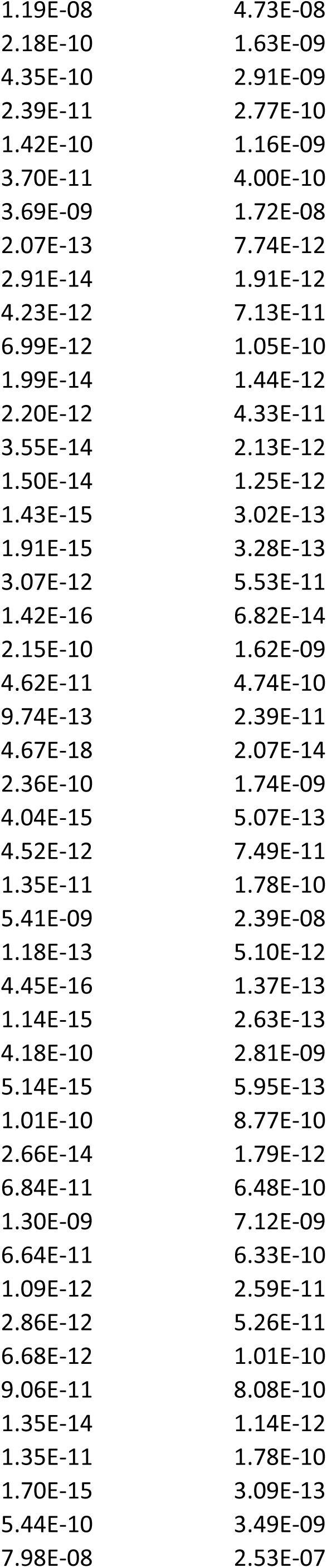
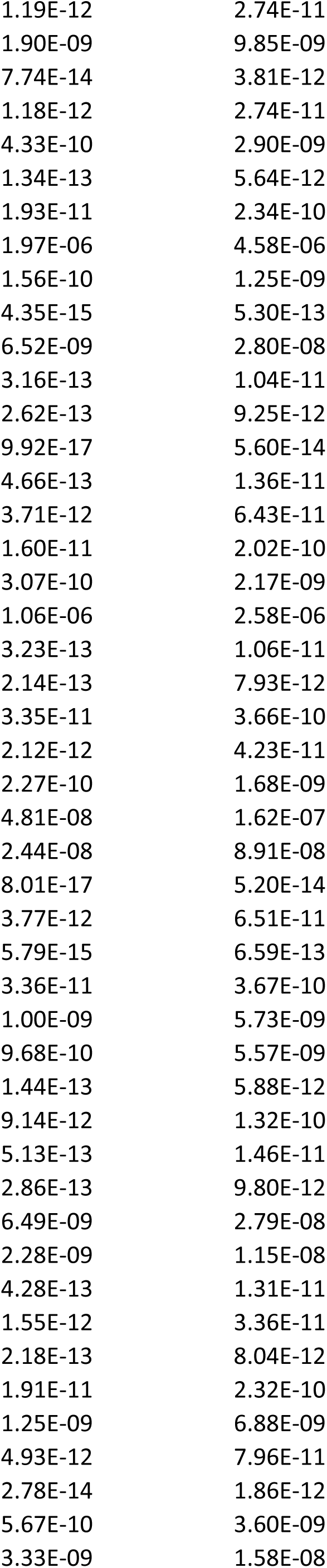
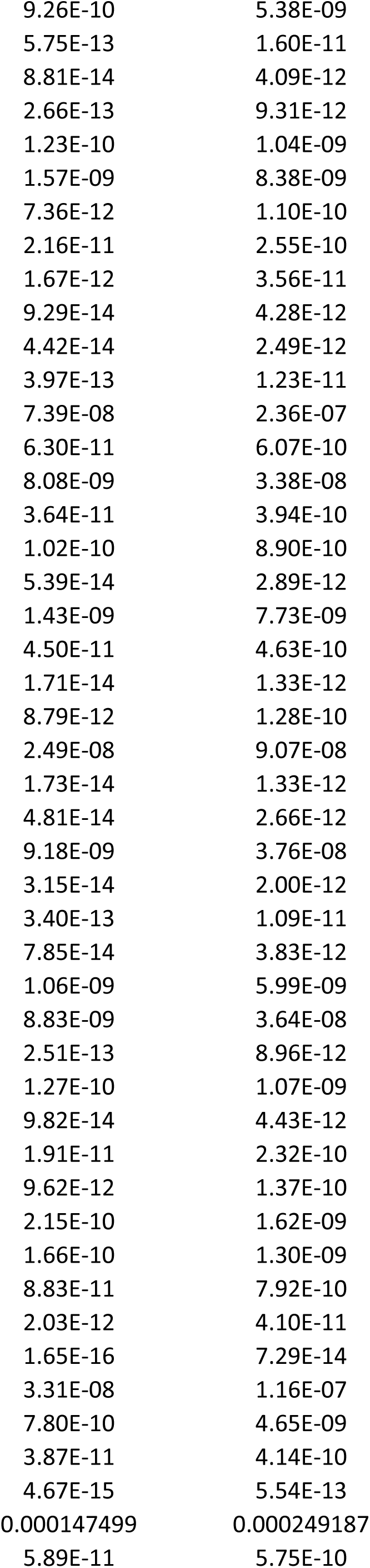
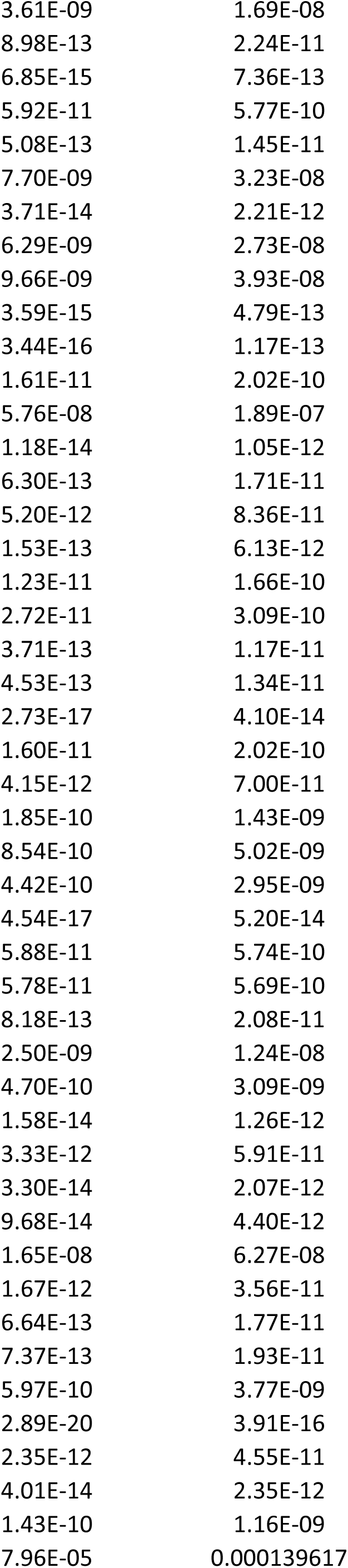
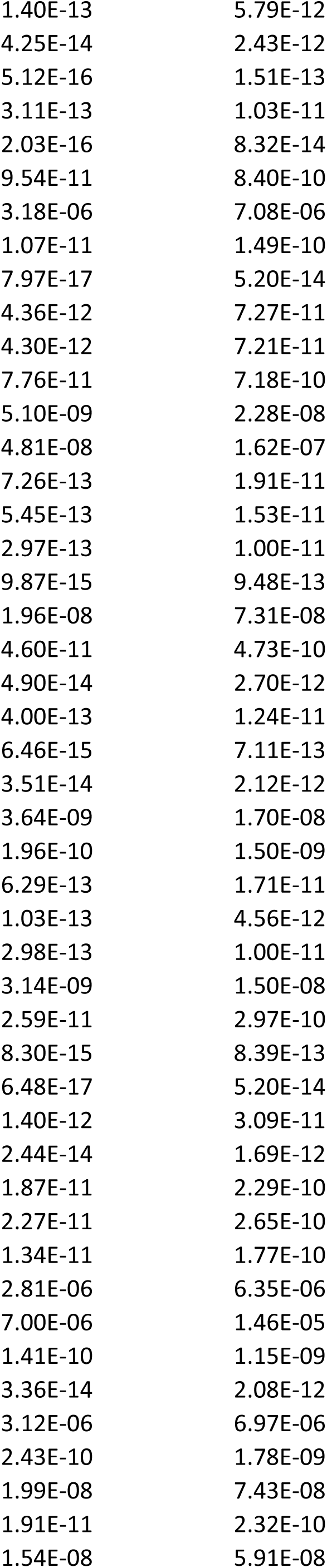
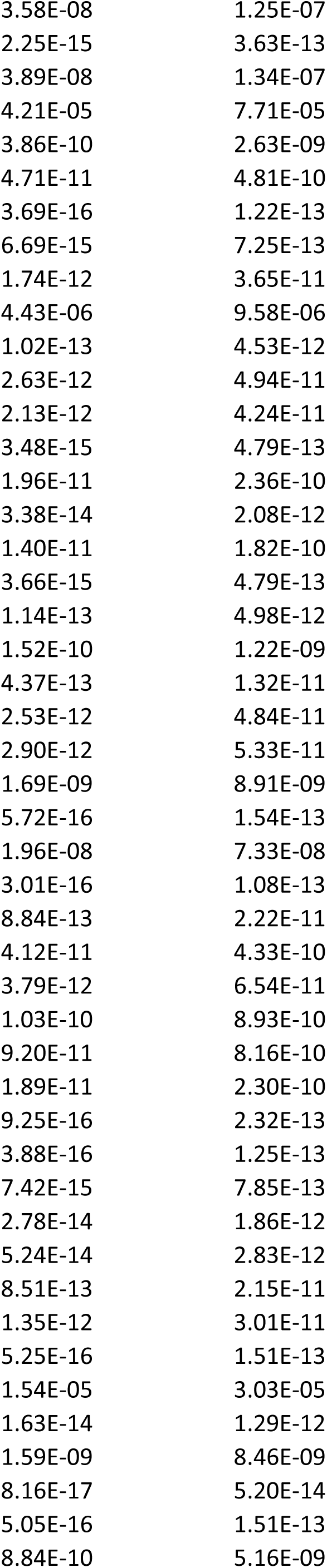
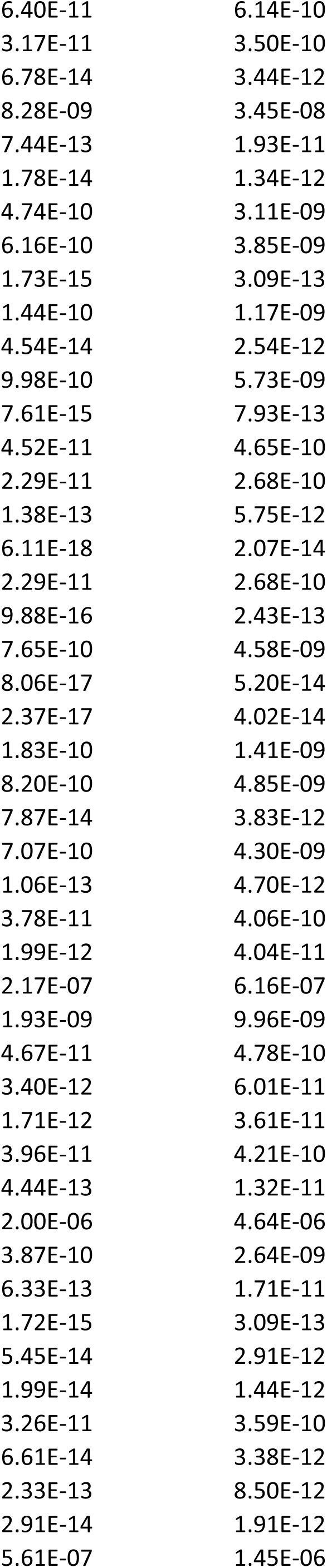
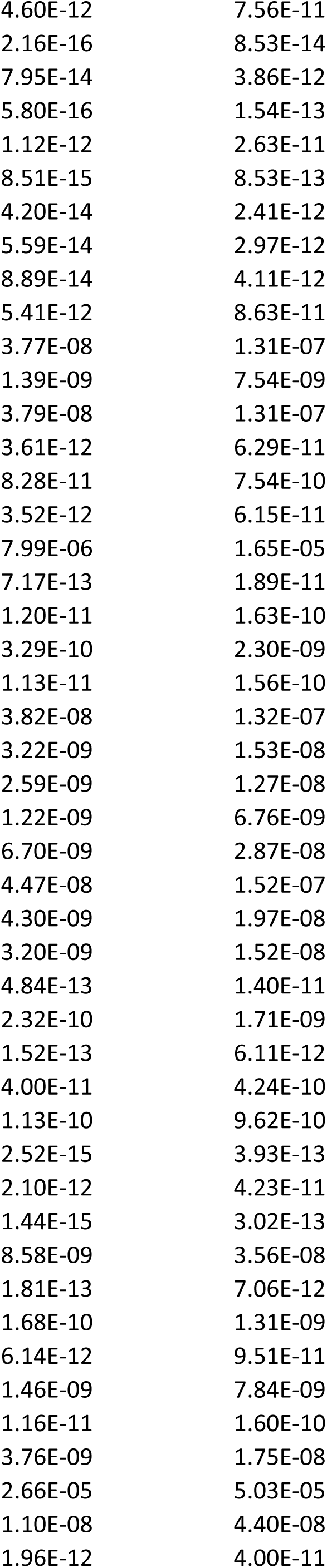
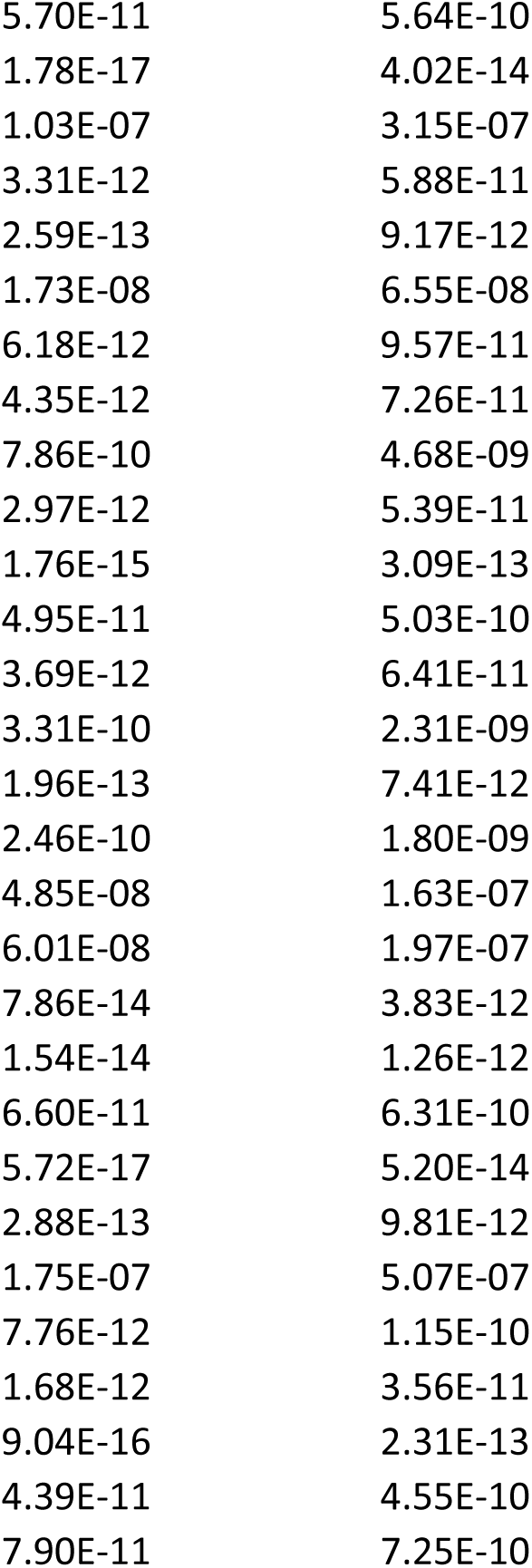
Gene expression during choroid plexus development. Table S2.2 Developmentally upregulated genes during choroid plexus development.

**Table S3.**
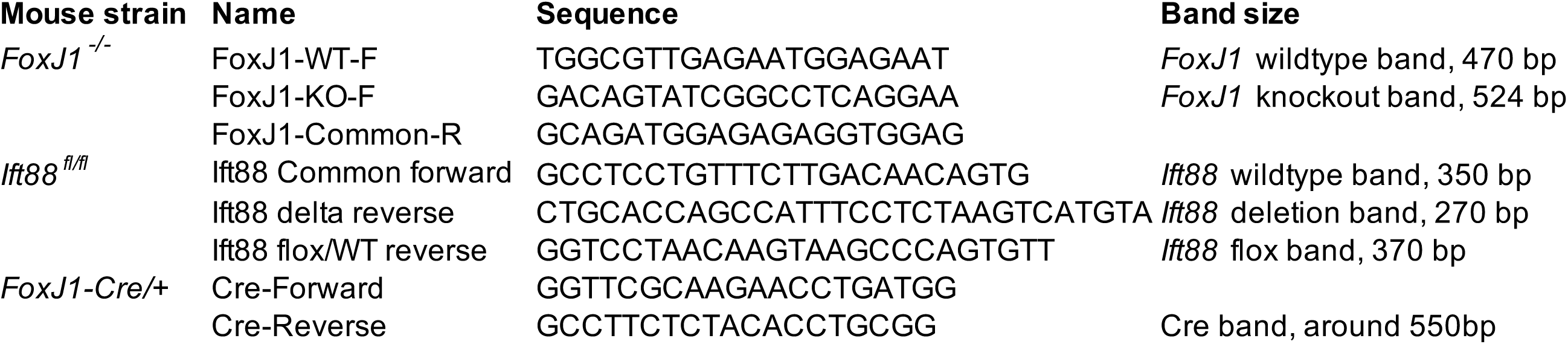
Primer sequences for mouse genotype and qRT-PCR. Table S3.1 Primer sequences for mouse genotype.

**Table S3.**
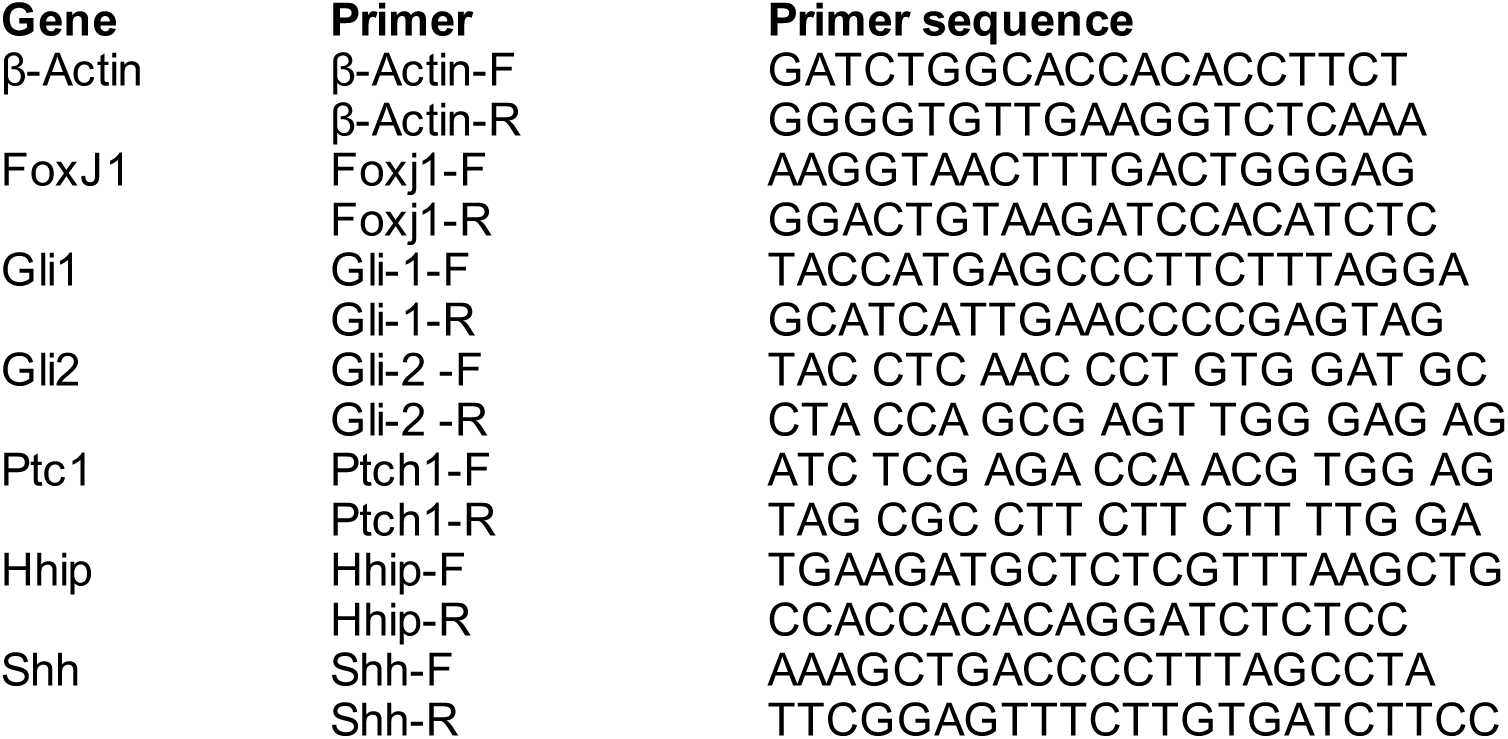
Primer sequences for mouse genotype and qRT-PCR. Table S3.2 Primer sequences for real time PCR analyses.

## Supplementary movie legend

**Supplementary movie S1.**

A 3D reconstruction of P14 mature choroid plexus cilia.

The supplementary movie was uploaded to a Google Drive below: https://drive.google.com/drive/folders/14BDzIF5Ct68rjznDu5kiT1FUXUurLzhA?usp=drive_link

